# Z-REX: Shepherding Reactive Electrophiles to Specific Proteins Expressed either Tissue-Specifically or Ubiquitously, and Recording the Resultant Functional Electrophile-Induced Redox Responses in Larval Fish

**DOI:** 10.1101/2022.10.06.511074

**Authors:** Kuan-Ting Huang, Jesse R. Poganik, Saba Parvez, Sruthi Raja, Brian Miller, Marcus J. C. Long, Joseph R. Fetcho, Yimon Aye

## Abstract

**Summary of the Protocol Extension:** This Protocol Extension describes the adaptation of an existing Nature Protocol detailing the use of T-REX (targetable reactive electrophiles and oxidants)—an on-demand redox targeting toolset in cultured cells. The adaptation described here is for use of REX technologies in live zebrafish embryos (Z-REX). Zebrafish embryos expressing a Halo-tagged protein of interest (POI)—either ubiquitously or tissue-specifically—are treated with a HaloTag-specific small-molecule probe housing a photocaged reactive electrophile (either natural electrophiles or synthetic electrophilic drug-like fragments). The reactive electrophile is then photouncaged at a user-defined time, enabling proximity-assisted electrophile-modification of a POI. Functional and phenotypic ramifications of POI-specific modification can then be monitored, by coupling to standard downstream assays, such as, Click chemistry-based POI-labeling and target-occupancy quantification; immunofluorescence or live imaging; RNA-Seq and qRT-PCR analyses of downstream-transcript modulations. Transient expression of requisite Halo-POI in zebrafish embryos is achieved by mRNA injection. Procedures associated with generation of transgenic zebrafish expressing a tissue-specific Halo-POI are also described. The Z-REX experiments can be completed in <1-week using standard techniques. To successfully execute Z-REX, researchers should have basic skills in fish husbandry, imaging, and pathway analysis. Experience with protein or proteome manipulation is useful. This protocol extension is aimed at helping chemical biologists study precision redox events in a model organism and fish biologists perform redox chemical biology.

## INTRODUCTION

Signaling pathways are complex, exquisitely-orchestrated processes, often fine-tuned for dynamic regulation in response to both beneficial and harmful stimuli that life encounters from early development to death^1^. Our current knowledge of signaling pathways such as kinase cascades, ubiquitination, and transcriptional switches underpins many of our current efforts in prophylaxis, drug discovery, and personalized medicine. Much of our current understanding has come to light from studies in cultured cells. However, it has become apparent that where possible, whole organism studies are more appropriate. This is, at least in part, because non-transformed cells are used in their appropriate tissue and organismal context with their relevant endogenous regulation^2^.

Nature has learned to harness numerous small-molecule messengers that are ‘interpreted’ by proteins, which in turn propagate such directed biological messages for decision making at cellular and organismal levels. Many traditional messengers covalently modify a protein of interest (POI) in an enzyme-catalyzed post-translational modification (PTM) reaction, leading to (a) change(s) in activity, function, locale, stability, etc., of the POI^1^. Beyond small-molecule-based messengers, such as acetate^3^ or phosphate^4^, single-domain proteins such as ubiquitin^5^ constitute among the most well-recognized canonical PTMs. By contrast, reactive small-molecule signaling mediators^6–11^ such as reactive electrophilic and oxidative species (RES and ROS)^10, 11^ orchestrate non-canonical PTMs distinct from canonical enzyme-assisted PTMs. This protocol will focus on a class of RES, namely, endogenously-generated reactive lipid-derived electrophiles (LDEs)^8, 10^, typified by 4-hydroxynonenal (HNE).

In canonical enzyme-assisted PTMs, substoichiometric modifications of a single POI at a specific residue are sufficient to drive functional responses. Likewise, LDE modifications at low occupancy can functionally modulate phenotypic responses at low ligand occupancy^10^. However, the lack of enzyme mediation in LDE-signaling renders the functional consequences of such non-canonical PTMs not directly or precisely tractable using conventional genetics and biochemical tools. Consequently, despite their initial discoveries in the 60’s and long-standing appreciation of LDE’s roles in pathophysiology^8–10^, little is known about how these individual LDE–POI engagement modulates context-specific cell signaling and decision-making.

LDEs such as HNE when introduced from outside the cell or animals, will modify multiple proteins^10, 12–15^. This single fact renders precision deconvolution of protein HNEylation intrinsically difficult. Organismal models of HNE stress are reasonably commonplace and include injured heart^16^, retina^17^, liver^18^, and so on. Doubtless HNE-signaling does happen in such injury models. However, these inherently-stressed systems have undergone significant reprogramming or impaired regulation, and hence are unlikely to reflect *regulated* HNE-signaling that occurs at basal levels to *maintain* fitness. On the other hand, the most general experimental means to model RES- and LDE-signaling involves bulk addition of HNE to cells or whole organisms^8–10, 12, 13, 19^. Although the starting state is defined in this case, the intrinsic reactivity, promiscuity and pleiotropic attributes of these reactive and toxic LDEs, or related electrophilic drugs^20^, following bulk/bolus administration of at supraphysiological concentrations, render on-target phenotypic outputs challenging to parse from off-target behaviors. As these off-target effects accrue with time, and given the high end-point toxicity associated with HNE^10, 21^, indirect off-target effects eventually dominate over beneficial signaling. Thus, negative effects of acute HNE exposure under uncontrolled settings contribute significantly, and in a complex way, to measured outputs^21^. Furthermore, lack of tractability in terms of subcellular distribution, electrophile dosage, precise electrophile chemotype (due to metabolic vulnerability of LDEs^10, 21^), and time of POI-LDE engagement, etc., render understanding of non-canonical PTMs mediated by reactive electrophiles hugely limited. Unfortunately, similar to the injury models above, this ‘bolus dosing’ method remains at best an approximation to physiological signaling in otherwise unperturbed and undamaged live specimens. Bolus dosing likely falls short of accurately representing the nuances associated with *bona fide* context-specific redox-linked signaling pathways^10^. Moreover, bolus approach certainly lacks the ability to interrogate individual protein- and ligand-specific signaling behaviors that is necessary for gaining molecular-level understanding of how LDEs function in cells and organisms, and how this understanding can be effectively leveraged for electrophilic fragment and target design and discovery^22–24^.

### Development of the Protocol

We here describe a unique chemical-biology protocol to interrogate cause and consequences of RES-signaling in live larval fish, mediated through a class of prevalent endogenous LDEs^8, 10^. We refer to this method as “Z-REX” (Zebrafish targeting of Reactive Electrophiles and oXidants, Fig. 1). We have previously disclosed a protocol to deliver a number of natural and preternatural LDEs to specific POIs in cultured cells^25^ (referred to as T-REX). We here focus our Z-REX protocol on the delivery of HNE as a representative LDE. Although they are not discussed here, we have recently shown that related LDEs, such as 4-hydroxydodecenal (HDE), function equally well in Z-REX^26^.

**Figure 1.**
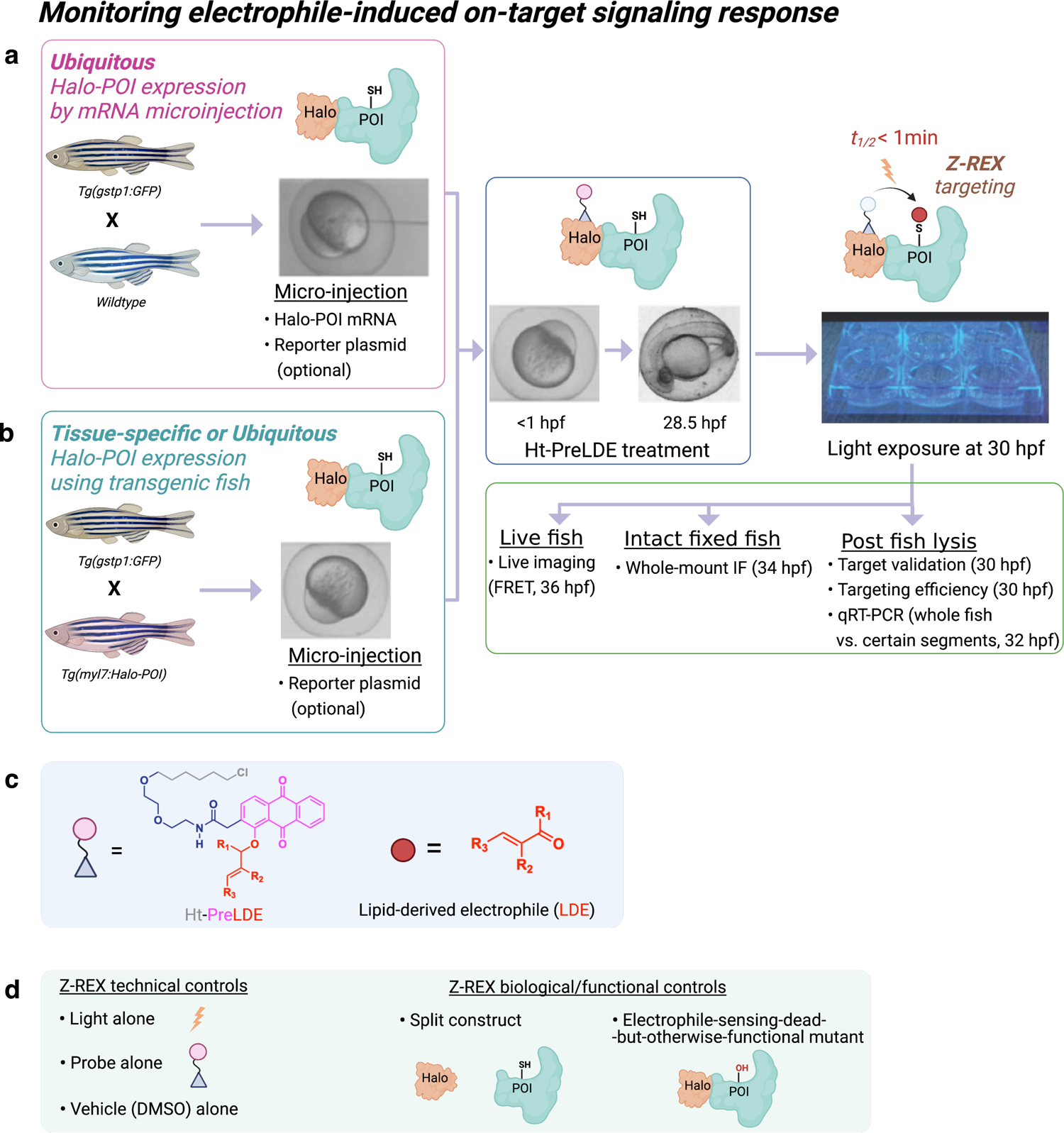
*Workflow for Z-REX*. (**a**-**c**) Zebrafish embryos are injected either at the 1–4 cell stage for injection of mRNA encoding a Halo-tagged protein of interest (POI), or at the 1 cell stage for co-injecting reporter plasmid and mRNA encoding a Halo-tagged POI (**a**). Alternatively, embryos expressing desired Halo-POI fusion protein either ubiquitously or in certain tissues, can be obtained by crossing appropriate transgenic fish lines (**b**). Embryos are then treated with the HaloTag-targetable small-molecule ligand (**grey)**, the photocaged precursor (pink) to lipid-derived electrophile (**red**) (**c**, Ht-PreLDE). After 24 h, embryos are washed 3 times, then exposed to a hand-held lamp (365 nm, 3 mW/cm^2^) for 2–5 min, which enables rapid liberation (*t*_1/2_ < 1 min) of a designated lipid-derived electrophile (**c**, LDE). Should the POI be a kinetically-privileged electrophile sensor, the liberated LDE rapidly reacts with the POI (top-right), prior to LDE diffusion. The extent of LDE occupancy on target POI, downstream signaling, and indicated functional changes triggered as a result of LDE modification of a specific POI, can then be assessed (middle-right). (**d**) Relevant sets of Z-REX technical and functional and biological controls should be implemented to help rule out potential artificial outcomes: by assessing effects (if any) due to light alone, probe alone, or DMSO alone treatment (left panel), or by replicating Z-REX conditions using the split construct or electrophile-sensing-defective POI (right panel). These controls should not give the behavioral or signaling changes observed under Z-REX conditions.

We have taken a different approach to model endogenous physiologic RES signaling: this setup, broadly termed REX technologies, is built on ‘precision localized electrophile delivery’ concept that mimics how these natural LDEs are locally produced *in vivo* with spatiotemporal precision^27^. REX technologies as a whole have not only enabled us to identify novel LDE-sensor proteins^14, 15, 25, 28–30^ but also to study POI-LDE-modification-dependent redox signaling and downstream phenotypic behaviors^15, 26, 27, 30–32^. These technologies have also paved novel avenues toward precision medicine design and development^22–24^. Our inaugural method, under the auspices of precision localized electrophile delivery paradigm, termed T-REX, is applied in live cells^25, 33^, and involves the use of POI as a Halo-fusion. The Halo domain reacts irreversibly and in a 1:1 stoichiometry with an otherwise inert chloroalkane-functionalized photocaged precursor to LDEs. Upon light illumination, T-REX enables proximity-directed covalent labeling of the POI with an LDE, provided the POI reacts with the LDE rapidly before the latter diffuses away^21^. T-REX is an approximation of the true signaling state. However, the protocols we have established allow us to scrutinize and mechanistically interrogate a variety of functionally-distinct LDEylated POIs for the first time in a number of complex redox-regulated pathways^14, 26, 27^. By identifying kinetically-privileged electrophile-sensor proteins, typically via protein-cysteines^21^, T-REX is ultimately primed for pinpointing optimal targets and functional residues for covalent drug design, discovery of novel drug mechanisms, and understanding the ramifications of disease-specific cysteine mutations^23, 24^.

To expand the remit of REX technologies, we recently established Z-REX procedures to deliver HNE (and related natural or preternatural LDEs) specifically to target POIs in live zebrafish^15, 26, 30^. This zebrafish platform, Z-REX, allows study of specific signaling pathways in live zebrafish. Zebrafish are genetically-tractable vertebrate organisms that are transparent during development and ideal for opto-chemical and opto-genetic techniques^34^. They possess many organs analogous to those found in humans and develop quickly. Indeed, one of the key conclusions of our protocol is that our regimen affords instantaneous delivery and rapid dissemination of protein-specific ‘on-target’ electrophile modification in fish. By contrast, the corresponding bolus-dosing methods can be toxic, inefficiently label the “intended target protein”, and give responses that are no better than those observed for T-REX^14, 25, 26, 29, 31–33^.

This Z-REX procedure—including plasmid creation, mRNA expression, and validation of POI-labeling and POI-electrophile-modification-dependent pathway signaling—has been demonstrated for several different ectopic human proteins expressed in fish, including Akt3^30^, Keap1^26^, and Ube2V2^15^; zebrafish orthologs have also been shown to be electrophile sensitive in cultured cells using T-REX. Data emerging in our laboratory show that REX-technologies in general are applicable to most electrophile-sensing proteins^27^. Although the protocol is optimized for zebrafish (*Danio rerio*), given that the protocol was readily transposed from cultured mammalian cells, it will likely work well on other genetically-tractable fish, such as medaka (*Oryzias latipes*) and mumichong (*Fundulus heteroclitus*)^35^. The broad applicability of the T- and Z-REX method is likely ascribable to the pseudo-intramolecular mechanism of LDE-delivery^21, 27, 36^, which means that LDE-modification is POI-expression-level- and environment-independent, but POI-specific. We have previously disclosed evidence for this mechanism in cells^25, 27, 33^. We recently provided evidence for the mechanism of Z-REX^26^, which is also discussed in this protocol.

In addition, we have shown that standard assays that are commonly used in fish and some that we have repurposed are fully compatible with Z-REX (Fig. 1). Levels of ectopic Halo-POI are assessed by IF (Figs. 2c– d, 3a–b, and Extended Data Fig. 1a–d) or western blot (Fig. 3c–d). Labeling is assessed by affinity capture post click chemistry-mediated tagging of electrophilic-modified proteins (Fig. 3c–d). Downstream signaling is assessed by: transgenic reporter fish available from the NBRP (*vide supra*) for our transcriptional readout of interest (Figs. 4a–c, Fig. 5 and Extended Data Figs. 2a, 3 and 4); quantitative reverse transcription PCR (qRT-PCR) (Fig. 4d, endogenous loci); and also using fluorescence reporter assay (Fig. 6).

**Figure 2.**
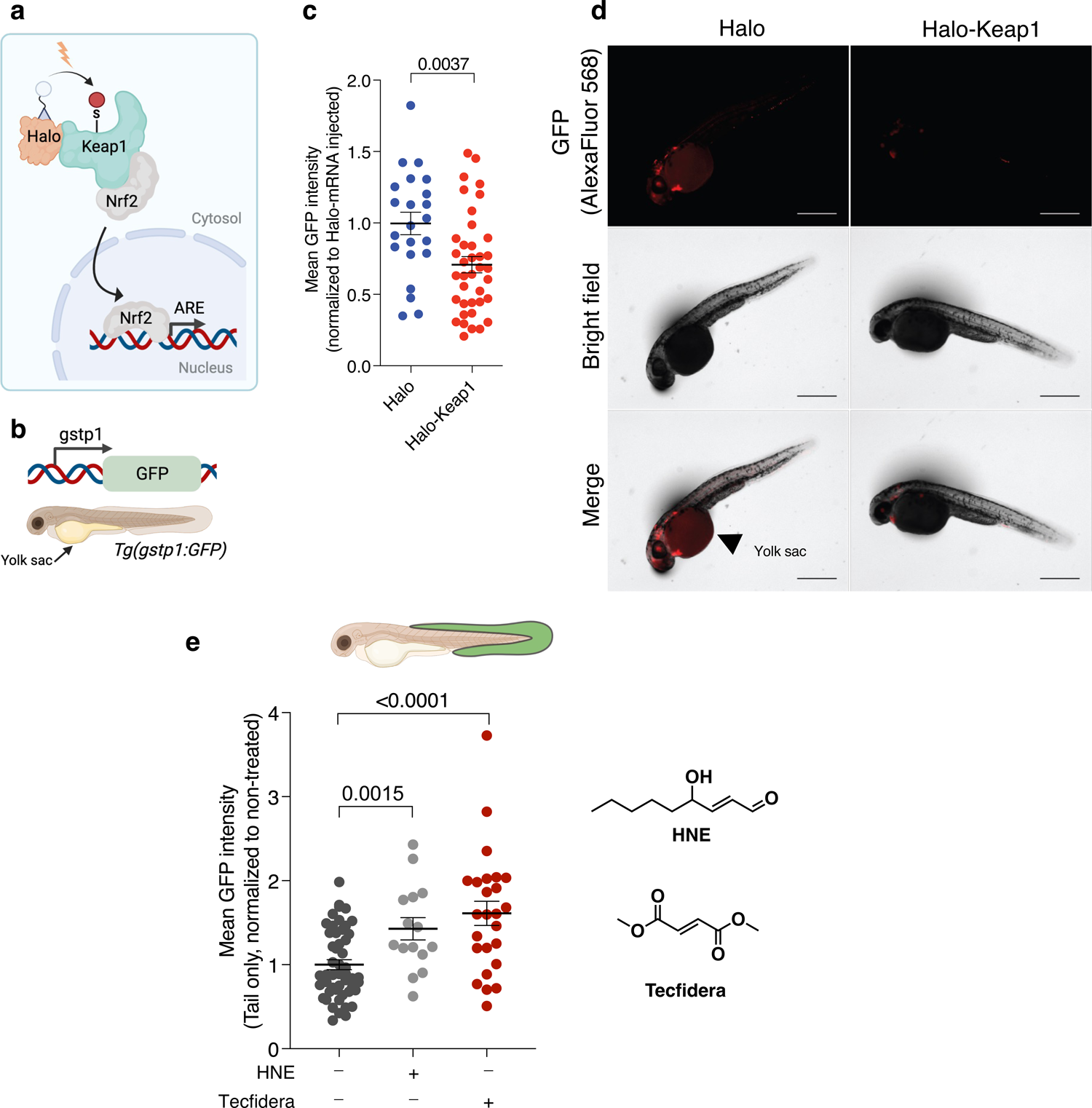
*Validation of POI expression and POI-selective targeting in zebrafish larvae*. The established electrophile-sensor POI, Keap1, and the pathway it regulates, Keap1–Nrf2–AR^25^, was used as a representative example. (**a**) Schematic of the light-driven precision targeting of native lipid-derived electrophiles such as HNE to ectopic Halo-Keap1 elicits upregulation of antioxidant response (AR) element (ARE)-driven genes. (**b**) Schematic of *Tg(gstp1:GFP)* reporter fish available from NBRP (see discussion). (**c**) *Tg(gstp1:GFP)* were injected with mRNA coding for either only Halo or Halo-Keap1. After 34 hpf, AR was assessed by IF imaging for GFP. Halo: n=22, SEM=0.0788; Halo-Keap1: n=38, SEM=0.0567. (**d**) Representative images from (c). Age: 34 hpf. Scale bar, 500 µm in all images. (**e**) *Tg(gstp1:GFP)* were injected with mRNA coding for Halo-Keap1. At 30 hpf, embryos were treated with DMSO, HNE (25 µM) or Tecfidera (25 µM) for 4 h. Then the extent of AR upregulation specifically in the tail was assessed by IF imaging for GFP (age: 34 hpf). Non-treated: n=50, SEM=0.0582; HNE-treated: n=15, SEM=0.1325; Tecfidera-treated: n=25, SEM=0.1442. ***Inset*:** chemical structure of HNE (a well-studied natural bioactive reactive lipid-derived electrophile) and Tecfidera (an approved multiple sclerosis drug). P values were calculated with two-tailed unpaired Student’s t test.

**Figure 3.**
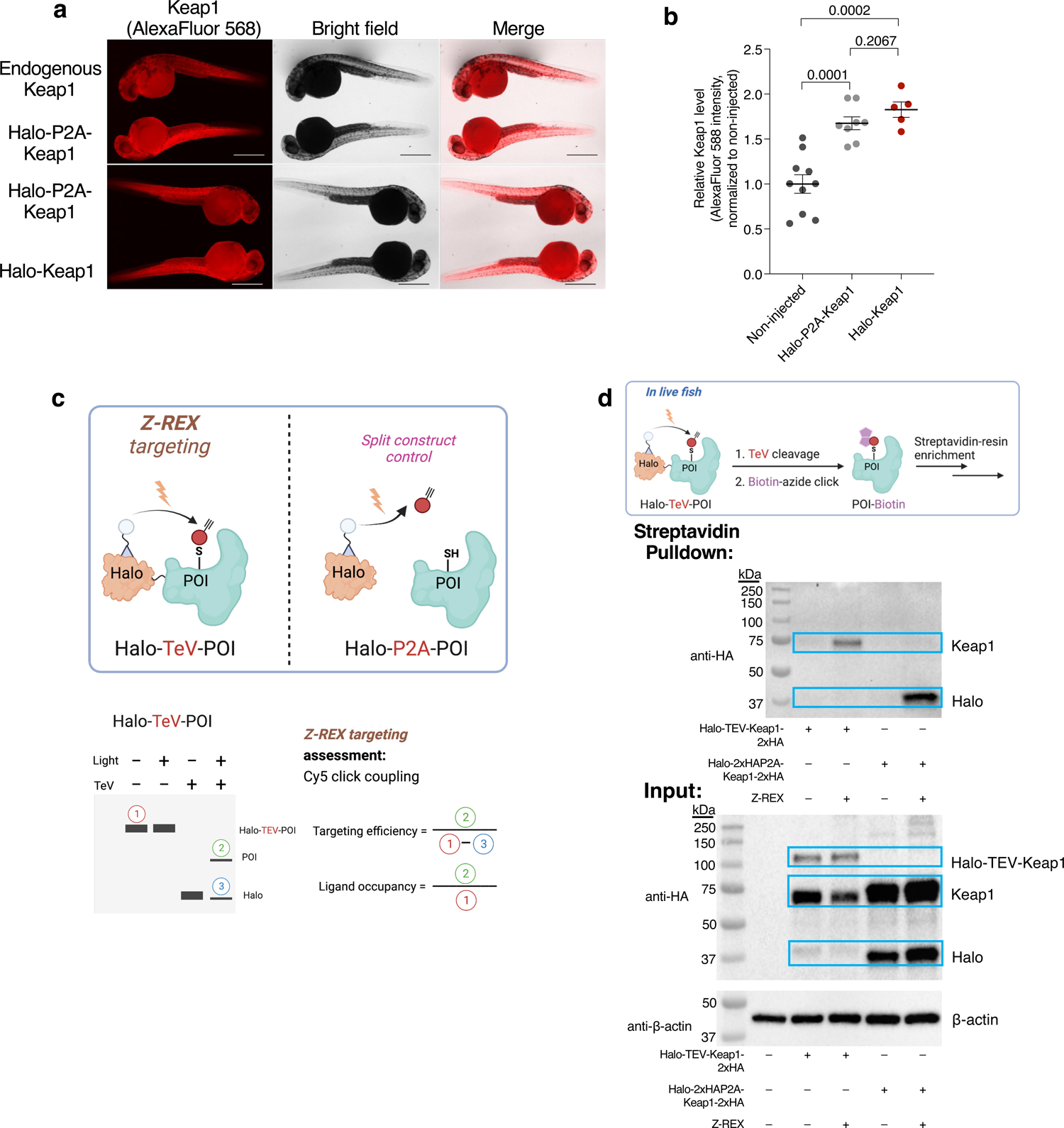
Halo-P2A-POI serves as an ideal negative biological control in Z-REX that proceeds via a quasi-intramolecular mode of POI-specific electrophile modification in vivo. In this example, the established electrophile-sensor Keap1^25^ was used as a representative POI. (**a**) Expression of POI (Keap1 as an example in this case) in non-injected fish and fish injected with the indicated mRNA, was analyzed by IF, by probing with anti-Keap1 antibody (this antibody can detect human Keap1 and both isoforms of zebrafish Keap1) (age: 34 hpf). For negative control (fish not treated with primary antibody), see Extended Data Fig. 1a. Scale bar, 500 µm in all images. (**b**) Quantitation of Keap1 signals (background subtraction by sample treated with secondary antibody only, but not primary antibody) in the indicated sets. **Age: 34 hpf.** Non injected: n=10, SEM=0.1031; Halo-P2A-Keap1 mRNA-injected: n=8, SEM=0.0712; Halo-Keap1 mRNA-injected: n=6, SEM=0.0856. P values were calculated with two-tailed unpaired Student’s t test. (**c**) **Top:** Illustration of the two protein constructs used in this protocol. The Z-REX POI-electrophile modification only happens when the POI is fused to Halo (Halo-TeV-POI), but not when Halo and POI are split (Halo-P2A-POI). (Also see Fig. 1, bottom panel, Z-REX biological or functional control). ***Bottom schematic gel or blot:*** Through Cy5 Click assay (fluorescence gel), the targeting efficiency and ligand (electrophile) occupancy can be calculated. (**d**) WT embryos were injected with mRNA encoding either Halo-(TEV)-Keap1-2XHA or Halo-2XHA-P2A-(TEV)-Keap1-2XHA. Embryos were then treated with Ht-PreHNE (+) or DMSO (–) only. After 30 hpf, all embryos (samples from both + and – lanes) were exposed to light, then immediately harvested. After embryo lysis, the lysate was treated with TEV protease before biotin pulldown assay, to selectively enrich the proteins that had been modified by HNE(alkyne-functionalized). [**Note:** (1) The Halo from Halo-P2A-Keap1 was tagged with HA, but not the Halo from Halo-TEV-Keap1: see labels within the figure for a more detailed description of constructs; (2) the photo-uncaging process was not fully completed, which resulted in the enriched Halo band in the last lane in the top blot].

**Figure 4.**
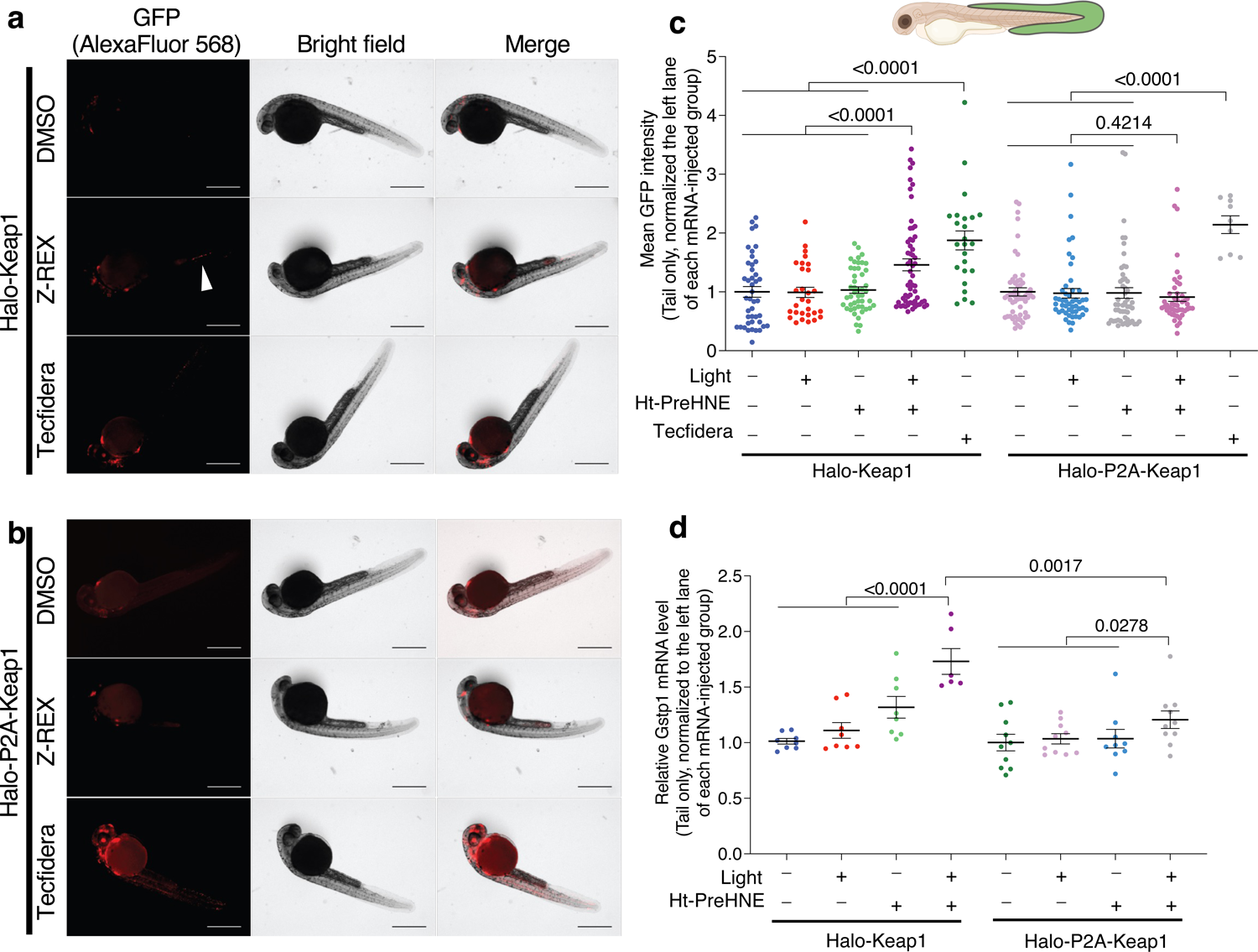
The simultaneous use of genome-encoded fluorescent reporter fish and qRT-PCR method allows an independent validation of pathway activation. In this example, the use of Tg(gstp1:GFP) reporter fish and qRT-PCR, on the established electrophile-sensing Keap1–Nrf2–AR pathway^25^ shows that Z-REX upregulates AR via targeted mechanism that requires association of HaloTag and Keap1. (**a**) Representative images of Tg(gstp1:GFP) fish injected with Halo-(TEV)-Keap1-2xHA exposed to either DMSO, Z-REX (4 h incubation post light exposure), or Tecfidera (25 µM, 4 h treatment). The GFP expression (reporting AR upregulation) induced by Z-REX in tail is indicated with an arrow. Age: 34 hpf. (**b**) Identical to (a) except Halo-2xHA-P2A-Keap1-2xHA construct was used in place of Halo-(TEV)-Keap1-2xHA. Age: 34 hpf. (**c**) Quantification of mean GFP intensity of the reporter fish in (a) and (b). Age: 34 hpf. Halo-Keap1 mRNA-injected (from left to right): n=43, SEM=0.0898; n=29, SEM=0.0872; n=47, SEM=0.0547; n=58, SEM=0.1010; n=24, SEM=0.1613. Halo-P2A-Keap1 mRNA-injected (from left to right): n=55, SEM=0.0689; n=49, SEM=0.0805; n=52, SEM=0.0905; n=47, SEM=0.0721; n=9, SEM=0.1509. (**d**) Similar experimental design to (c), except mRNA was extracted 2-h post light exposure (age: 32 hpf) and relative abundance of *gstp1* (an endogenous downstream gene driven by the conserved Keap1–Nrf2–AR pathway) relative to β-actin (house-keeping gene) was assessed by qRT-PCR. Halo-Keap1 mRNA-injected (from left to right): n=8, SEM=0.0257; n=8, SEM=0.0706; n=8, SEM=0.0975; n=6, SEM=0.1158. Halo-P2A-Keap1 mRNA-injected (from left to right): n=10, SEM=0.0749; n=10, SEM=0.0456; n=9, SEM=0.0836; n=10, SEM=0.0786. Scale bar in (a**–**b), 200 µm. Scale bar, 500 µm in all images. P values were calculated with two-tailed unpaired Student’s t test.

**Figure 5.**
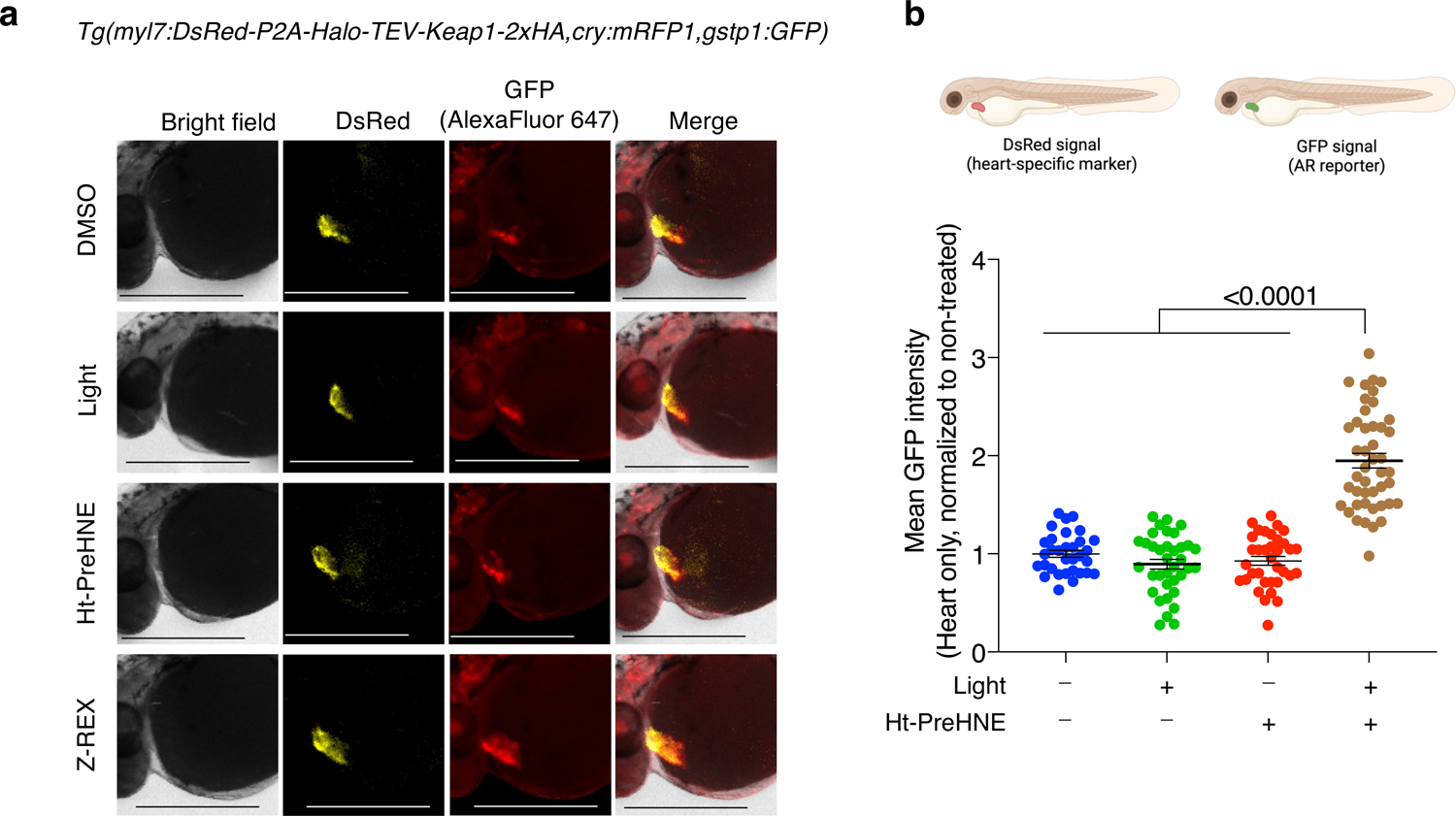
*Z-REX enables precision investigations into tissue-specific and POI-specific electrophile signaling.* The result shows that Z-REX precisely upregulates the AR signal only in cardiomyocytes, but not in other regions. (**a**) Representative images of (*myl7:DsRed-P2A-Halo-TEV-Keap1-2xHA, cry:mRFP1, gstp1:GFP*) fish treated with DMSO, light, 0.3 μM Ht-PreHNE, or Z-REX (with 0.3 μM Ht-PreHNE). Scale bar, 500 µm in all images. Whole fish images are in Extended Data Fig. 4a. Age: 34 hpf. (**b**) Quantification of mean GFP intensity. The quantification strategy is described in the discussion. Briefly, the heart region is defined by DsRed signal, which marks the cardiomyocytes. Z-REX-treated groups show two-fold higher AR signal than other negative control groups (DMSO, light or Ht-PreHNE alone). Also see Extended Data Fig. 4b for the quantification of AR signal in head, tail and whole fish. Age: 34 hpf. From left to right: n=32, SEM=0.0359; n=36, SEM=0.0502; n=35, SEM=0.0446; n=45, SEM=0.0748. P values were calculated with two-tailed unpaired Student’s t test.

**Figure 6.**
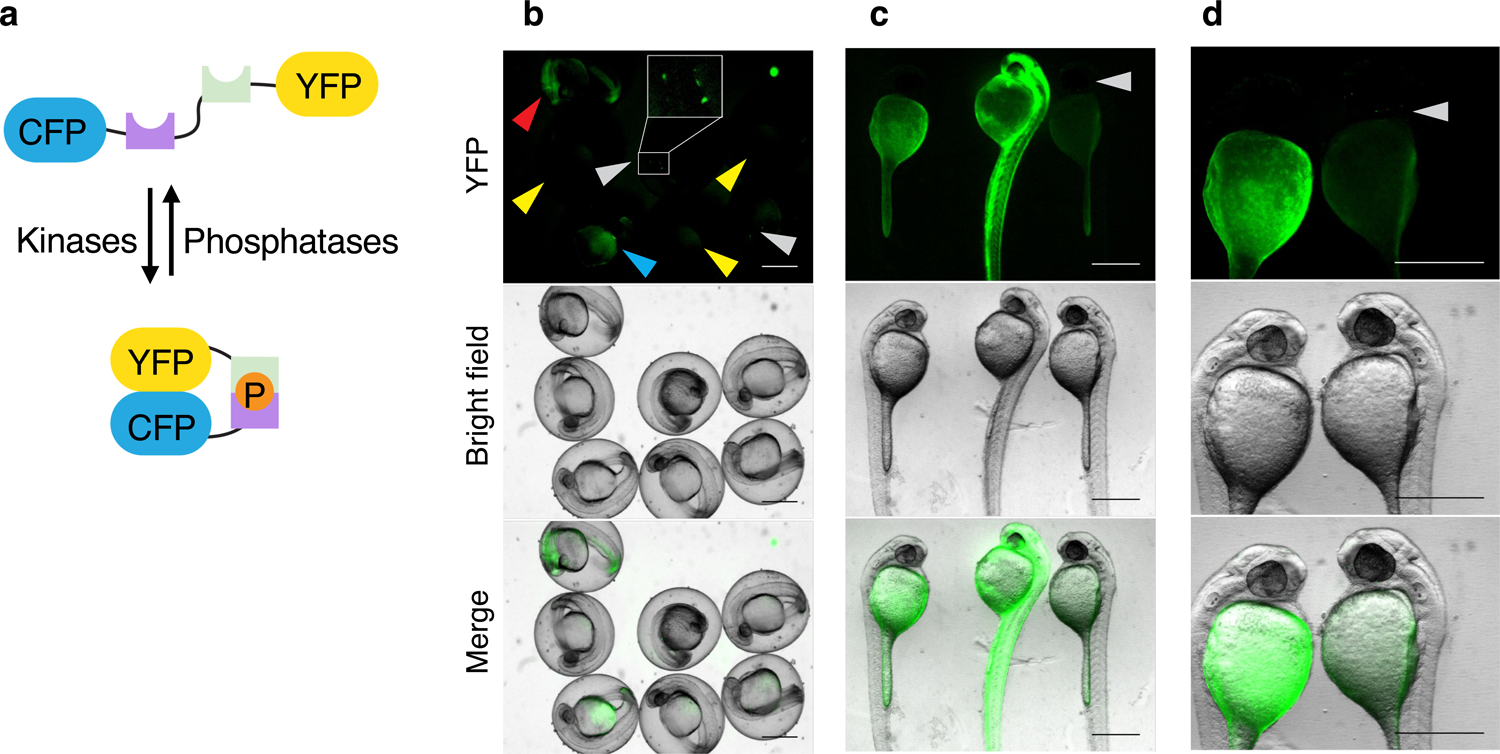
*Pre-screening of fish embryos using a representative fluorescent reporter construct*. In this example, Akt-kinase activity reporter construct (AktAR)^50^ was used. (**a**) Schematic of the ratiometric dual-fluorescence protein-based reporter, AktAR plasmid [The ratiometric FRET-based Akt-kinase-activity reporter (AktAR)^50^ directly reports phosphorylation of a peptide substrate (purple) by Akt isozymes (POI in this example) which is coupled to in an increase in FRET through interaction of the phospho-peptide-binding domain (green) to the phosphorylated peptide substrate]. (**b**) Embryos were co-injected with the mRNA encoding Halo-Akt and 25 ng/µL AktAR plasmid (into the single cell; see Extended Data Fig. 7f). After 24 h, embryos in the chorion were screened for fluorescence using a fluorescence stereomicroscope (ex. 495 nm, em. 500–550nm). **Grey** arrows indicate animals expressing reporter gene that are ideal for imaging; **yellow** arrows indicate wild-type animals; **blue** arrows indicate embryos expressing reporter gene and have with high yolk-sac fluorescence; **red** arrow indicates an embryo with reporter gene expression level highly deviated from other fish, and thus less likely to yield high-quality data. The zoom-in image of a fish ideal for imaging is shown in the white box. (**c**) Fish in (b) were dechorionated, placed on a 2% wt/vol agarose pad and imaged similarly, but at higher magnification than in (b). Fish furthest right (arrow) is best for confocal imaging. (**d**) Higher-magnification image showing one embryo with high fluorescence in yolk sac and the other in the head region (marked by **grey** arrow). We here only demonstrate the pre-screening images. Applications of this reporter plasmid in Z-REX and their results are reported in our previous work. Scale bar, 500 µm in all images.

### Overview of the Procedure

The Z-REX protocol is linked to a complementary paper detailing T-REX^25^. T-REX, which centers on on-demand redox-targeting protocol in cultured cells, involves: procuring the Halo-POI from a commercial repository of human or mouse Halo-tagged genes; screening libraries of POIs in a medium-throughput format; and carrying out downstream signaling assays. Here in Z-REX protocol, we provide a detailed procedure for: (i) how to transpose these commercial plasmids to a vector that can be used to create 5ʹ-capped mRNA that provides good expression in zebrafish (**Steps 1-37**); (ii) how to produce and microinject zebrafish embryos, and perform Z-REX-assisted Halo-POI-specific LDE delivery (exemplified by HNE) (**Steps 38-45**); (iii) how to validate expression and labeling of POI with HNE using Click-chemistry-based functionalization of an alkyne preinstalled on the Halo-targetable photocaged HNE (**Steps 46A-B**); and (iv) how to perform several informative downstream signaling assays (including in transgenic fish), such as qRT-PCR (**Step 46C**) and immunofluorescence (IF) imaging (**Step 46D**). We further describe how Z-REX Procedure can be adapted to use in transgenic zebrafish lines (see **Experimental Design** section for further details).

Similar to T-REX applied in live cell culture, Z-REX allows scrutiny of complex signaling pathways using a reasonably-standardized method applicable to zebrafish. In Z-REX, embryos express an ectopic protein (Halo-POI fusion protein) either through transient expression, or through transgenesis. The Halo-POI is able to undergo targeted pseudo-intramolecular delivery of a specific, endogenous LDE, such as HNE (Fig. 1), at a user-prescribed time. Delivery of LDE to the POI within the fish is achieved by photouncaging (2–5 min light exposure). Downstream phenotypic changes can be measured over the next 2–10 h. That being said, simply because a protein is designated an electrophile sensor from prior work, or even from T-REX in cultured cells, one cannot assume that protein will function as an electrophile sensor during Z-REX. Many reported electrophile sensor proteins are derived from *in vitro* assays performed with supraphysiological concentrations of reactive and toxic electrophiles, or for prolonged times^10, 21, 37^. On the other hand, the environments of cancer cells and developing fish embryos are not similar. We thus describe both protein biochemistry and chemical biology assays in live fish that evaluates protein-specific ligand binding that is general to both bolus LDE treatment and Z-REX application in fish. For all of the above procedures, we describe how to perform them from the perspective of a beginner in each area.

### Applications of Z-REX

Z-REX aims to study signaling pathways downstream of POI-specific HNEylation. We describe using fluorescence, qPCR, and reporter assays to enable a researcher to study principally cytoprotective antioxidant response (AR) signals^38^, including how this Procedure can be adapted for use in transgenic zebrafish lines to investigate tissue-specific effects of redox-dependent cytoprotective signaling. Furthermore, additional Z-REX-based approaches to study: proapoptotic signals triggered upon target-specific LDE modification (in which, for instance, innate immune reporter zebrafish lines were used for monitoring neutrophil- and macrophage-specific apoptosis^26^; and ratiometric FRET-based live imaging assays were used to monitor LDE-targeted kinase-isoform-specific inhibition in vivo^30^); upregulation of γ-H2AX due to hyperactivation of a positive regulator of a DNA damage-inducing ubiquitin-conjugating enzyme^15^ (in which, for instance, γ-H2AX expression levels were assessed by IF; and ubiquitylation by immunoprecipitation and western blot analyses).

Beyond individual POIs or pathways, Z-REX is compatible with large-scale systems-level investigations. For instance, recently, we used RNA-seq to compare the transcriptomic profiles of Z-REX-treated and non-treated embryos^26^. By targeting HNE (and other related native reactive metabolites) to Halo-Keap1 using Z-REX in this example, we discovered simultaneous downregulation of immune-relevant transcripts and upregulation of antioxidant response (AR) transcripts. These data were validated using qRT-PCR^26^. Other large-scale omics investigations, such as mass-spectrometry-based spatial proteomics and interactomics are proven compatible with other REX-technologies workflows^14, 26^, and are thus also anticipated to be readily coupled to Z-REX.

Last but not least, on-target and on-demand POI-LDE-modification enabled by Z-REX proves critically helpful in the identification of functionally-important players in signaling pathways. For instance, electrophile labeling of Keap1 stimulates upregulation of AR in a Nrf2-dependent manner. In Zebrafish, of the two Nrf2 paralogs, Nrf2a is believed to be particularly important for upregulation of AR in these circumstances^39^. Thus, if Z-REX-mediated upregulation of AR through electrophile modification of Keap1 occurs through the expected pathway, knockdown of Nrf2a should suppress AR upregulation. When this experiment was performed (using embryos prepared as above, but co-injected with a morpholino targeting Nrf2a, or a control morpholino), we found that only in embryos derived from the control morpholino was AR upregulation observed (Extended Data Fig. 5 and Table 1). This validates that the signaling pathway to upregulate AR in Z-REX is canonical. Similar experiments assessing on-target electrophile modification of other potential players in signaling pathways can be performed through (co-)injection of a morpholino enabling targeted knockdown of a specific protein of choice^26^.

## ALTERNATIVE METHODS

No other directly analogous method exists. We will discuss the current standard way to study RES and LDE signaling (such as HNE signaling) by bolus dosing, although this method does not bear a strong similarity to Z-REX^21^.

### Comparison with other methods

Bolus reactive-electrophile dosing has been used to study various aspects of fish development and toxicology^40^. This sort of experimental design reflects acute toxic exposure, possibly experienced by aquatic organisms in polluted environments, or can be used to model drug exposure, such as in high-throughput screening. With the reinvigoration of covalent electrophilic drugs^20, 22, 41, 42^, bolus electrophile dosing takes on new importance as dosing with reactive compounds like Tecfidera, an approved electrophilic drug to treat relapsing multiple sclerosis^20^, can be used as a platform to understand the complex time-dependent behaviors of these pleiotropic small molecules. However, many nuanced aspects of electrophile signaling are lost in the complexity of these studies incurred through, for instance: (i) changes in cellular redox state as LDEs react with cellular glutathione; (ii) uncontrolled covalent modifications inducing damage and crosslinking of multiple proteins by reactive electrophiles made available in supraphysiological and toxic quantities further deteriorating the cell health, oftentimes inducing cell death; (iii) generation of additional reactive species (from endogenous metabolic conversion of initially-administered LDEs) causing further damage to the underlying proteome and genome. As these issues accrue, nuanced aspects and ramifications of discrete electrophile modification events of individual electrophile-sensor proteins are very difficult to parse using the bolus-dosing setup^21^.

Unsurprisingly, Z-REX has been useful in unraveling the mechanism of Tecfidera and how it leads to loss of innate immune cells in fish^26^. Indeed, the main issue when considering a precision response to a specific reactive small-molecule is: what is/are the necessary (health-promoting) target(s) causing a specific signaling output? But when considering drug design and development, the question extends to what target(s) is/are beneficial and what target(s) is/are deleterious. The requirement of high concentrations of electrophile in bolus dosing renders this approach unable to satisfactorily answer either question: the experimental set up is far from accurate and gives responses that are hard to link to specific events due to long latency and low labeling of genuine kinetically-privileged electrophile-sensor proteins.

### Limitations and key considerations

Z-REX addresses many of the issues associated with T-REX: the expression of the ectopic protein is closer to (or below) endogenous levels; the setting is much more biologically relevant than cell culture; and there is also a much more scope to focus on locale-specific aspects of signaling in a live vertebrate animal, both in terms of precision target-engagement upstream, and pathway- or phenotypic-level resolution downstream. The principal issues with most examples of Z-REX are associated with the use of mRNA to introduce ectopic protein expression and the reliance of 365-nm light (despite low-power: 3.0 mW/cm^2^) to uncage the designated reactive electrophile. Use of mRNA allows expression of a non-native POI, which is a benefit because it allows a streamlined pipeline for the creation of Halo fusion genes via the human and mouse Halo-tagged ORFeome libraries (KAZusa collection, Promega). However, mRNA gives only transient expression that tapers off as the fish develops. We have observed POI-expression up to 2 days post mRNA-injection. Furthermore, mRNA-expression gives transient expression of POI ubiquitously and does not allow for tissue-specific expression.

We do however herein show tissue-specific upregulation of antioxidant response (AR) using transgenic embryos that remedies many of the aforementioned limitations (Fig. 5). Halo-POI transgenic fish also give more options for studying animals older than 2 days post fertilization, potentially to adult fish, depending on the target organ, and the photocaged small-molecule used. An alternative option is the use of CRISPR/cas9 to knock in Halo-POI to an endogenous gene locus under its native promoter. In terms of UV light exposure, 3–5 min light exposure (3 mW/cm^2^) at 30-70 hpf, is not detrimental to fish embryonic development, morphology, or viability^15, 26, 30^. However, UV-light’s overall tissue penetrance is poor, especially in older fish. These limitations once again restrict use to younger fish. Multi-photon Z-REX and engineering novel biocompatible photocage scaffolds, are a few potential solutions, and these new tools are being developed in our lab currently.

## EXPERIMENTAL DESIGN in Z-REX

### General considerations

The Z-REX protocol (particularly in transgenic animals) is not an optimal forum to screen electrophile sensitivity of a large number of POIs. This is because protein expression is relatively low in fish, which makes downstream processing time-consuming. Thus, it is best to begin Z-REX experiments already with a (few) defined POI(s) to test. These POIs could come from unbiased profiling or focused-screening assays. For unbiased screening of novel electrophile-sensor proteins, we suggest companion REX-technologies (e.g., Localis-REX^14^, G-REX^15^) that can quantitatively and rapidly identify native electrophile-sensor proteins with precise spatiotemporal, dosage, and electrophile-chemotype controls—unattainable by any other profiling methods. Localis-REX and G-REX applications have thus bar been demonstrated with HNE-sensor discovery both globally^15^ and in subcellular locales^14^. Should focused screening for electrophile sensitivity of a small set of POIs (∼50-100 POIs) be the goal, we recommend T-REX screening in cultured cells^30^, as opposed to Z-REX in *D. rerio*. Our identification of Akt3-HNE-signaling and development of novel isoform-specific covalent inhibitors stems from such a T-REX setup in HEK293T cells^30^, where we further took advantage of commercially-available Halo-POI ORF library encoding mouse and human genes (available from Promega).

It is further worth noting that Z-REX gives a much more contextually-relevant system for the measurements of downstream signaling responses, especially for phenotypic behaviors and biological facets such as metabolism are not recapitulated in cell culture models. Taken together, Z-REX can be put to its best benefit, if the best planning and prior validations are in place in deciding which Halo-POI, phenotypic or pathway-level responses are to be studied. We further recommend, particularly to those researchers more invested in downstream biological measurements, that the availability of reporter fish lines for measuring pathway or phenotypic responses be taken into consideration in planning for their Z-REX experiments. For instance, in this protocol, we use, as representative examples, a specific transgenic strain *Tg(−3.5gstp1:GFP)/it416b* reporter fish [hereafter *Tg(gstp1:GFP)*; available from the Japanese National BioResource Project (NBRP; http://www.nbrp.jp)]^43^ or a reporter plasmid bearing a specific fluorescent reporter of interest driven by the CMV promoter is used. Several viral promoters have been shown to work well in fish and should also be applicable to this specific protocol^44^.

Furthermore, prior to performing Z-REX, it is helpful to have validated POI-specific electrophile sensing, and electrophile-sensing residue(s) using T-REX in cultured cells [either deploying the human or zebrafish protein(s) as needed]. Moreover, basic pathway validations to help inform on reliable activity signatures of the target POI(s) (upregulated mRNAs, reporter genes, substrates) can also be more rapidly screened in cells using T-REX than in fish using Z-REX. In our experience, these signatures, such as changes in FRET in activity reporter, upregulation of DNA damage response, etc., have been transmissible from cells to fish^15, 30^. Based on these experiments, reporter fish strains can be sourced and ordered, necessary crosses can be performed (for instance, crossing into one’s chosen wild-type line), and preliminary validations can be undertaken during other preparation steps outlined below. In most instances, genotyping of lines will be relatively simple, as the reporter lines are usually fluorescent, and their function can be validated by measuring changes in fluorescence due to a stimulus (such as electrophilic stress or an inhibitor). The Halo-POI transgenic line we use here for Z-REX-assisted POI(Keap1)-targeted LDE delivery specifically in cardiac tissue, is also marked by fluorescent reporters (both in the specific tissue where Halo-POI is expressed and in the eye). The latter is a common strategy to mark transgenic fish that does not interfere with assays.

In terms of Halo-POI expression, this protocol (Z-REX) discusses two approaches: **(I)** generation of zebrafish embryos expressing an ectopic Halo-tagged POI through mRNA-injection; and **(II)** generation of transgenic zebrafish expressing a tissue-specific Halo-tagged POI. Transgenic zebrafish lines are commonplace, and are principally produced using CRISPR/cas9^45^ or Tol-2-mediated recombination^46^, and the latter is used in our case in Approach **(II)**. On the other hand, the procedure underlying Approach **(I)** is a rapid method to drive ubiquitous expression of ectopic POIs in numerous types of fish^47, 48^. Such processes are not tissue-specific, causing potential limitations and minimizing the benefits of using model organisms, at least from some perspectives. Nonetheless, our published data show that at least in certain cases, cell-specific effects can be observed, for instance, should the effect of protein-specific electrophile signaling manifest only in specific cells (e.g., Z-REX-assisted Keap1-specific HNEylation results in selective apoptosis of neutrophils and macrophages with no observable proapoptotic effects on other cells in the fish^26^, and Z-REX assisted Akt3-isoform-specific HNE-sensing inhibits Akt3 enzymatic activity that can be monitored in live fish, and suppresses Akt/FOXO signaling and endogenous downstream genes as analyzed by qRT-PCR^30^). By contrast, using Approach **(II)**, we demonstrate a method to upregulate heart-specific electrophile signaling using zebrafish with heart-specific Halo-Keap1 expression created using Tol-2 transgenesis. The nuts and bolts of mRNA-injection and transgenic procedures are the same, although creation of transgenic fish is more time-consuming than mRNA injection.

In terms of monitoring specific downstream pathways, we demonstrate the use of an injected signaling reporter encoded by a plasmid. Unlike mRNA injection that is relatively simple and gives relatively consistent levels of expression in the whole fish, plasmid injection is more complex and gives mosaic expression. The reasons for these differences include export mechanisms that selectively export mRNA over plasmid DNA from the yolk sac to the single cell, making mRNA distribute evenly throughout the whole embryo^49^. It is thus advisable to rapidly pre-screen injected fish for expression prior to imaging for quantitative analysis on a confocal microscope when the experiments involve injection of plasmids encoding fluorescent proteins^30, 50^. We suggest selecting fish with mosaic fluorescence in the head and body but little in the yolk sac (Fig. 6).

Finally, although Z- and T-REX offers significant advantages in terms of targeting efficiency and selectivity, compared to bolus experiments, where possible, it is best to validate specificity using hypomorphic mutants^30^. An ideal hypomorphic mutant is unable to sense electrophiles, and hence cannot trigger downstream signaling in an otherwise analogous system. The best way to design such a mutant is through identification of the sensing residue, and use of mutagenesis to genetically ablate the privileged sensing residue. So far we have only been able to identify sensor residues from assays performed in cell culture due to low POI-expression in zebrafish. However, in our previous publication, we have shown that only one privileged cysteine within Akt3 (8 total cysteines in humans) senses HNE both in cells (T-REX; via LC/MS-MS and mutagenesis studies) and in fish (Z-REX; via mutagenesis)^30^. Indeed, mutation of the privileged sensor residue furnishes a hypomorphic mutant—Akt3(C119S)—unable to trigger downstream signaling in both models when HNEylated. On the other hand, Keap1 is very cysteine-rich (27 total in human)^37^. In contrast to Akt3, many of these cysteines are electrophile sensors. We have successfully deployed a hypomorphic (C151S,C273W,C288E) mutant of Keap1—reported to be incapable of electrophile sensing but retaining wild-type Keap1’s ability to bind its interactors (e.g., Nrf2)^51^—to verify on-target and site(s)-specific electrophile-sensing and -signaling using Z-REX^26^. Should such mutants not be available, we have also developed the P2A system^26, 27^ that enables 1:1 expression of Halo and POI as separate domains with similar expression levels to the full-length Halo-POI fusion protein (Fig. 3). This system provides a convenient and reasonably-fair control, not available to other methods.

### Plasmid design (for transient Halo-POI expression through mRNA injection)

In order to validate Z-REX, the ability to visualize POIs via western blot is critical. We thus suggest that a construct first be designed bearing epitope tags. Here we describe the use of both HA (YPYDVPDYA) and FLAG (DYKDDDDK) tags. To enable proximity-guided electrophile sensitivity assessment, as well as absolute quantification of electrophile occupancy on POIs^27^, the POI and Halo should be linked by a protease-cleavable sequence such as that involving the TEV protease site (ENLYFQ/G; where “/” represents the protease cut site) that is present in the Halo ORFeome libraries (available, for instance, from Promega, Kazusa collection; backbone vector pFN21a). This design element allows for separation of Halo and POI, post lysis, a step that is required for the labeling of the POI and precise quantification of POI electrophile occupancy to be accurately determined by gel-based analyses we describe below. Halo-POI ORF libraries are available for human and mouse genes, although genes from other organisms can be incorporated using standard cloning methods^28^. Note: in this protocol, we will exclusively describe Halo-tagged genes of human origin. To transfer the DNA coding for Halo-TEV-POI, and simultaneously introduce a C-terminal 2xHA (tandem HA tags) or FLAG tag, into pCS2+8, the following sequence can be performed (Extended Data Fig. 6a): (*1*) the gene of interest is amplified by polymerase chain reaction (PCR) using primers that anneal in the forward direction to the N-terminus of Halo and the reverse direction to the C-terminus of the POI (the annealing temperature of each primer should be 65–68 °C, and this section of the primer should be approximately 20 bp long); the Halo primer has a 5’ flanking region that can anneal to the pCS2+8 plasmid. Note: the forward extender primer contains a Kozak sequence [CCACC(ATG)] and this MUST be retained to ensure translation); the reverse primer has a 5’ flanking sequence that introduces 2xHA (YPYDVPDYA) or FLAG (DYKDDDK; not included in primer table) tag. (*2*) The product of the first PCR reaction, after PCR clean up, will be “extended” sequentially using two sets of forward and reverse primers that ultimately give a PCR product that has around 40–60 bp overlap with the destination vector. (*3*) The product of these sequential reactions will then be used to prime a PCR reaction of linearized, empty pCS2+8 plasmid. The progress of the final “nicked plasmid”-forming reaction can be visualized on a gel, by measuring loss of the Halo-POI primer band and increase in the product band. (*4*) The resulting product, a “nicked plasmid” is circularized upon transformation into *E. coli*. Colonies with the desired insert can be screened by colony PCR or directly sequenced.

Once the plasmid is created, validate the full sequence of the construct (including the Kozak sequence, and annealing sites to the plasmid of interest at both the 3’ and 5’ ends). This is now ready to be transcribed. The pCS2+8 plasmid contains an SP6 primer site to create capped mRNA. Follow the mMessage mMachine® kit instructions to generate 5ʹmG-capped mRNA. We use PCR to amplify out the gene of interest with the SP6 and polyadenylation signal (often AAUAAA) intact rather than using digestion (Table 2, Primers for mRNA preparation). It is best to have some space between the SP6 site and the 5’ end of the amplicon for forming mRNA. We typically pool two preparations to furnish high-concentration mRNA in sufficient quantities that can be used directly for injection. The concentration we use is typically 1.2–1.6 mg/ml.

We have previously described a protocol to control for non-POI targeted electrophile released during photouncaging, and any effects of photochemical reactions occurring with or catalyzed by the probe in cell culture^25, 27, 31, 33^. This procedure involved co-expression of POI and Halo separately (i.e., unfused). These so-called “split” conditions gave no targeting to POI (e.g., human Keap1) and did not trigger downstream phenotypes (e.g., Nrf2-driven antioxidant response). In fish, we found such a protocol difficult to replicate because Halo-mRNA gave significantly higher expression than Halo-Keap1 (unlike in cell culture). We accordingly used the P2A, a self-cleaving peptide, to express *in situ* Halo-2xHA and Keap1-2xHA separately^26^, originating from a single fused mRNA with P2A sequence in-between the two genes^52^. Such a construct does not lead to modificaiton of the expressed POI, and is amorphic for Z-REX-associated phenotypes and does not lead to modification of the co-expressed POI^26^. Similar methods for protein expression like the internal ribosomal entry site (IRES) may be used, although these have not worked particularly well in zebrafish despite their versatility in cultured cells, at least to our experience^25, 33, 53, 54^. Bicistronic IRES expression also gives significantly different expression levels of the gene before vs. after the IRES, which is not optimal^55, 56^.

The Halo-2xHA-P2A-TEV-POI-2xHA construct can be created using a modified protocol to that outlined above (Extended Data Fig. 6b): (*1*) Halo is PCR-amplified from any Halo-containing vector (we use pFN21a for illustration). The forward primer is the same as the one used in Step 1 above, the reverse primer has 2xHA and P2A sequence added to the 5’-end (this gives product X in the figure); (*2*) POI is PCR-amplified using a forward primer with 5’-ends adding 2xHA and P2A with the reverse primer is the same as used for Step 1 above (this gives product Y in the figure); (*3*) the two products (X and Y) are mixed together in a self-priming PCR (i.e., all usual components, but no primers) to form the Halo-2xHA-P2A-POI construct; (*4*) this product is amplified using primers in Table 2; (*5*) additional primer-extension step as above; (*6*) the product of these steps is used to prime a PCR reaction of the whole plasmid. The crude product is transformed into *E. coli* and products are verified by colony PCR and sequencing.

Finally, although we do not touch on this point significantly in this manuscript, when designing transgenic lines, it is important to consider not only suitable controls and other necessary regulatory elements, but also the promoter used. The choice of promoter will determine the expression of the gene of interest. In this paper we chose to use the Myl7 promoter to drive expression of our POI in the heart. The heart was chosen because our AR-reporter line expresses well in the heart (otherwise, we could not readily monitor or measure AR signaling), and because the heart is a commonly-used model for studying redox stress and the roles of LDEs in oxidative stress are linked to cardiomyopathies, although underlying molecular mechanisms remain largely unknown^8, 10, 57^. Judicious choice of these elements is critical and should be done prior to creation of transgenic lines.

### Fish lines

Click chemistry-based assays (Steps 46A-B) and qRT-PCR (Step 46C) can be performed using embryos from Casper or wild-type zebrafish injected with Halo-POI mRNA and treated with Z-REX. IF assays (step 46D) can be performed using embryos from *Tg(gstp1:GFP)* and wild-type zebrafish injected with Halo-POI mRNA treated with Z-REX. We also demonstrate tissue-specific (heart) Z-REX targeting Keap1 using embryos from *Tg(myl7:DsRed-P2A-Halo-TEV-Keap1-2xHA,cry:mRFP1)* and *Tg(gstp1:GFP)*. Note: *myl7* is a cardiomyocyte-specific promoter; *cry:mRFP1* is a transgenesis marker used to determine whether the Tol2-mediated recombination is successful when producing the transgenic fish line. Sources of these lines are listed in **Materials**.

### Fish crossing

In the case of generating *Tg(gstp1:GFP)* embryos, parent transgenic fish should be always crossed into WT to generate heterozygous transgenic embryos. If self-crosses between *Tg(gstp1:GFP)* fish are inadvertently carried out, this may lead to homozygous reporter fish, which may have higher background fluorescence than heterozygotes. Note: the above analysis assumes that there is one copy of the reporter per fish. Outcrossing lines one receives into WT fish multiple times prior to starting your experiment (as we have already done), and examining inheritance patterns of heterozygotic fish crossed to wildtype to confirm Mendelian inheritance can help ensure that this be the case. Regardless, it is best to set up experiments in this manner to homogenize fluorescence background of the reporter strains used.

We illustrate this general concept using the following Punnett square below: here, *Tg* represents *gstp1:GFP*, and WT represents wildtype. If the transgenic parent is homozygous all progeny will be heterozygous transgenic: if the transgenic parent is heterozygous there will be 50:50 heterozygous transgenic:WT. Either way all transgenic fish will be isogenic. This is a general outcome (i.e., progeny genotypes from crosses of reporter strains into WT will always be heterozygous), so it is best to perform any cross to make reporter embryos this way to ensure isogenic progeny as far as the reporter is concerned.

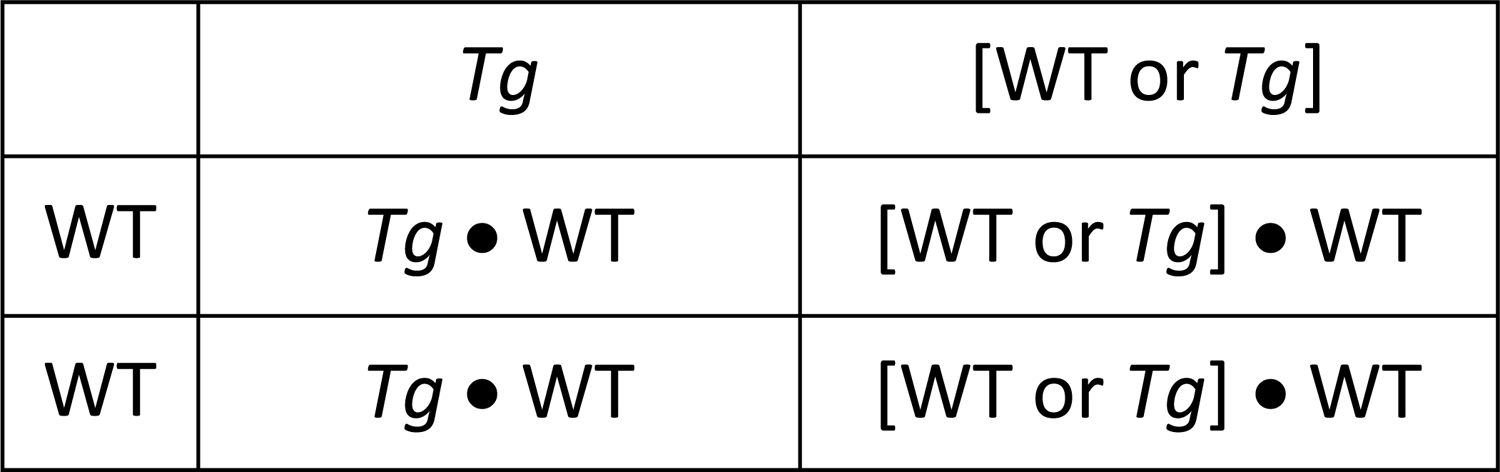

|  |  |  |
| --- | --- | --- |
|  | *Tg* | [WT or *Tg*] |
| WT | *Tg* • WT | [WT or *Tg*] • WT |
| WT | *Tg* • WT | [WT or *Tg*] • WT |

In the case of generating Tg(*gstp1:GFP;myl7:DsRed-P2A-Halo-TEV-Keap1-2xHA,cry:mRFP1*) embryos, crosses between Tg(gstp1:GFP) and Tg(*myl7:DsRed-P2A-Halo-TEV-Keap1-2xHA,cry:mRFP1*) always give heterozygous progeny. We illustrate this concept using the following Punnett square: here, *Tg 1* represents *gstp1:GFP* and *Tg 2* represents *myl7:DsRed-P2A-Halo-TEV-Keap1-2xHA,cry:mRFP1*. The progeny carrying both transgenes can be isolated by the fluorescent markers (GFP and DsRed).

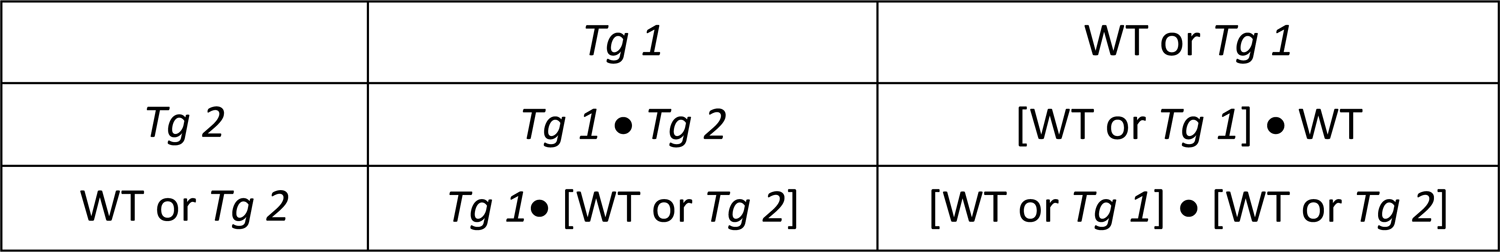

|  |  |  |
| --- | --- | --- |
|  | *Tg 1* | WT or *Tg 1* |
| *Tg 2* | *Tg 1* • *Tg 2* | [WT or *Tg 1*] • WT |
| WT or *Tg 2* | *Tg 1* • [WT or *Tg 2*] | [WT or *Tg 1*] • [WT or *Tg 2*] |

### Fish age / developmental stage

Z-REX using mRNA injection is performed on embryonic fish. All experiments here are performed on embryos 1–1.5 days post fertilization (dpf). The longevity of ectopic proteins expressed using mRNA is variable, although we have found several proteins are detectable in fish embryos 1-1.5 dpf. Should older embryos, or potentially adult fish, be required (or tissue specific effects be of particular interest), transgenic animals are likely a better choice.

### Microinjection: mRNA [with morpholino (MO) or plasmid DNA]

Zebrafish embryos can be injected (1) directly into the yolk sac with mRNA (and MO, Table 1) between the 1- and 4-cell stages (Extended Data Fig. 7e) or (2) into cells with plasmid (and mRNA) at the 1-cell stage (Extended Data Fig. 7f) to give robust ubiquitous expression (Extended Data Fig. 7 and 8). Because mRNA is unstable, we suggest only loading the needle once fish eggs have been set up in the injection wells. Until this point, maintain the mRNA on ice (or likely better, dry ice). Zebrafish spawn at the beginning of their daily light cycle, so injections should be planned for the morning. For assessing delivery by biotin pulldown or in-gel fluorescence analyses, both leveraging Click chemistry (*vide infra*), we use injected fish derived from multiple spawnings occurring over the whole breeding period (approximately 3 h). For phenotypic assessments, it is best to maintain the injected embryos at a similar age group (i.e., spawned at a similar time). If one round of egg laying provides a small number of eggs, these can be used for negative controls (non-injected) or euthanized.

### Treatment with photocaged precursor and uncaging of active electrophile

The photocaged precursor (0.1-6 µM) does not affect development and is not toxic^26, 30^. Embryos can be treated with the photocaged precursor immediately post injection. For phenotypic assays, 3 washings of the embryos for 30 min each is sufficient to remove excess compound. The principal issue we have observed with the photocaged precursor is that excess can lead to high background fluorescence in live fish imaging or IF assays, unless used at low (1 µM) concentration. As low as 0.1 µM of photocaged precursor has been used effectively. We do not recommend using higher concentrations of photocaged molecules (>6 µM), as this may cause non-specific triggering of some responsive phenotypes.

### Dechorionation (and deyolking)

Dechorionation is principally used for imaging of embryos. The chorion is a transparent shell that surrounds the embryo and is pointed out in Extended Data Fig. 7b. Embryos in the chorion can be difficult to visualize because they do not lay in a single focal plane (for instance, compare Figs. 6a vs**. 4b**). Thus, dechorionating (at 1 dpf) greatly assists imaging. Note: dechorionation should not be performed until 1 dpf. For whole mount immunofluorescence (IF) (*vide infra*), it is critical to dechorionate prior to fixing, otherwise fish adopt a curved shape that cannot be smoothed out. This is best achieved using sharp tweezers that can pierce the chorion and can be expanded to help create a hole therein. The principal error when dechorionating is bursting the yolk sac, which is irreparable.

Dechorionation and deyolking must be performed prior to running western blot analysis. The yolk sac (pointed to in Figs. 2b and d) is densely packed with yolk proteins that can dominate the zebrafish lysates and lead to non-specific binding of the antibody. The two manipulations (dechorionation and deyolking) can be done in a single step with tweezers, detailed in **Step 46A (iii)**.

### Cy5 and biotin click assays

Click assays involve preparing the fish as above and treating one set with Ht-PreHNE and the other with DMSO. 28.5 hpf, fish are rinsed 3 times over 1.5 h (total), then both sets are exposed to light for 5 min. Fish are dechorionated and deyolked, harvested, and pelleted. Pellets can be frozen in liquid nitrogen and stored at −80 °C for at least 1 week, or carried on to the next step. The approximate yield protein per 30 hpf deyolked fish is approximately 2 µg.

For Cy5 Click in-gel fluorescence analysis (previously detailed in our T-REX cell-based protocol^25^ and recently extended to zebrafish embryos), approximately 25 µg of lysate is required. For biotin Click pulldown from fish lysate, at least 170 µg of lysate is required. It is typically best to aim for 1.3-fold more fish than the minimum requirement: (i.e., for Cy5 Click, **minimum** 17 fish; for biotin Click, **minimum** 110 embryos). Biotin Click assay is currently the most sensitive assay we use to identify POI-specific delivery in zebrafish.

### Downstream response assay and IF analysis

Here we use *Tg(gstp1:GFP)* reporter fish available from NBRP (*vide supra*) (Figs. 2b–e**, 4, 5,** and Extended Data Figs. 2–3, and **4**). We found that although live fish are fluorescent relative to non-transgenic embryos of the same age, and that the background green fluorescence, especially from the non-fluorescent parts of the fish, significantly affects quantitation. We therefore developed an IF method to detect GFP that uses a red fluorophore-conjugated secondary antibody to give enhanced signal-to-noise. This method shows fluorescence primarily in the head, heart and tail as previously reported for native GFP^43^.

Analyze fluorescence using ImageJ (NIH). Whole fish, head (eyes excluded), or tail (median fin fold) fluorescence can be quantitated by measuring the average fluorescence intensity in a freehand selection in ImageJ. For assessing tissues, organs or regions that cannot be accurately determined with bright-field image, one can select the region according to tissue- or organ-specific marker-protein expression (e.g., fluorescent protein signal or fluorophore-conjugated antibody staining).

In some cases, fluorescent proteins are still detectable after the fish is fixed, such as the DsRed signal we co-express in the heart of transgenic fish (Fig. 5a, and Extended Data Fig. 4a). In cases where the fluorescent protein is totally inactive after fixation, antibodies recognizing the fluorescent protein can be used for staining to visualized protein expression.

### Validation of Z-REX in transgenic embryos

Microinjection of mRNA is not required in Z-REX using transgenic embryos, although all other procedures remain largely the same as described above. However, transgenic fish must be created using methods outlined above that are discussed at length elsewhere^58^. Here we use an in-house-generated line that expresses Halo-TEV-Keap1 in the heart, due to Halo-TEV-Keap1 expression being driven by the zebrafish myl7-promoter (Extended Data Fig. 9). Transgenicity is marked by expression of DsRed, which is co-expressed with Halo-TeV-Keap1 by virtue of a P2A sequence. These fish are crossed with *Tg(gstp1:GFP)* reporter fish and data are reported for fish that are transgenic for both markers (i.e., GFP- and DsRed-positive). Z-REX-treated embryos are immunostained for GFP (Alexa-647) and fluorescence is quantified in the heart, head, and the tail, which is a particularly responsive region of the fish (as shown above), and compared across controls versus Z-REX fish. Orthogonal assays can bolster results from reporter fish.

Specifically, we established a method to measure upregulation of AR signaling in both the tail and the head of zebrafish embryos separately. This involves separation of head and tail from euthanized fish, followed by separate mRNA extractions.

## MATERIALS

### FISH LINES

- Casper, wild-type zebrafish, such as AB (can be obtained from ZIRC) or Brian’s wildtype (a strain of wt-zebrafish that shows low pigmentation). This strain is available upon request, from the Fetcho laboratory, Cornell University (USA), or the Aye laboratory, EPFL (Switzerland).
- *Tg(gstp1:GFP)* (can be obtained from the Japanese National Bioresource Project; http://shigen.nig.ac.jp/zebra/index_en.html)
- *Tg(myl7:DsRed-P2A-Halo-TEV-Keap1-2xHA,cry:mRFP1)* (produced by Prof. Yimon Aye’s lab, EPFL, can be obtained upon request) (genotyping data and live fish images in Extended Data Fig. 9)

**! CAUTION** All procedures related to zebrafish studies at Cornell University conform to the US National Institutes of Health guidelines regarding animal experimentation and were approved by Cornell University’s Institutional Animal Care and Use committees. All procedures related to zebrafish studies at EPFL comply with the Swiss regulations on animal experimentation (Animal Welfare Act SR 455 and Animal Welfare Ordinance SR 455.1), and were performed at the EPFL zebrafish unit (cantonal veterinary authorization VD-H23).

### REAGENTS

#### Reagents for mRNA and plasmid preparation

- Chemically Competent *E. coli* (e.g. Invitrogen TOP10, cat. no. C404003; Agilent Technologies XL-10, cat. no. 200315)
- Tryptone (IBI Scientific, cat. no. IB49182)
- Yeast extract (Fisher, cat. no. BP1422-2)
- Agar (Fisher, cat. no. EC232-658-1)
- Chloramphenicol (Goldbio, cat. no. C-105-5)
- Kanamycin (Goldbio, cat. no. K-120-5)
- Ampicillin (Goldbio, cat. no. A-301-5)
- mMessage mMachine® SP6 Transcription Kit (Ambion cat. no. AM1340)
- Phusion Hotstart II DNA polymerase (ThermoFisher Scientific, cat. no. F549L; includes 5x HF buffer)
- dNTPs (Qiagen cat. no. 201912)
- pFN21a-Halo-TEV-Keap1 (Promega, Kazusa Collection)
- pCS2+8 empty vector (e.g., Addgene #34931)
- Dimethyl sulfoxide (DMSO) (Fisher, cat. no. D128-500)
- Standard reagents for DNA gel electrophoresis
- Orange G (Sigma, cat. no. O3756)
- Agarose LE (Gold Biotechnology cat. no. A201-100)
- Ethidium bromide (Sigma cat. no. E1510)
- PCR products purification kit (BioBasic cat. no. BS363)
- Miniprep kit (Bio Basic, cat. no. BS614)
- Rnase*Zap*® RNA decontamination solution (ThermoFisher Scientific, cat. no. AM9780)
- Glass beads (Sigma, cat. no. 45-G1145-100G)
- TRIzol Reagent (ThermoFisher, 15596018)
- SuperScript III reverse transcriptase (ThermoFisher Scientific, cat. no. 18080085)
- RNAseOUT™ recombinant ribonuclease inhibitor (ThermoFisher Scientific, cat. no. 10777019)
- Diethyl pyrocarbonate (DEPC), High purity (VWR, cat. no. E174-110G)
- EcoRI (NEB cat. no. R0101S)
- Dpn1 (NEB cat. no. R0176S)
- Oligo(dT)_20_ (Order the following DNA sequence from IDT: TTTTTTTTTTTTTTTTTTTT)
- pCS2+8 empty vector (Addgene: #34931)

#### Reagents for Z-REX

- pCS2+8 Halo-TEV-Keap1-2xHA (see Steps 1–33)
- pCS2+8 Halo-2XHA-P2A-TEV-Keap1-2xHA (See Steps 1–33)
- HaloTag-targetable precursor to HNE(alkyne) [Ht-PreHNE (see original protocol^25^)]
- HNE(alkyne) (see original protocol)
- cOmplete, Mini, EDTA-free Protease Inhibitor Cocktail (Roche cat. no. 11836170001)
- Trypsin inhibitor from *Glycine max* (soybean) (Sigma, cat. no. T9003)
- 4-(2-hydroxyethyl)-1-piperazineethanesulfonic acid (HEPES; Fisher, cat. no. BP310-1)
- Tris(2-carboxyethyl)phosphine hydrochloride (TCEP-HCl, Gold Biotechnology cat. no. TCEP1)
- Zirconia beads (BioSpec cat. no. 11079107zx)
- *t*-Butanol (Fisher, cat. no. A401-1)
- Sulfo-Cy5 azide (Lumiprobe, cat. no. B3330)
- Biotin-dPEG11-azide (Quanta Biodesign, cat. no. 102856-142)
- Copper Tris[(1-benzyl-1H-1,2,3-triazol-4-yl)methyl]amine (Cu-TBTA; Lumiprobe, cat. no. 21050)
- Copper sulfate pentahydrate (Sigma, cat. no. 209198-100G)
- Lithium dodecyl sulfate (LDS; Chem-Impex, cat. no. 00269)
- Sodium dodecyl sulfate (SDS; Teknova cat. no. S9974)
- β-Mercaptoethanol (BME; Sigma-Aldrich, cat. no. M6250)
- NP-40 (CalBioChem cat. no. 492016)
- Standard reagents for protein gel electrophoresis and western blotting
- Bradford Dye (BioRad cat. no. 5000205)
- Dimethyl sulfoxide (DMSO, Fisher, cat. no. D128-500)
- Triton X-100 (Fisher, cat. no. BP151-100)
- Tween 20 (Fisher, cat. no. BP337-500)
- BSA (Fisher, cat. no. BP1600-100)
- Heat inactivated FBS (Sigma, cat. no. F2442)
- His6-TEV-S219V (see original protocol^25^)
- Paraformaldehyde (Sigma, cat. no. P6148)

**! CAUTION** Carcinogen: Handle inside fume hood with gloves, goggles, and lab coat, when preparing formaldehyde solution.

- 1x Dulbecco’s phosphate buffered saline (no calcium, no magnesium) (1x PBS)(Gibco, cat. no. 14190144)
- Hank’s Balanced Salt Solution (HBSS, see reagent setup below)
- Mouse monoclonal antibody to Keap1 (Abcam cat. no. ab119403; RRID: AB_10903761; IF, 1:200-500); this antibody can detect human Keap1 (expressed ectopically) and both isoforms of zebrafish Keap1
- Goat polyclonal antibody to GFP, FITC-conjugated (Abcam cat. no. ab6662; RRID: AB_305635; IF, 1:1000)
- Rat monoclonal antibody to HA-HRP (Sigma, cat. no. H3663; RRID: AB_262051; WB, 1:500; IF, 1:500)
- Mouse monoclonal antibody to β-actin HRP (Sigma cat. no. A3854; RRID: AB_262011; WB, 1:30000)
- Donkey polyclonal secondary antibody to mouse IgG-H&L, AlexaFluor 568-conjugated (Abcam cat. no. ab175472; RRID: AB_2636996; IF, 1:1000)
- Donkey polyclonal secondary antibody to rat IgG-H&L, AlexaFluor 568-conjugated (Abcam cat. no. ab175475; RRID: AB_2636887; IF, 1:500)
- Rabbit polyclonal secondary antibody to goat IgG-H&L, AlexaFluor 568-conjugated (Abcam cat. no. ab175707; RRID: AB_2923275, IF, 1:2000)
- Donkey polyclonal secondary antibody to goat IgG-H&L, AlexaFluor 647-conjugated (Abcam cat. no. ab150131; RRID: AB_2732857; IF, 1:2000)
- High capacity streptavidin agarose (Thermo Scientific cat. no. 20359)

### EQUIPMENT

- Standard equipment for zebrafish husbandry and breeding
- Standard equipment for nucleic acid agarose gel electrophoresis
- Thermal cycler (e.g. Applied Biosystems Veriti)
- Hand-held UV lamp with 365-nm light (e.g., Spectroline ENF 240C)
- Accumax UV meter (Spectroline, cat. no. XS-365)
- Petri dishes and multiwell plates (Celltreat 229692 and 229506)
- Disposable transfer pipets (e.g. Fisher cat. no. 13-711-9D)
- Fluorescence stereomicroscope (e.g., Leica M165 FC)
- Transparent red-colored film plastic sheet (e.g., Pangda, 11.7 by 8.3 Inches, red)
- Confocal microscope (e.g., Zeiss LSM 710)
- Microscope for micro-injection (Olympus cat. no. SZ61)
- Microscope light source (Olympus cat. no. KL 1600 LED)
- Standard equipment for protein gel electrophoresis and western blotting
- ChemiDoc-MP imaging system (Bio-Rad) or Fusion FX imager (Vilber)
- Picoliter injector (e.g., Harvard Apparatus PLI-90A) or Pneumatic PicoPump (WPI cat. no. SYS-PV820) with Micromanipulator (Narishige cat. no. MN-153)
- Flaming/Brown Micropipette Puller (Sutter Instrument Co. cat. no. P-97)
- Capillary tubes (World Precision Instruments, Inc. cat. no. TW100 F6) or VWR cat. no. HARV30-00200)
- Hemocytometer (Hausser Scientific cat. no. 1492) or glass stage micrometer (Meiji Techno. cat. no. MA285)
- Injection plates
- Microinjection mold (Adaptive Science Tools cat. no. TU-1)
- Dumont #5 forceps (Fine Science Tools cat. no. 11252-40)
- Microloader tips (Eppendorf cat. no. 930001007)
- Nanodrop spectrophotometer (NanoDrop 1000)
- Vortexer (e.g. Vortex-Genie 2)
- Refrigerated centrifuge (e.g. Eppendorf 5417R)
- Microwave (Sunbeam)
- Heated ultrasonic bath (e.g. VWR cat. no. 89375-470)
- ImageJ/FIJI (NIH)
- Glass backed imaging plate (for confocal microscopy) in vitro scientific

### REAGENT SETUP

- **Hank’s Balanced Salt Solution (HBSS):** For a 10x concentrate, dissolve 8 g NaCl, 0.4 g KCl, 0.19 g CaCl_2_•2H_2_O, 0.12 g MgSO_4_•7H_2_O, 0.25 g MgCl_2_•7H_2_O, 0.035 g Na_2_HPO_4_•2H_2_O, 0.06 g KH_2_PO_4_, and 0.35 g NaHCO_3_ in 1 litre water and filter sterilize. Dilute to 1x prior to use. Store concentrated solution at 4 °C and make fresh every 3 months. Dilute to 1x and warm up to 28.5 °C prior to use.
- **LB Medium:** Dissolve 10 g NaCl, 10 g tryptone and 5 g yeast extract in 1 l water. Autoclave to sterilize. Store at room temperature (15–22 °C) up to 1 week.
- **TCEP solution:** Prepare 100 mM TCEP in 500 mM HEPES pH 7.6. Aliquot and store at −20°C for up to 6 months. Do not re-freeze after thawing.
- **Cy5:** Prepare 0.5 mM Cy5 in DMSO. Aliquot and store at −20°C for up to 6 months, protected from light.
- **20% wt/vol LDS:** Dissolve 20 g LDS in 100 ml water. Vortex and heat (50–60 °C) as necessary to ensure complete dissolution. This can be sonicated and reheated if a precipitate forms. It is stable indefinitely at room temperature.
- **20% wt/vol SDS:** Dissolve 20 g SDS in 100 ml water. Vortex and heat (50–60 °C) as necessary to ensure complete dissolution. This can be sonicated and reheated if a precipitate forms. It is stable indefinitely at room temperature.
- **100 mM CuSO_4_:** Dissolve 1.25 g CuSO4.5H2O in 50 ml water. Filter sterilize. Store at room temperature indefinitely.
- **1x PBST:** Supplement 1x PBS with 0.1% vol/vol Tween-20. Store at room temperature (15–22 °C) indefinitely.
- **Fish lysis buffer:** Dissolve 50 mM HEPES (pH 7.6); 1% Triton X-100; 0.1 mg/ml soybean trypsin inhibitor in ddH_2_O. Chill on ice. Add 2x Roche protease inhibitors and 0.1 mg/ml soybean trypsin inhibitor just before use. These need to be used within <30 minutes of being made, for optimal efficacy.
- **IF Blocking Buffer:** Supplement 1x PBST with 10% vol/vol heat-inactivated FBS and 2% wt/vol BSA. Aliquot and store at −20°C for up to a month.
- **1x PDT:** Supplement 1x PBST with 0.3% vol/vol Triton-X100 and 1% vol/vol DMSO. Store at 4 °C for up to a month.
- **Methanol:** Chill methanol at −20°C. It is stable indefinitely.
- **4 and 10%** (wt/vol) **Paraformaldehyde (PFA):** Add 20 g of PFA powder to 160 ml PBS, then heat the solution to 50 °C with stirring in a fume hood. Slowly add the saturated NaOH _(aq)_ until the solution is clear, and adjust the pH to 7 with 1-5 N HCl (instead of pH meter, use pH paper or test strip to measure pH since they can be discarded afterward). Fill the solution to 200 ml with PBS to make a 10% w/v PFA stock solution. Aliquot 4-6 ml solution in each 15-ml falcon tube, and store them at −20 °C for up to 6 months. Before the fixation experiment, simply dilute the stock solution with PBS to 4% w/v.

**▴CRITICAL STEP** Formaldehyde can be oxidized in air. Always use aliquoted PFA solution, do not repeatedly freeze and thaw the solution. PFA stock solution can be prepared in a lab equipped with fume hood and water bath.

**! CAUTION** Formaldehyde is a known carcinogen. Handle in a fume hood with protective clothing (goggles, gloves, labcoat).

- **10 mM dNTP mix:** Make 100 µl stock of 10 mM dNTPs by mixing 10 µl each of 100 mM dATP, dCTP, dGTP and dTTP. Bring volume to 100 µl with ddH_2_O. Mix and aliquot. Aliquots can be stored at −20°C for up to 6 months.

- **DEPC treated water**: Add 1 ml of DEPC to 1 l ddH_2_O. Stir overnight. Autoclave for 45 min to remove DEPC. Let water cool to room temperature before using. DEPC treated water can be stored at 4 °C for up to 1 month.
- **Tris-Acetate-EDTA buffer (TAE; 50x solution):** Dissolve 242 g Tris free base, 18.61 g Disodium EDTA, and 57.1 ml Glacial Acetic Acid, in ddH_2_O and make up to 1 liter. Do not pH. 1x solution has a final concentration of: 40 mM Tris, 20 mM Acetate and 1 mM EDTA and typically has a pH around 8.6. It is stable for up to 6 months, but check regularly for contamination.
- **Laemelli buffer (4x)**: Dissolve 200 mM Tris pH 6.8, 8% wt/vol SDS, 40% vol/vol glycerol, 0.4% wt/vol bromophenol blue in ddH_2_O. Add 6% vol/vol BME immediately prior to use. Do not store the buffer plus reducing agent for any longer than 30 min^59^ at room temperature, although this can be stored at −20°C for 3 months.
- **Ethidium bromide solution**: Dissolve 100 mg EtBr in 10 ml ddH_2_O. EtBr solution can be stored at room temperature for > 6 months.
- **DNA loading dye (5x):** Dissolve 15% vol/vol glycerol and 50 mg of Orange G in 50 ml of ddH_2_O. Loading dye can be store at room temperature > 6 months.
- **TCEP:** Only add TCEP just prior to use. TCEP stock solution is pre-made as a 100 mM stock in Argon-saturated 500 mM HEPES (pH 7.5), aliquoted, flash frozen and stored at −20 °C. These one-shot aliquots should be taken out of freezer just prior to use (do not rest the solution on ice as it may precipitate), and should not be freeze-thawed.
- SDS-PAGE running buffer (10X): Dissolve 300 g Tris free base, 1440 g glycine and 100 g SDS in 10 litre of ddH_2_O. Store at room temperature indefinitely.

### EQUIPMENT SETUP

- **Injection plates:** Dissolve 2 g agarose in 100 ml 1X HBSS. Microwave until agarose is fully dissolved and allow to cool until it is warm, but not scolding to the touch (i.e., 50-60 °C). Pour agarose into petri dishes (∼3/4 full) and gently float a microinjection mold on top of the agarose. Leave to solidify. Injection plates can be reused several times if refrigerated between uses. Always microwave (for 4 s) prior to using for injection such that plates are warmed slightly.
- **Injection needle:** Prepare needle with Flaming/Brown Micropipette Puller (heat: 520 units; pull strength: 60 units; velocity: 70 units; delay: 155 units; pressure: 550 units; ramp: 530 units. Units of these values are default settings of the machine).

**▴CRITICAL STEP** Store prepared microinjection needles on modeling clay or similar to prevent needle breaking. Needles can be prepared ahead of time and stored indefinitely.

- **ChemiDoc-MP or Fusion FX Imaging system setup for Cy5 fluorescent gel imaging** Set the Cy5 excitation source as red epi illumination and emission filter as 695/55 filter.

## PROCEDURE

Setting up plasmid constructs for expression. **•** TIMING 7 d

**▴CRITICAL** See Extended Data Fig. 6 for schematic illustration.

### Primer design

**1|** Design primers to amplify the gene of interest (in this case, His_6_-Halo-TEV-Keap1-2xHA from pFN21a vector encoding Halo-TEV-Keap1) and insert His_6_-tag on the N-terminus and 2xHA-tag on the C-terminus (Table 2, Primers for gene amplification). **Note:** the Keap1 ectopically expressed in this paper is the human gene.

**▴ CRITICAL STEP** Melting temperature of the overlapping part of the primer should be around 65–68 °C and a GC content around 50%. Around 20 base pairs (bp) overlap with the gene of interest is required for proper priming

**▴ CRITICAL STEP** If cloning pCS2+8 His_6_-Halo-2xHA-P2A-TEV-Keap1-2xHA, design two pairs of amplification primers; Halo and Keap1 are amplified by two separate PCR reactions from pFN21a Halo-TEV-Keap1 using two different pairs of primers. (Table 2, Primers for gene amplification; Extended Data Fig. 6, lower panel).

**2|** Design primers to extend the N- and C-terminal flanking regions (Table 2, Extension primers 1).

**▴ CRITICAL STEP** For tandem or longer tag insertion (e.g. 2xHA), the tag sequence can be split, and inserted sequentially by step 1 and 2.

**▴ CRITICAL STEP** For tandem tag insertion, change the nucleobase of the wobble position without affecting the protein coding sequence. e.g., in one HA sequence (YPYDVPDYA) use TA**C** to code for tyrosine; in the second HA sequence (YPYDVPDYA), use TA**T** to code for tyrosine and so on. Wobble position in bold.

**▴ CRITICAL STEP** If cloning pCS2+8 His_6_-Halo-2xHA-P2A-TEV-Keap1-2xHA, design two pairs of amplification primers; Halo and Keap1 are extended separately using two different pairs of primers (Table 2, Extension primers 1).

**3|** If an additional extension step is required to give the required minimum overlap with destination plasmid, design primers to increase annealing between the “megaprimer” and the destination plasmid (Table 2, Extension primers 2).

**▴ CRITICAL STEP** The goal of steps 1-3 is to create linear DNA that has the desired gene, its epitope tags, and flanking regions with overlap to the destination vector on the 3’ and 5’ ends. The number of PCR steps required to achieve this will differ based on GC richness, length of primers used, etc. However, an absolute minimum of 30 bp overlap with the vector backbone is required for efficient insertion of the gene-of-interest in the desired linearized vector. Longer overlap (>50 bp) greatly increases the insertion efficiency.

**▴ CRITICAL STEP** Primers for cloning pCS2+8 His_6_-Halo-2xHA-P2A-TEV-Keap1-2xHA and pCS2+8 His_6_-Halo-2xHA-P2A-TEV-Keap1-2xHA are the same in this step.

### Ligase-free cloning

4|Dilute pFN21a-Halo-TEV-Keap1 plasmid to a concentration of 10 ng/µl in ddH_2_O.

5|In 33.5 µl ddH_2_O, add 10 µl 5x High Fidelity buffer, 1 µl of 10 µM forward primer, 1 µl of 10 µM reverse primer (Table 2, Primers for gene amplification), 1 µl dilute pFN21a-Halo-TEV-Keap1 (template), 1 µl of 10 mM dNTPs, 2 µl DMSO and 0.5 µl Phusion Hotstart II polymerase.

**▴ CRITICAL STEP** Mix 5x HF buffer thoroughly by vortexing (>5s) before taking out desired volume.

**▴CRITICAL** Also see Step 1 for experimental setup details for two different plasmid constructs.

6|Gently pipette up and down to mix the PCR mixture.

7|PCR-amplify using the following cycling conditions: 98 °C for 50 s, 30 cycles of [98 °C for 10s, 65 °C (or melting temperature of primers) for 30s, 72 °C for 45s (or required extension step for your gene)], 72 °C for 2 min, and 8°C afterwards.

**▴ CRITICAL STEP** Use 15 s extension time per 1 kilo bp of DNA.

**8|** Meanwhile, add 0.3 g agarose to 30 ml 1x Tris Acetate EDTA (TAE) buffer in an Erlenmeyer flask. Plug the mouth loosely with Kimwipes. Microwave the mix for around 1 min to completely dissolve agarose.

**! CAUTION** Glassware and melted agarose will be hot. Handle with care.

**9|** Let the agarose cool to 40–50 °C. Add 3 µl EtBr (10 mg/ml, stock concentration). Mix by swirling.

**▴ CRITICAL STEP** Don’t add EtBr when the agarose is too hot; don’t let agarose solidify; and ensure gel box is clean prior to pouring gel.

**! CAUTION** EtBr is a known carcinogen: wear gloves. Alternative formulations that are considered less dangerous can also be used, although they are not used by our laboratory.

**10|** Prepare the gel cast and pour the agarose. Remove or burst any air bubbles using a pipette tip. Let the gel solidify.

**11|** After PCR cycling is complete, add 1 µl of 5x DNA loading dye to 4 µl of the PCR reaction mix. Mix and load on the solidified agarose gel. Run gel electrophoresis for 5 min at 120 V in 1x Tris acetate EDTA (TAE) buffer. Visualize gel, check for primer dimers, then run for a further 6 min and assess the size of your amplicon.

**▴ CRITICAL STEP** We run the gel for a short time so that we can easily detect primer dimers. Primer dimers are visible as diffuse bands that run around 100 base pairs. These will negatively affect downstream steps and you should perform gel extraction if these are visible.

?TROUBLESHOOTING

**12|** Perform PCR cleanup to isolate the product. Measure concentration using nanodrop.

▪ **PAUSE POINT** Isolated PCR product can be stored at −20 °C for up to a week.

**13|** Set up a PCR reaction similar to Steps 5-6 except use 1 µl each of forward extender primer and reverse extender primer (10 µM, Table 2, Extension primers 1), and 2 µl of product from Step 12 as the template (up to 200 ng). Bring the volume to 50 µl using ddH_2_O.

**▴CRITICAL** See also Step 2 for experimental setup details for two different plasmid constructs.

**▴ CRITICAL STEP** If cloning pCS2+8 His_6_-Halo-2xHA-P2A-TEV-Keap1-2xHA, an additional PCR reaction using two extended PCR products (His_6_-Halo-2xHA and P2A-TEV-Keap1-2xHA with flanking regions annealing to each other) is performed without adding primers to generate His_6_-Halo-2xHA-P2A-TEV-Keap1-2xHA with N- and C-terminal flanking sequences (Extended Data Fig. 6, lower panel).

**14|** Set up a PCR cycling reaction as specified in Step 7.

**15|** Check that the PCR reaction is successful using gel electrophoresis and perform PCR cleanup to as outlined in Steps 8–12 to isolate the ‘megaprimer’.

**▴CRITICAL STEP** If multiple bands are present in the gel, load entire product from Step 15 and gel extract the correct band. If recovery is low, you can pool multiple reactions.

▪ **PAUSE POINT** Isolated PCR product can be stored at −20 °C for up to a week.

?TROUBLESHOOTING

**16|** (Optional) An additional extension step may be required to generate sufficient overlap with the destination vector. Repeat Steps 13–15 with the second set of extension primers if desired (Table 2, Extension primers 2).

**17|** Linearize pCS2+8 vector with EcoRI restriction enzyme. Follow manufacturer’s instruction.

**18|** Set up a PCR reaction using 10 µl 5x HF buffer, 200 ng of megaprimer, 50 ng of linearized plasmid, 1 µl dNTPs, 2 µl DMSO and 0.5 µl Phusion Hotstart II polymerase. Adjust total volume to 50 µl using ddH_2_O.

**19|** Split the PCR mix into 3 separate PCR tubes after mixing, 16 µl in each tube.

**20|** Set up a PCR reaction as in Step 7 except increase the elongation time 2 min 20 sec. Lower annealing temperature to 61 °C.

**21|** Take out each tube at the intervals of 12, 18 and 24 cycling steps.

**22|** Analyze samples using gel electrophoresis to determine formation of product and the consumption of megaprimers.

**▴CRITICAL STEP** Aim to transform any samples that have showed significant depletion of the mega primer. If the reaction is run without megaprimer for many cycles, this tends to lead to very high molecular weight PCR products (>10 kbp) and is typically not successful upon transformation.

**▴ CRITICAL STEP** Dpn1 digestion can be performed prior to Step 23 to remove template plasmid (digested pCS2+8, prepared from E. coli). Dpn1 digests methylated DNA (template plasmid), but not the PCR product. This reduces the numbers of colonies containing re-ligated or undigested template plasmid in Step 24.

▪ **PAUSE POINT** Crude PCR mix can be stored at −80 °C for 1–2 days.

?TROUBLESHOOTING

**23|** Transform all the remaining products into chemically-competent *E. Coli* (e.g., XL10 or TOP10). Grow overnight at 37 °C.

### Colony PCR

**24|** Label 10–20 single colonies from a plate with a unique identifier (e.g., 1–20).

**25|** Take half of each single, isolated colony using a clean pipette tip. Avoid taking up Agar from the plate.

**26|** Resuspend each ‘half-colony’ separately in tubes labeled with their specific identifier (e.g., 1–20) containing 50 µl 1x HF buffer.

**27|** Vortex samples (which should be cloudy due to *E. coli*), and then heat to 98 °C for 5 min. Centrifuge samples at 10000 × g for 1 min at room temperature.

**28|** Make a master mix of 1.1X PCR buffer containing all requisite components (see Step 5) except template.

**▴ CRITICAL STEP** It is best to use two primers that anneal uniquely to the inserted gene (e.g., Primers for gene amplification in Table 2).

**29|** Add 1 µl lysed *E. coli* to 10 µl 1.1X PCR buffer containing all requisite components except template.

**▴ CRITICAL STEP** Also perform both negative and positive controls, respectively, as follows: either non-transformed *E. coli* or empty pCS2+8 (negative controls); and plasmid containing gene of interest (positive controls, e.g., pFN21a Halo-TEV-Keap1).

**30|** Run the PCR reaction as described in Step 7.

**31|** Analyze on gel. There should be no amplicons for non-transformed *E. coli* or empty pCS2+8, and clones that undergo efficient PCR should give an amplicon with the same molecular weight as the positive control plasmid.

?TROUBLESHOOTING

**32|** Grow colonies that produced a PCR band in Step 31 in 5 ml LB media with 100 μg/ml Ampicillin overnight at 37 °C.

**33|** Isolate plasmids using plasmid miniprep kit and fully sequence to verify correct insertion.

▪ **PAUSE POINT** Purified plasmid can be stored at −20 °C for several years.

Creating mRNA **e** TIMING 0.5 d

**▴CRITICAL** Clean all pipettes, gloves, and work area with RNaseZap.

**34|** Follow mMessage mMachine® SP6 in vitro transcription kit protocol for two preps.

**▴CRITICAL** We recommend using PCR to amplify out the gene of interest with the SP6 and polyadenylation signal intact rather than using digestion (Table 2, Primers for mRNA preparation).

**35|** Resuspend mRNA pellet in nuclease free water (provided with the kit).

**36|** Determine concentration of mRNA by nanodrop. Concentration of RNA should be 1–2 mg/ml. Volumes of RNA are calibrated assuming a concentration of 1.5 mg/ml. Vary them accordingly. Check for RNA quality using gel electrophoresis.

**▴CRITICAL** RNA does not run at expected molecular weight because of secondary structures.

**37|** Aliquot immediately and store at −80 °C.

▪ **PAUSE POINT** Purified mRNA can be stored at −80 °C for 1–2 months.

Setting up fish crosses and obtaining fish eggs • TIMING 0.5 d

**38| Option A** describes crosses of Casper or WT fish used for qRT-PCR, and mosaic reporter fish formed from injection of plasmid DNA and mRNA. **Option B** elucidates the crosses of transgenic reporter fish line, such as antioxidant response (AR)-reporter line, *Tg(gstp1:GFP).* The transient POI-expression in reporter fish embryos can be obtained through mRNA Injection. **Option C** demonstrates how to obtain transgenic fish embryos both carrying reporter gene (e.g., *gstp1*:GFP on AR reporter) and tissue-specific Halo-POI (e.g., *myl7*:Halo-Keap1 in cardiomyocytes).

A. Generating Casper or WT fish embryos
  i. Set up 6 crossing pairs (Casper or WT fish) in 6 separate breeding tanks, with slotted-bottom inserts to allow eggs to fall through, and with males and females separated by a removable divider). **▴ CRITICAL STEP** Typically, one breeding tank contains 1 male and 1–3 females. Based on the experimental needs, the number of tanks can also be adjusted. **▴ CRITICAL STEP** Fish need a shaded place to breed, so adding foliage to the tanks is recommended.
  ii. The next morning (typically 9–10 AM), when ready to inject eggs, remove the dividers on 3 tanks and collect eggs from each tank every 15–20 min, by moving each set of fish to a new tank and pouring the fish water containing eggs through a strainer and rinsing out the embryos from the stainer into a 10-cm Petri dish. When the fish in the first 3 tanks stop producing eggs, remove the dividers on the second set of 3 tanks and proceed the same as above. Pool the tanks as required. **▴ CRITICAL STEP** If a specific set of breeding fish do not lay eggs, individual sets can be pooled, which typically promotes egg laying. ?TROUBLESHOOTING
B. Generating heterozygous *Tg(gstp1:GFP)* reporter fish embryos
  i. Set up mating pairs in breeding tanks as above, except in one set of 3 tanks, only the male should be a *Tg(gstp1:GFP)* reporter fish, whereas in the other set of 3 tanks, only the females should be *Tg(gstp1:GFP)* reporter fish. In each case the cross is into WT (the parent line for the *Tg(gstp1:GFP)* reporter strain is WT). **▴CRITICAL** The crossing setup ensures that the all *transgenic* progeny are heterozygous for the reporter strain minimizing variation. Note: the above analysis assumes that there is one (actively expressed) copy of the reporter per fish. Outcrossing lines one receives into WT fish multiple times prior to starting one’s experiment (as we have done here), and examining inheritance patterns of heterozygotic fish crossed to wildtype to confirm Mendelian inheritance can help ensure that this be the case. Regardless, it is best to set up experiments in this manner to homogenize fluorescence background of the reporter strains used. ▴**CRITICAL STEP** Tanks are typically set up such that 1 tank contains 1 male and 1–3 females. 6 tanks are typically used but based on experimental requirements (e.g., sample size), the number of tanks can be adjusted.
  ii. The next morning (typically 9–10 AM), when ready to inject eggs, remove the dividers on 3 tanks and collect eggs every 15–20 min, by moving fish to a new tank, pouring the fish water containing eggs through a strainer, and rinsing out the embryos from the stainer to a 10-cm Petri dish. When the fish in the first 3 tanks stop producing eggs, remove the dividers on the second set of 3 tanks and proceed the same as above. ▴**CRITICAL STEP** If a specific set of breeding fish do not lay eggs, only pool tanks with transgenic males or transgenic females. Do not pool transgenic males and transgenic females together since this will result in progeny with multiple copies of the transgenic reporter, which likely have progeny with higher fluorescence background than those derived from crosses with wt. ?TROUBLESHOOTING
C. Generating heterozygous Tg(gstp1:GFP;myl7:DsRed-P2A-Halo-TEV-Keap1-2xHA,cry:mRFP1) fish embryos
  i. Set up mating pairs in breeding tanks as in **(B)**, except for replacing WT fish with *Tg(myl7:DsRed-P2A-Halo-TEV-Keap1-2xHA,cry:mRFP1)* fish. ▴ **CRITICAL STEP** Typically, one tank contains 1 male and 1–3 females. Set up 6 tanks is recommended. Based on the experiment purposes, the tank number can be varied.
  ii. The next morning (typically 9–10 AM), remove all the dividers and collect eggs every 15-20 min, by moving fish to a new tank and pouring the fish water containing eggs through a strainer, and rinsing out the embryos from the strainer to a 10-cm Petri dish. In all instances, embryos can be stored prior to injecting in 10% vol/vol HBSS in a petridish. When the fish in the first 3 tanks stop producing eggs, remove the dividers on the second set of 3 tanks and proceed the same as above. ?TROUBLESHOOTING

Microinjection, for embryos collected from 38 (A) and (B) **•** TIMING 3h

**39|** Option A describes microinjection of mRNA whereas Option B describes a protocol to co-inject mRNA and plasmid DNA in zebrafish embryos.

A. Microinjection of mRNA
  i. Align freshly laid embryos (for mRNA injection they should be 1–4 cell stage) in agarose injection plates (see Equipment Setup) filled with 10% vol/vol HBSS with blunt forceps. When using only mRNA, injection should be directly into the yolk sac (Extended Data Fig. 7e) of embryos at the 1–4 cell stage. Setting up the embryos is relatively simple as there is no specific orientation required to achieve optimal injection. 100–150 embryos can be aligned in a single 10-cm plate. ▴**CRITICAL STEP** Ensure the back pressure (0.2-0.5 psi) is on, and injection pressure is set at 25-35 psi.
  ii. Load mRNA (2 µL, 1.5 mg/ml) into a micro-injection needle using a microloader tip. **▴ CRITICAL STEP** mRNA should be kept on ice, or dry ice, until needed
  iii. Attach the needle to the microinjector apparatus.
  iv. Snap off the tip of the needle using sharp forceps.
  v. Lower the needle through a drop of mineral oil on a hemocytometer so that it almost touches the surface and release a drop. Based on the hemocytometer markings, calibrate the size of the drop to 2 nl (Extended Data Fig. 7a).
  vi. Rinse the tip by lowering into a petri dish of HBSS, then inject embryos. **▴ CRITICAL STEP** Here, we describe the mRNA-loading procedures with micromanipulator (Narishige, MN-153), and the needles are prepared using capillary tubes (VWR, HARV30-0020) and needle puller (SUTTER, P-97). If using different microinjectors, the mRNA-loading procedures may be slightly different. **▴ CRITICAL STEP** The best way to avoid clogging is to keep the needle in 10% HBSS at all times: while harvesting injected embryos, have a 10-cm dish half-filled with 10% HBSS to place needle in. Before next round of injections, clear needle twice by releasing two drops. If the needle is clogged between the injections, repeat step (ii)-(vi) to load mRNA into a new needle.
  vii. Once embryos have all been injected, rinse embryos out of the injection wells using a squirt bottle (filled with 10% HBSS) and collect them into a petri dish.
  viii. If more eggs are left, add them to the injection plate and arrange them in the wells as before. During this time the needle should be kept in a petri dish filled with 10 % HBSS. When ready to inject clear the needle twice, and inject.
  ix. When all embryos have been injected, separate them into respective 10-cm dishes, for instance, separating different constructs expressed in accordance to a specific experimental design, number of conditions required, etc.
  x. In a dark room, replace the media with 30 ml 10% HBSS with or without 1 µM Ht-PreHNE, place the dishes in an incubator (28.5 °C), covered with alumnium foil to incubate the embryos in dark. **▴ CRITICAL STEP** Ht-PreHNE solution should be made up in 10% HBSS prior to adding to embryos and this should be performed in a dark room. Place the dishes in a box covered in foil. Ht-PreHNE stock solutions should not be freeze-thawed. **▴ CRITICAL STEP** For all the control and experiment groups, including bolus dosing group, fish should be harvested all at the same time. This is to ensure that all fish are the same age for comparing the outcomes with those from Z-REX. ?TROUBLESHOOTING
B. Co-microinjection of mRNA and plasmid DNA **▴ CRITICAL STEP** Use embryos at the 1-cell stage; later embryos can be used, but this leads to very poor expression levels of the reporter and mosaic expression of the mRNA.
  i. Align embryos in small batches with the single cells pointing upwards towards the needle (Extended Data Fig. 7f). Note this is different to the random orientation used in mRNA injection.
  ii. Prepare an injection cocktail, just before loading the needle, by mixing the mRNA and DNA such that a 30 μg/ml plasmid and 1.4 mg/ml mRNA concentrations are achieved.
  iii. Load the mix and calibrate the injection as described in Steps 39A (ii)–(viii). ▴**CRITICAL STEP** Ensure the back pressure (0.2-0.5 psi) is on, and injection pressure is set at 25-35 psi.
  iv. Lower the needle such that it is close to the cell, then advance the needle swiftly such that it penetrates the single cell. **▴ CRITICAL STEP** Piercing the cell membrane typically requires more force than piercing the yolk-sac membrane, so the needle should be advanced into the cell swiftly and confidently. Being too timid when attempting a cell injection typically leads to the embryo rolling or the needle breaking.
  v. Inject the mix.
  vi. When all embryos have been injected, separate into respective 10-cm dishes, for instance based on different constructs expressed, depending on specific experimental design, etc.
  vii. In a dark room, replace the media with 30 ml 10% HBSS with or without 1 µM Ht-PreHNE, and incubate the embryos at 28.5 °C in an incubator (covered with aluminium foil to protect from light). **▴ CRITICAL STEP** Ht-PreHNE solution should be made up in 10% HBSS prior to adding to embryos and this should be performed in a dark room. Place the dishes in a box covered in foil. We further recommend that as many one-shot stock-aliquots of Ht-PreHNE as are necessary should be first pooled and mixed together to homogenize any potential stock concentration differences across different tubes. Ht-PreHNE stock solutions should not be freeze-thawed. **▴ CRITICAL STEP** For all the control and experiment groups, including bolus dosing group, fish should be harvested all at the same time. This is to ensure that all fish are the same age for comparing the outcomes with those from Z-REX. ?TROUBLESHOOTING

Z-REX procedure **e** TIMING 2–8h

**40|** At 28.5 h, move the embryos (that are protected from light with foil) into a dark room with red light illumination. Pour off the 10% HBSS. Collect any 10% HBSS that cannot be poured off using a pipette. Be careful not to lose embryos. This step is usually carried out on embryos aged 28.5 h. However, a similar protocol is likely applicable to younger, and older embryos, especially if transgenic embryos are used in the case of the latter.

**41|** Replace with fresh 10% HBSS (30 ml for a 10 cm dish).

**▴ CRITICAL STEP** For this step only use 10% HBSS without methylene blue (methylene blue can be added to 10% HBSS prior to this). Make sure the 10% HBSS is at 28.5 °C.

**42|** Incubate for 30 min at 28.5 °C in the dark.

**43|** Repeat Steps 41–42 two more times.

**44|** When last rinse is completed, turn on UV light.

**45|** Expose embryos to UV light (see below for timings). This step is usually performed on 30 hpf embryos. A similar protocol is likely applicable to younger, and older embryos.

**▴ CRITICAL STEP** Pre-warm the UV-lamp by turning it on for ∼5-min before using. Make sure the petri dish lids are taken off when shining light.

The following options are suggested:

A. If performing Cy5 (Step 46A) or biotin click (Step 46B) procedure, fish can be irradiated directly in the 10 cm dish for 5 min, then chilled in ice.
B. If performing downstream signaling or phenotypic investigations, e.g., qRT-PCR (Step 46C) or IF assays (Step 46D), separate compound treated and untreated embryos into two sets each (a 6 well plate can be used for irradiation). One set of compound treated fish and untreated fish should be irradiated with light (3 min) and the other should be kept in dark. If performing bolus dosing add the requisite compound at the required time such that all samples can be harvested at the same time.

**46|** Downstream analyses

A. Assessing targeting on POI by click coupling with Cy5 azide **•** TIMING 7 h **▴ CRITICAL STEP** Samples should be kept on ice if having been exposed to T-REX conditions or bolus dosing.
  i. Immobilize around 30 embryos per condition by chilling them on ice for 10 min. **▴ CRITICAL STEP** This step is carried out right after step 45 (B).
  ii. Put fish embryos in a 10-cm petri dish in around 30 ml HBSS. Using a light microscope, move fish into middle of dish by swirling or gently pushing them using a transfer pipette. **▴ CRITICAL STEP** If using compound treated samples that have not been exposed to light, one must use red light illumination. This can be achieved by placing a red filter over the illuminated base or light source.
  iii. Perform dechorionation and deyolking by holding the chorion with one pair of sharp forceps (use your weaker hand) then with another pair of forceps, aiming directly at the yolk sac and piercing the chorion, penetrating the yolk sac. In one smooth motion drag the forceps away from the body of the fish, while maintaining contact with the yolk sac to burst it. **▴ CRITICAL STEP** The chorion must be peeled away from the remaining body of the fish. The solution will become increasingly cloudy as more fish are prepared. This is not an issue.
  iv. Swirl the plate to help congregate the fish bodies in the center of the dish while pushing the lighter chorions to the sides. Collect the embryos into an Eppendorf tube.
  v. Remove the solution, and add 1 ml ice-cold 50 mM HEPES buffer (pH 7.6).
  vi. Pipette the fish up and down 3 times to help dislodge debris and mix the yolk proteins with the solution (the lighter debris can float in the solution). Let the embryos settle to the bottom. Remove buffer. **▴ CRITICAL STEP** Care must be taken not to lose embryos during wash steps.
  vii. Repeat Steps (v)-(vi). **▪ PAUSE POINT** After final removal of wash buffer, embryos can be flash frozen in liquid nitrogen and stored at −80 °C for 2 weeks.
  viii. Resuspend fish in lysis buffer, making sure 2x Roche protease inhibitors and soybean trypsin inhibitor have been added immediately before use. We suggest aiming for an initial concentration of approximately 2.5 mg/ml lysate. For 30 embryos, one can expect around 60 µg protein, thus lysis should be performed in 24 µl.
  ix. Add zirconia beads to the fish pellets and vortex for 20 s.
  x. Freeze samples in liquid nitrogen. Thaw mixture rapidly either by warming by hand, or by putting in warm water. Repeat the preceding 3 steps (vortex, freeze, thaw) two more times. Then centrifuge lysates at 21,000 × g at 4 °C for 10 min. **! CAUTION** Liquid nitrogen can cause low temperature burns. Liquid nitrogen can also cause Eppendorf tube caps to open with extreme force: wear goggles and make sure lids of tubes are closed firmly before performing freeze thaw.
  xi. Remove clarified supernatant from tubes and add to fresh prechilled Eppendorf tubes. Measure protein concentration using Bradford assay relative to BSA. **▴ CRITICAL STEP** Make sure all foliage etc. is removed from the embryos prior to lysis. These can release dyes etc. during lysis that can give erroneous (too high) readings in the Bradford analysis.
  xii. Dilute the clarified lysate to 1 mg/ml in a final volume of 25 µl with lysis buffer [comprising in final concentrations: 50 mM HEPES (pH 7.6), 0.3 mM TCEP], and 2.5 µl of 2 mg/ml stock solution of His_6_-TEV-S219V. Incubate at 37 °C for 30 min. For samples that do not require TEV protease treatment, TEV protease addition and heating is omitted. **▴ CRITICAL STEP** The optimal concentration of lysate protein is ∼1.0 mg/mL. High concentration of lysate protein causes failure of Click coupling. Precipitates may appear upon adding His_6_-TEV-S219V. Gentle resuspension is necessary to ensure the success of Click coupling. For samples without light exposure, His_6_-TEV-S219V is omitted. Do not freeze-thaw TEV protease enzyme stock solution.
  xiii. Prepare 10x Click coupling master mix: 10% wt/vol SDS, 10 mM CuSO_4_, 1 mM Cu(TBTA), 100 µM Cy5 azide and 20 mM TCEP. **▴ CRITICAL STEP** All the concentrations above are critical to the success of Click coupling. Mix well to make sure that the solution is homogeneous. **▴ CRITICAL STEP** Add TCEP solution to the mix just before use.
  xiv. Add 5% vol/vol t-BuOH and 2.5 µl 10x Click coupling master mix to the TEV-protease-treated lysate from 46(A)(xii). Incubate the resulting mixture at 37 °C for 30 min.
  xv. Quench with 10 µl of 4x Laemmli buffer that contains 6% vol/vol BME and further incubate for 5 min at 37 °C. Load 20 µl into each well of 10% polyacrylamide gel, and resolve by electrophoresis. **▴ CRITICAL STEP** 4x Laemmli buffer should be warmed in advance to ensure homogeneity. Fresh SDS-PAGE running buffer should be used to reduce background signal. It is recommended to rinse the wells of polyacrylamide gel (remove buffer in the wells using a P-200 pipet with a loading tip, repeat for 4–5 times) before loading the samples to enhance signal-to-noise ratio.
  xvi. Upon completion of the gel-electrophoresis, rinse the gel with ddH_2_O (twice, 5 min each rinse) and analyze for Cy5 signal using a gel imager with Cy5-detection capabilities or similar instrument (Fig. 3c). ?TROUBLESHOOTING
B. **(B)** Assessing targeting on POI by click coupling with Biotin pull down **•** TIMING 3 d
  i. Manually deyolk, dechorionate and lyse around 120 embryos per condition. Follow Steps 46 A (i)–(xi) except for using 100 µl lysis buffer for 120 embryos. **▴ CRITICAL STEP** This step is carried out right after step 45 (A).
  ii. Remove 30–60 µg of zebrafish lysate for input signals. Add Laemmli buffer to input samples, flash freeze in liq N_2_, and store at −80 °C.
  iii. Dilute the remaining lysates to 1 mg/ml (using Bradford dye relative to BSA standard) and aliquot at least 170 µg of each into a 2 ml Eppendorf tube.
  iv. Add TEV protease (His6-TEV-S219V in final concentration: 0.2 mg/ml) to each sample and incubate at 37 °C for 30 min. For samples that do not require TEV protease treatment, this step is omitted.
  v. Make a 10x stock solution containing, in final concentrations, 10% wt/vol SDS, 10 mM CuSO_4_, 1 mM Cu-TBTA, 1 mM biotin-azide and 20 mM TCEP. **▴ CRITICAL STEP** Add TCEP just before use.
  vi. Add 5% vol/vol *t*-BuOH, and 10% vol/vol cocktail from step (v) to each sample. Vortex mixture for 10 s, centrifuge at 10,000 × g for 5 s and incubate at 37 °C for 30 min. After 15 min incubation, add an extra 1 mM TCEP, vortex for 10 s, centrifuge at 10,000 × g for 5 s and continue with incubation for another 15 min. The incubation time is 30 min in total.
  vii. Add EtOH (that had been prechilled at −20 °C; approximately 600 µl, if 170 µg protein used), to give rise to 75% vol/vol final concentration of EtOH.
  viii. Vortex and incubate at −80 °C for at least overnight
  ix. ▪ **PAUSE POINT** Samples can be maintained at −80 °C for at least one week. There is no appreciable improvement in precipitation after 24 h.
  x. Centrifuge samples (21,000 × g) at 4 °C for 1 h.
  xi. Remove supernatant and add 1 ml EtOH (−20 °C), vortex for 15 s, and centrifuge again (21,000 × g) for 10 min.
  xii. Repeat Step x.
  xiii. Remove EtOH and add chilled 75% vol/vol EtOH in water, vortex for 15 s, and centrifuge again (21,000 × g) for 10 min.
  xiv. Repeat Step xii.
  xv. Remove supernatant and add 1 ml acetone (−20 °C), vortex for 15 s, and centrifuge again (21,000 × g) for 10 min.
  xvi. Remove acetone and allow samples to air dry. **▴ CRITICAL STEP** Do not let pellet dry out completely. A trivial amount of acetone is not an issue. However, completely drying the pellet may lead to protein aggregation, rendering the pellet difficult to redissolve in the following step.
  xvii. Add 8% wt/vol LDS in 50 mM HEPES (pH 7.6), 1 mM EDTA (100 µl is recommended), and sonicate until samples are fully dissolved.
  xviii. Centrifuge the samples at 21,000 × g for 5 min and remove supernatant (10 µl is mixed with Laemmli buffer as flowthrough samples) and place it in a clean tube. There should be very little to no pellet in the samples. Some pellet is sometimes observed, and should be ignored if it does not re-solubilize after sonication. These pellets are likely fragmented genomic DNA or hydrophobic protein aggregates.
  xix. Dilute samples 16-fold in 50 mM HEPES (pH 7.6), to give a final concentration of LDS = 0.5% wt/vol.
  xx. Add the diluted sample to 100 µl bed-volume prewashed streptavidin high-capacity resin. **▴ CRITICAL STEP** Streptavidin high-capacity resin is prewashed with washed 1 time with 1 mL ddH2O, and 2 times with 1 mL 0.5% LDS in 50 mM HEPES pH = 7.6.
  xxi. Incubate mixture for 4–6 h at rt with end-over-end rotation.
  xxii. Centrifuge resin at 1,500 × g for 2 min, and remove supernatant completely. Replace with 1 ml of 0.5% wt/vol LDS in 50 mM HEPES (pH 7.6). Incubate mixture for 30 min at rt with end-over-end rotation. **▴ CRITICAL STEP** To improve washing steps, 900 µl wash buffer can be removed using a P-1000 pipette, then the remaining buffer can be removed using a P-200 pipette fitted with a gel loading tip. This minimizes loss of resin. **▴ CRITICAL STEP** Wash steps of up to 1% wt/vol LDS can be used if your protein shows high extent of non-specific binding. Binding step must always be carried out in no more than 0.5% wt/vol LDS.
  xxiii. Repeat Step xxi two times.
  xxiv. Elute bound proteins in 2x Laemmli buffer (40 µl) at 100 °C. If loading gel straight away, centrifuge samples at 18,000 × g for 5 min prior to loading. Load as much of sample as possible (around 28 µL on a 10 lane mini gel). ▪ **PAUSE POINT** Samples can be stored at −20 °C at least overnight. If samples are stored this way, heat to 100 °C for 3 min, and centrifuge at 18,000 × g for 2 min prior to loading.
  xxv. Analyze by western blot (Fig. 3c). ?TROUBLESHOOTING
C. Assessing downstream transcriptional response by qRT-PCR **•** TIMING 3 d **▴CRITICAL** This is best performed by two people, one dechorionating embryos and the other separating fish and extracting mRNA.
  i. 2 h post light exposure, immobilize fish by chilling them on ice.
  ii. Dechorionate fish by holding the embryo with one pair of sharp forceps (use your weak hand). Then, with another pair of sharp forceps pinch a piece of the chorion and pull it away to release the fish from chorion.
  iii. (Optional) Use a pair of sharp forceps to separate the head from the tail.
  iv. **▴CRITICAL** This step is done because there is a large disparity between ARE-driven mRNA in the head and the tail. Thus whole fish mRNA is dominated by mRNA from the head.
  v. Pool 10–12 heads and tails (or 5–7 whole fish) in 1.5-ml tube to give one single replicate (we typically perform 3–5 replicates).
  vi. Add glass beads and 1 ml of TRIzol reagent to each tube.
  vii. **▴CRITICAL** We have found zirconia beads are not optimal for this extraction procedure, so we suggest glass beads.
  viii. Vortex samples for 2 min. ▪ **PAUSE POINT** mRNA in trizol can be stored at −80 °C for 1–3 weeks.
  ix. Isolate total RNA per manufacturer’s instruction for TRIzol reagent (the protocol from Thermo Fisher). Analyze RNA quality using gel electrophoresis. ?TROUBLESHOOTING
  x. Reverse transcribe 600 ng of total RNA using SuperScript III reverse transcriptase and Oligo-dT for priming. Follow manufacturer’s instructions (the protocol from Thermo Fisher).
  xi. Follow appropriate protocol for primer design and qRT-PCR^60^.
  xii. Analyze data using ΔΔCT method^61^. ?TROUBLESHOOTING
D. Assessing downstream signaling response by IF **•** TIMING 5 d
  i. Dechorionate embryos using fine tip forceps [see Steps 46C (ii)]. Transfer embryos to 1.5 ml tubes. **▴ CRITICAL STEP** Take care not to damage the fish. If the fish is accidentally damaged, euthanize the fish, and do not use it for IF. Minimization of collateral damage mainly just takes practice.
  ii. Aspirate 10% HBSS and wash the embryos twice with 1 ml of PBS. **▴ CRITICAL STEP** Do not expose embryos to air or allow them to dry out at any point.
  iii. Aspirate PBS and add 1 ml of 4% wt/vol PFA in PBS. Gently rock tubes on their sides overnight (18 h) at 4 °C. **▴ CRITICAL STEP** Put up to 40 embryos per tube. **! CAUTION** Formaldehyde is a known carcinogen. Handle with extreme care. ▪ **PAUSE POINT** Embryos can be stored in PFA at 4 °C for at least one week. **▴ CRITICAL STEP** Handle embryos with care past this step as they are fragile after being fixed. Two general and useful techniques: (1) Always first stand tubes upright and allow embryos to fall to the bottom of the tube before replacing solution. (2) Carefully lay tubes on their sides for incubation.
  iv. Stand tubes upright and allow embryos to fall to the bottom of the tube. Aspirate PFA and carefully add 1 ml of ice-cold methanol.
  v. Gently lay tubes on their sides and place in a −20 °C freezer overnight (18 h). **▪ PAUSE POINT** Embryos can be maintained in methanol at −20°C indefinitely.
  vi. Stand tubes upright and allow embryos to fall to the bottom of the tube. Aspirate methanol and carefully add 1 ml of PDT.
  vii. After embryos have re-settled at the bottom of the tube, remove the initial PDT wash and replace with 1 ml of fresh PDT.
  viii. Carefully lay tubes on their sides and rock the tubes gently at room temperature for 30 minutes
  ix. Stand tubes upright and allow embryos to fall to the bottom of the tube. Aspirate PDT and carefully add 1 ml of IF blocking buffer. Rock gently at room temperature for 1 h.
  x. Prepare the primary antibody solution by diluting primary antibodies to the appropriate concentration in 1.2x number of samples x 700 μl of IF blocking buffer.
  xi. Stand tubes upright and allow embryos to fall to the bottom of the tube. Aspirate blocking buffer and carefully add 200 μl the primary antibody solution prepared in Step 46D(x). Remove this solution and add 500 μl the primary antibody. Rock gently for 2h at room temperature or overnight at 4 °C. **▴ CRITICAL STEP** For proteins with low expression levels or less-sensitive antibodies, incubation of embryos with the primary antibody at 4°C overnight is recommended.
  xii. Stand tubes upright and allow embryos to fall to the bottom of the tube. Aspirate antibody solution and carefully add 1 ml of PDT. Rock gently for 30 min.
  xiii. Stand tubes upright and allow embryos to fall to the bottom of the tube. Aspirate PDT and carefully add 1 ml of fresh PDT. Rock gently for 30 min.
  xiv. Stand tubes upright and allow embryos to fall to the bottom of the tube. Aspirate PDT and carefully add 1 ml of IF blocking buffer. Rock gently for 1 h. **▴ CRITICAL STEP** Perform all further steps under low light to protect fluorescent antibodies. Incubate tubes in an opaque box or cover with foil.
  xv. Prepare the secondary antibody solution by diluting the antibodies 1:1000 (or other optimized dilution factor) in 1.2 x number of samples x 700 μl of IF blocking buffer. **▴ CRITICAL STEP** For less sensitive antibodies, the secondary antibody can also be increased (1:500).
  xvi. Stand tubes upright and allow embryos to fall to the bottom of the tube. Aspirate blocking buffer and carefully add the 200 μl secondary antibody solution prepared in step 16, remove this and add 500 μl secondary antibody solution. Rock gently for 1.5 h at room temperature.
  xvii. Stand tubes upright and allow embryos to fall to the bottom of the tube. Aspirate antibody solution and carefully add 1 ml of PDT. Rock gently for 30 min.
  xviii. Stand tubes upright and allow embryos to fall to the bottom of the tube. Aspirate PDT and carefully add 1 ml of fresh PDT. Rock gently for 30 min. **▴ CRITICAL STEP** Imaging should be performed immediately following the end of the staining procedure. Store prepared samples on ice in a covered box prior to imaging
  xix. Image stained fish using a fluorescence stereomicroscope or confocal microscope according to the manufacturer’s instructions. For imaging on a stereomicroscope we use a 0.5 cm deep 2% wt/vol agarose in PBS pad. For imaging on a confocal microscope we use a glass backed imaging plate. **▴ CRITICAL STEP** Add a few drops of PBST on agarose plate to keep the fixed fish moistened if using a stereomicroscope. **▴ CRITICAL STEP** Sufficient numbers of fish or cells must be quantitated to generate reliable data. We typically aim to quantity at least 7 fish per experimental condition or hundreds of cells from at least 3 fish. ?TROUBLESHOOTING

**TABLE 1.**
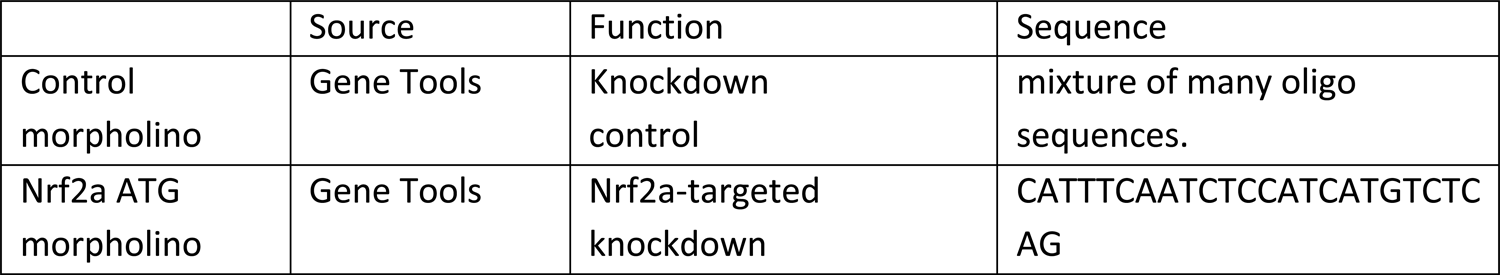
Morpholinos for gene knockdown in zebrafish.

**TABLE 2.**
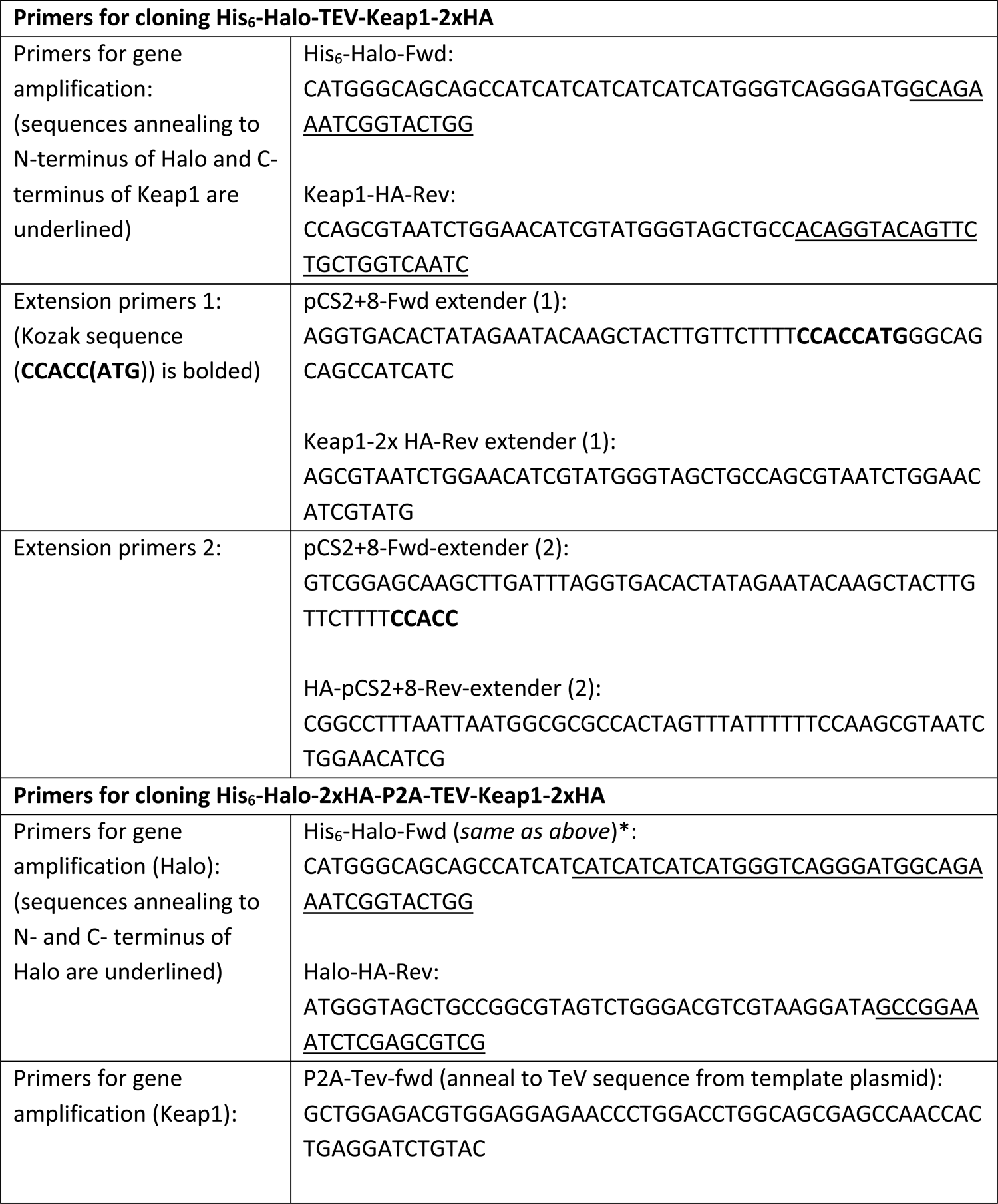
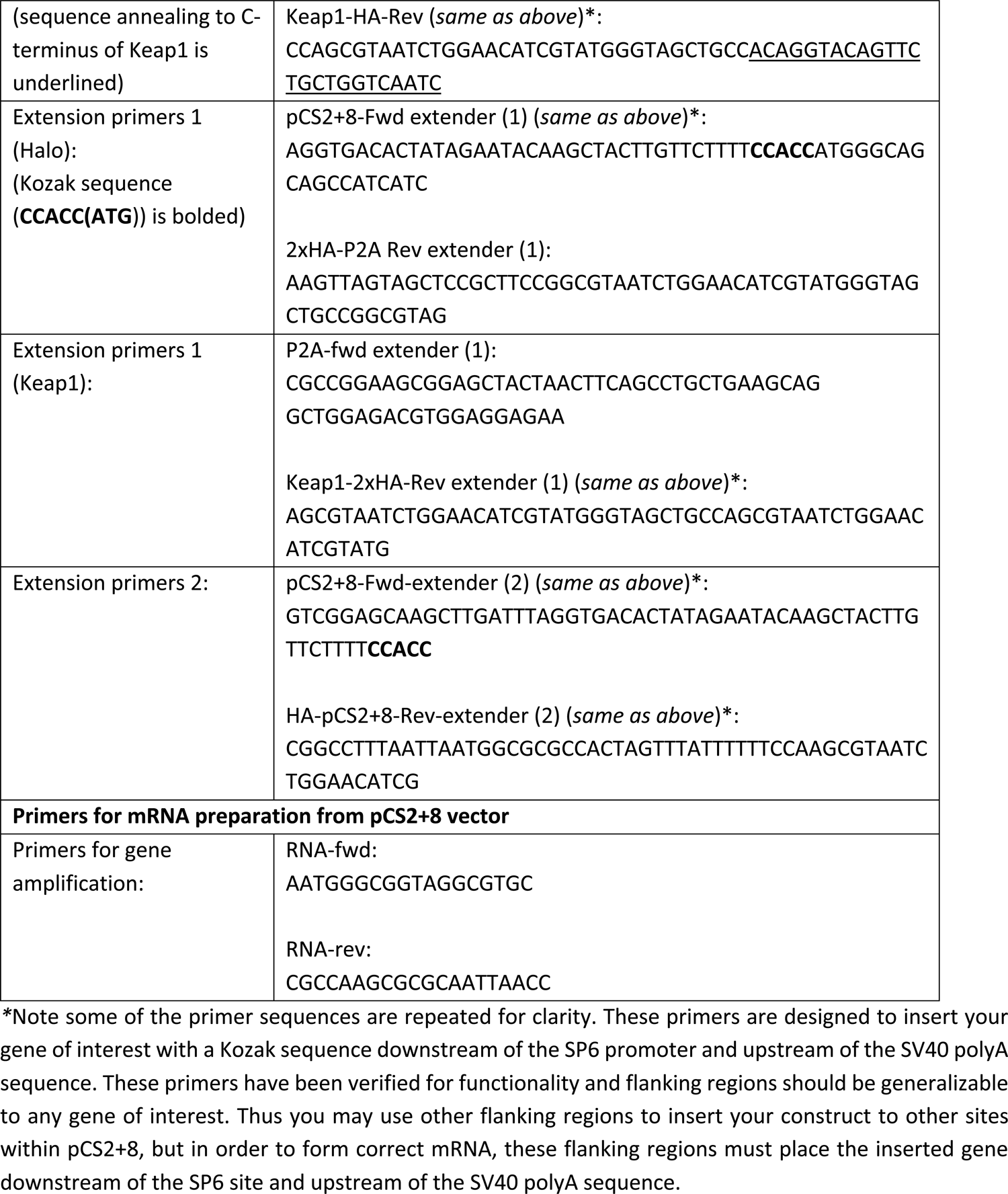
Primers for cloning His_6_-Halo-TEV-Keap1-2xHA and His_6_-Halo-2xHA-P2A-TEV-Keap1-2xHA and

**TABLE 3.**
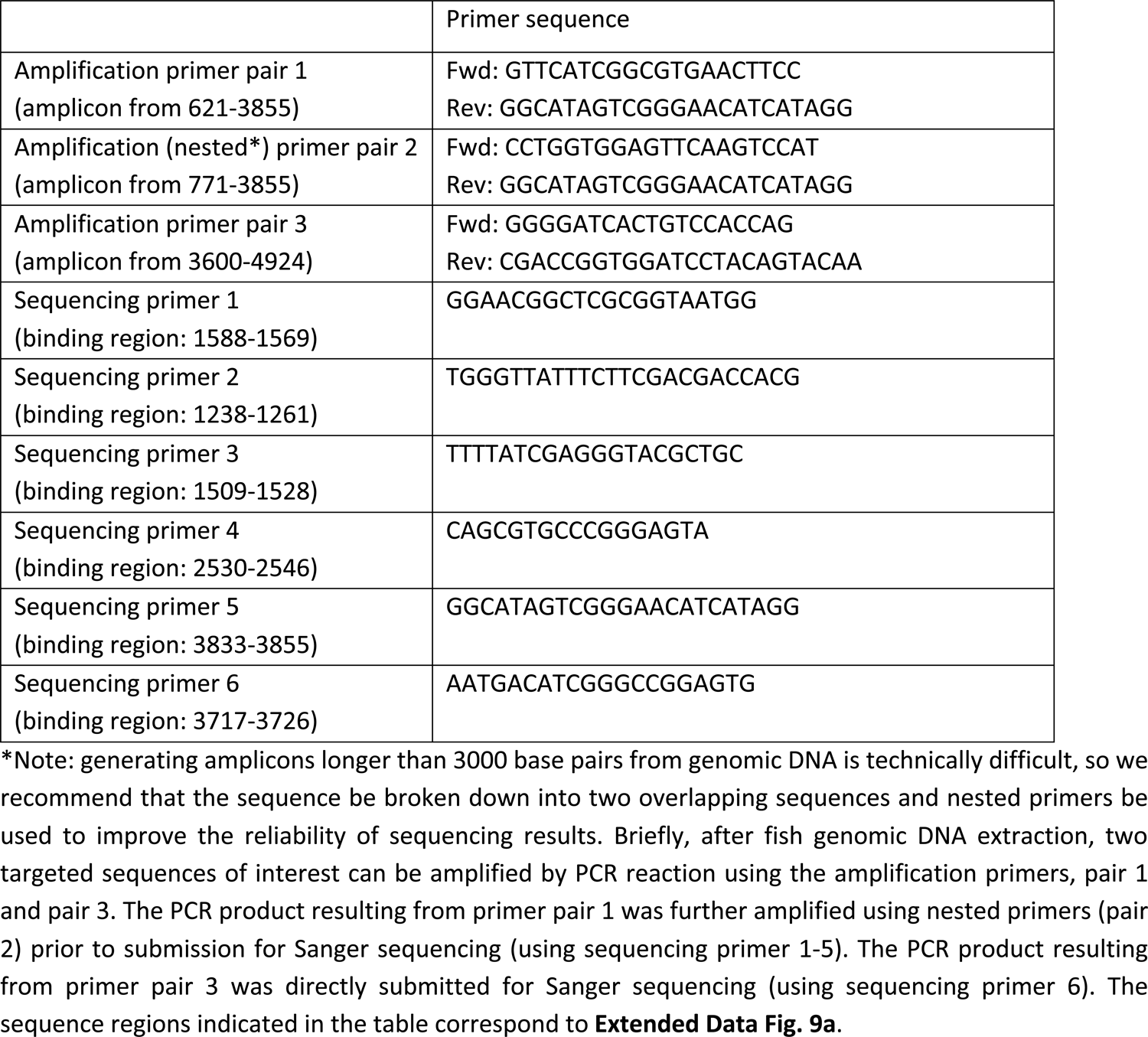
Primers for genotyping Tg(myl7:DsRed-P2A-Halo-TEV-Keap1-2xHA,cry:mRFP1).

## TROUBLESHOOTING

See Table 4 for troubleshooting guidelines.

**TABLE 4.**
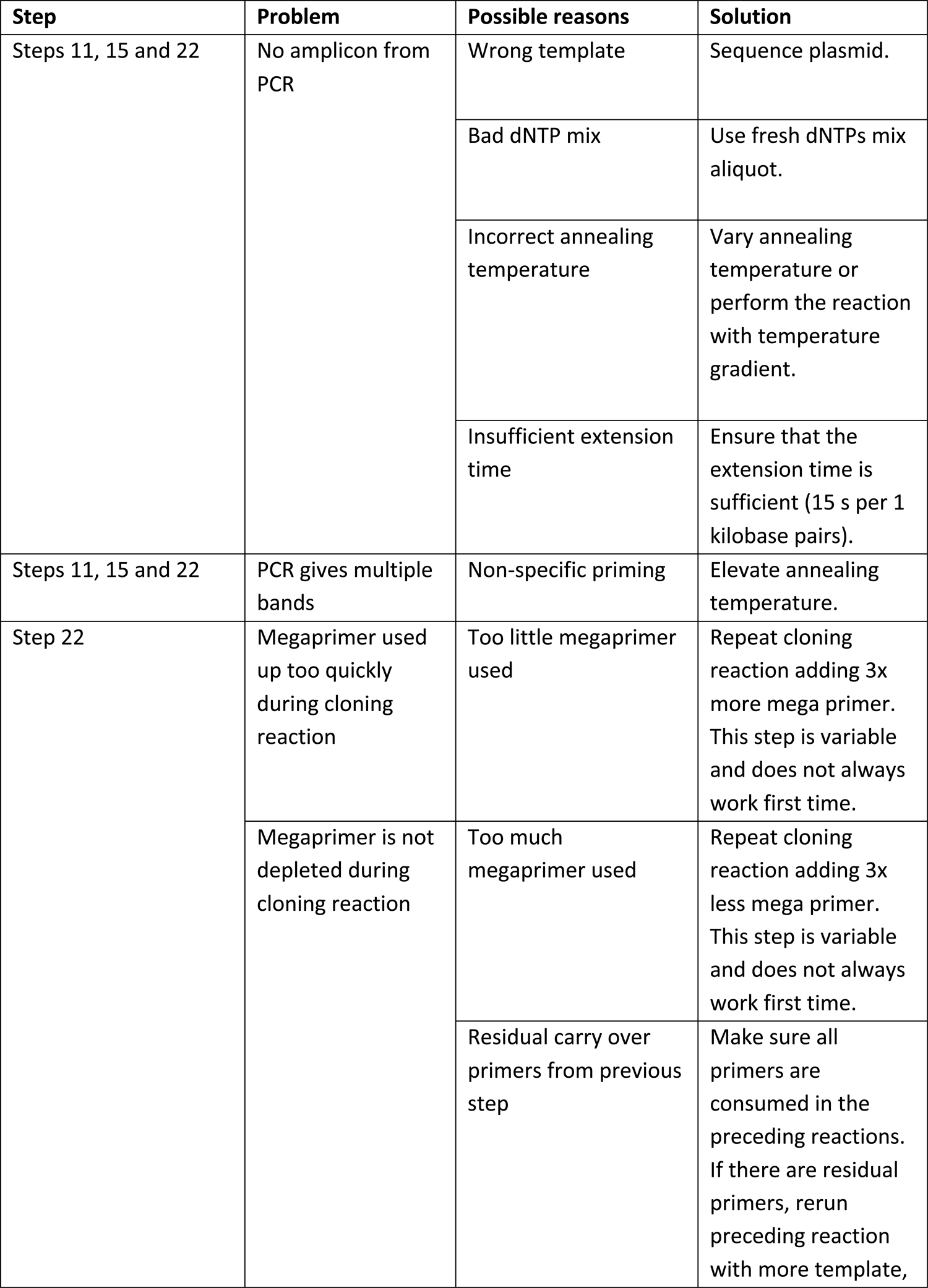
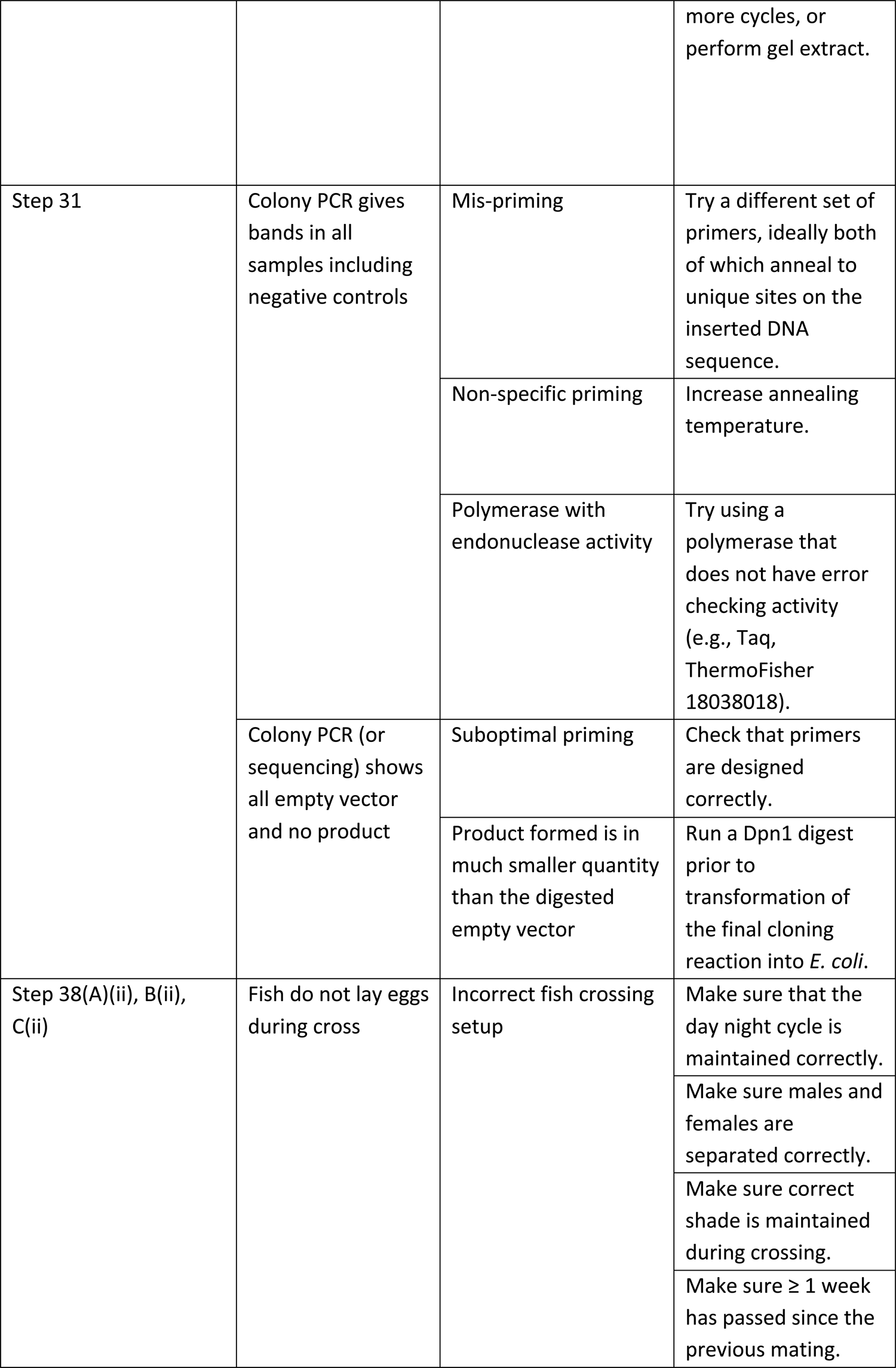
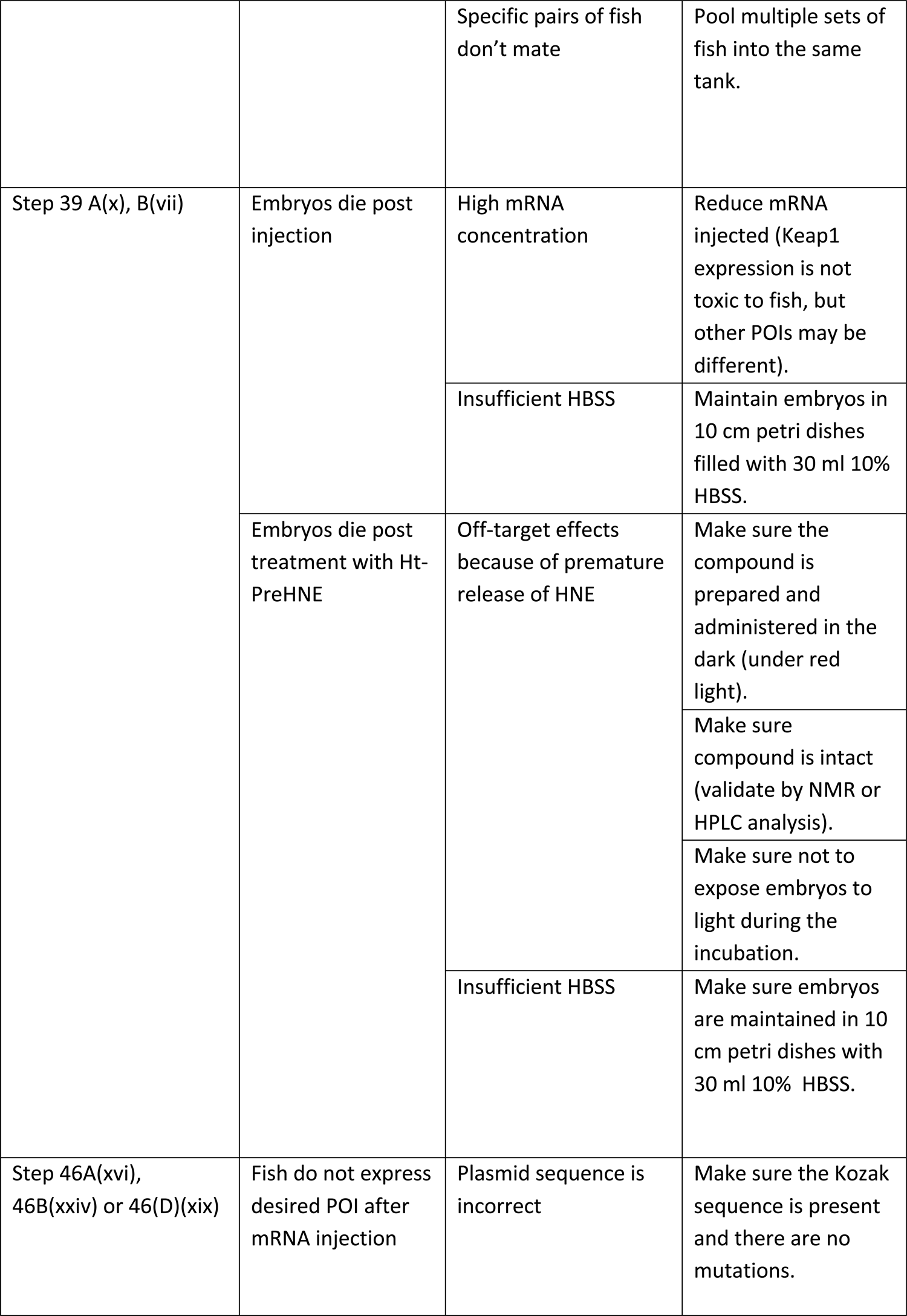
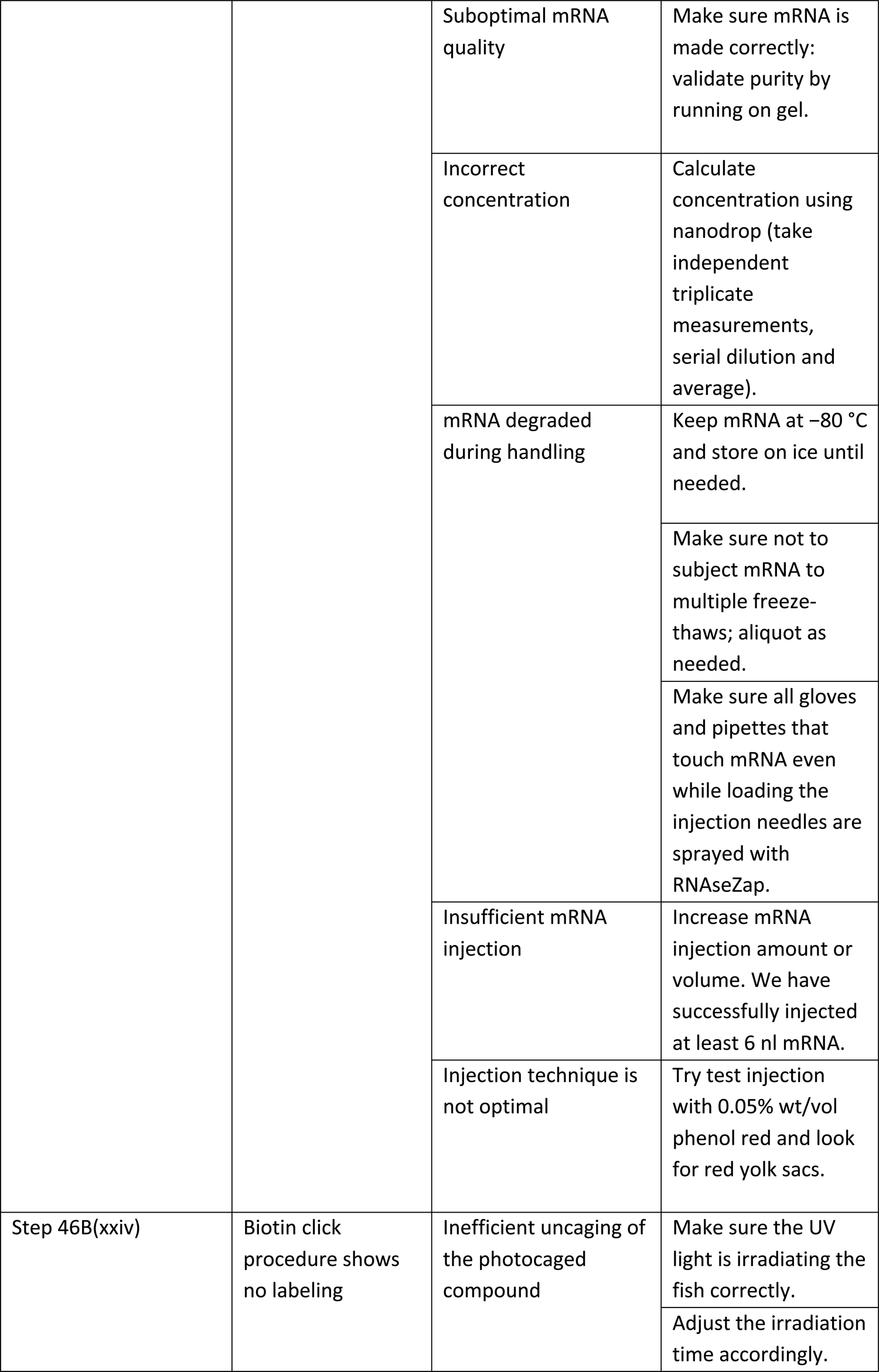
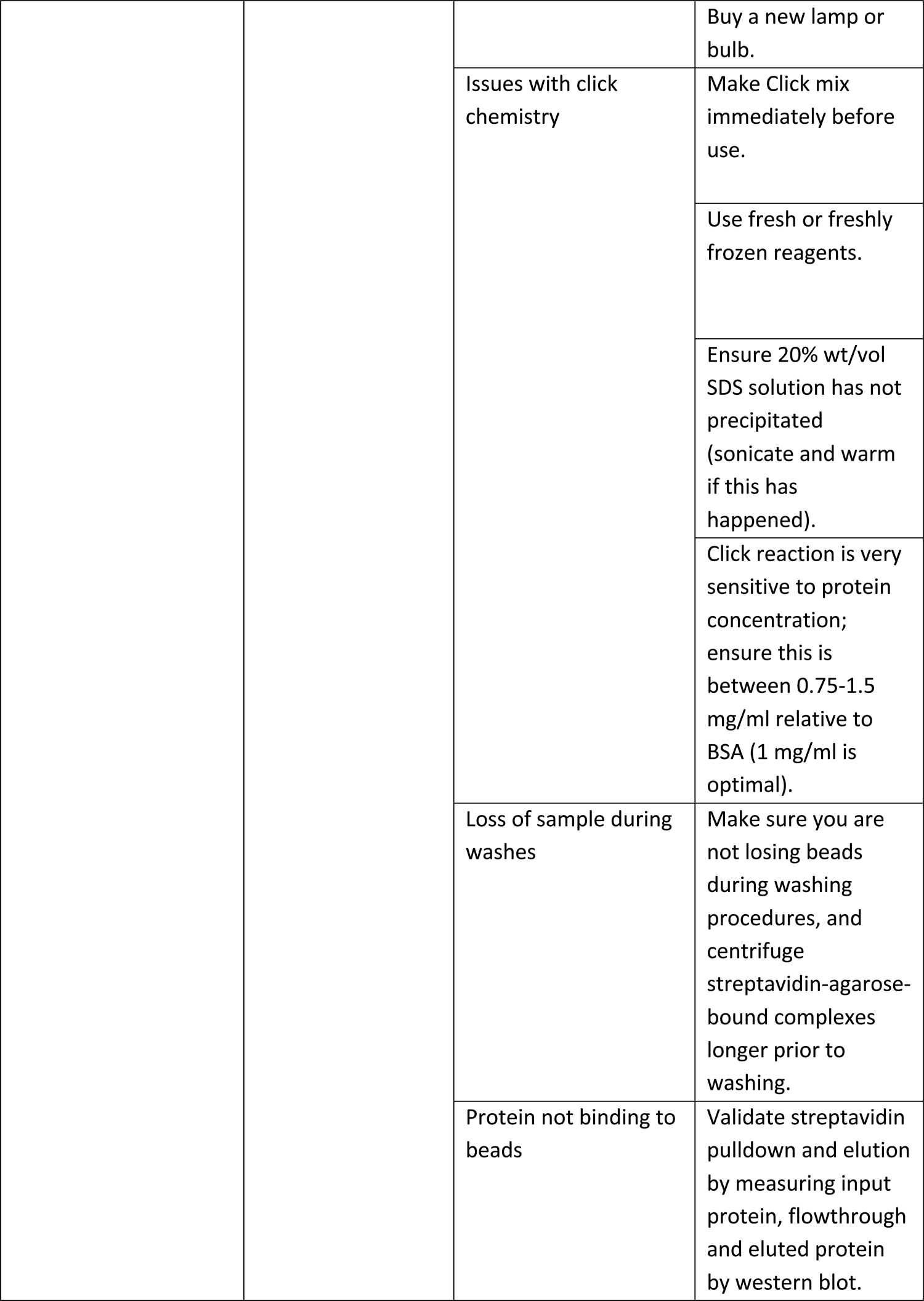
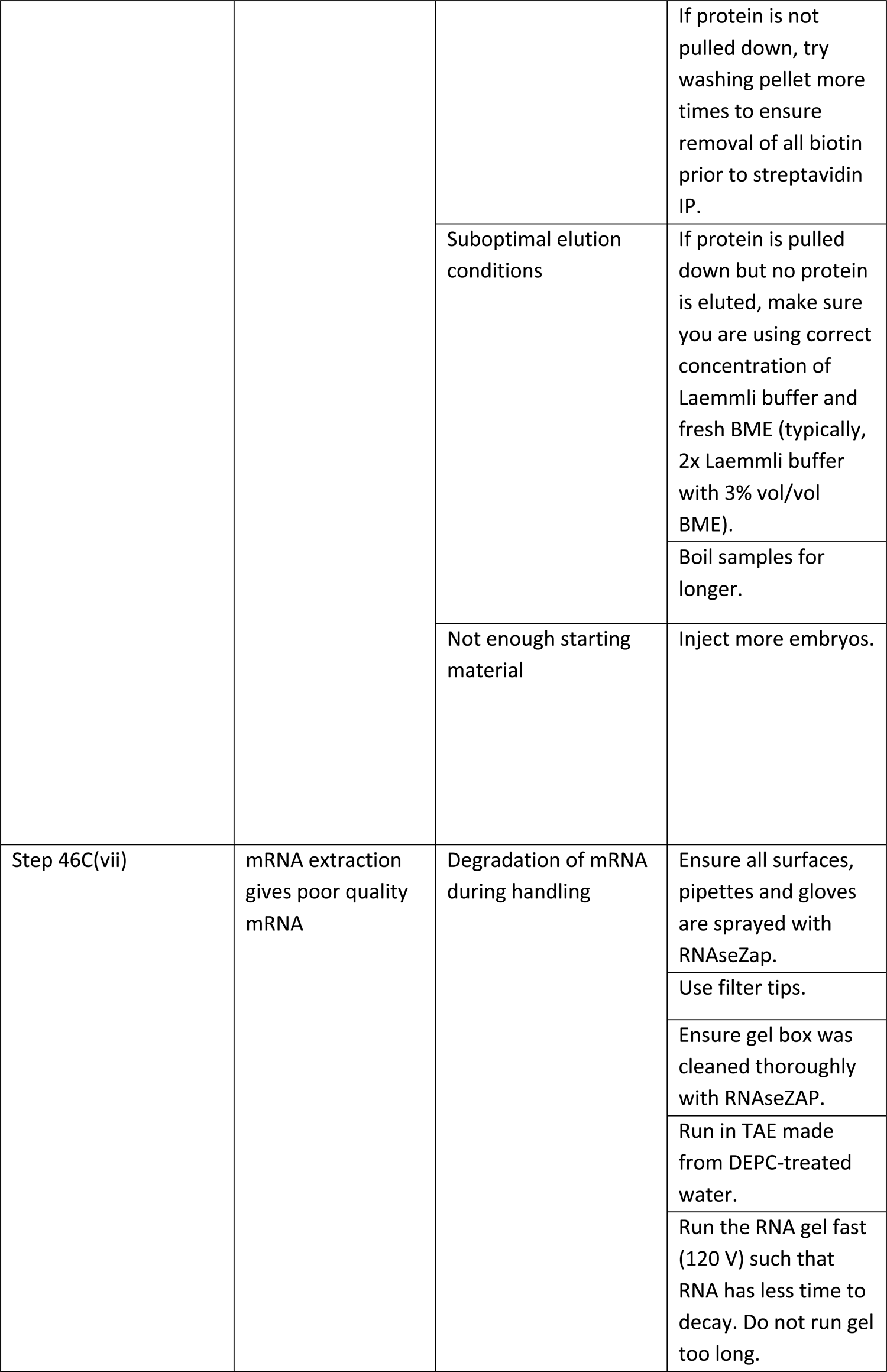
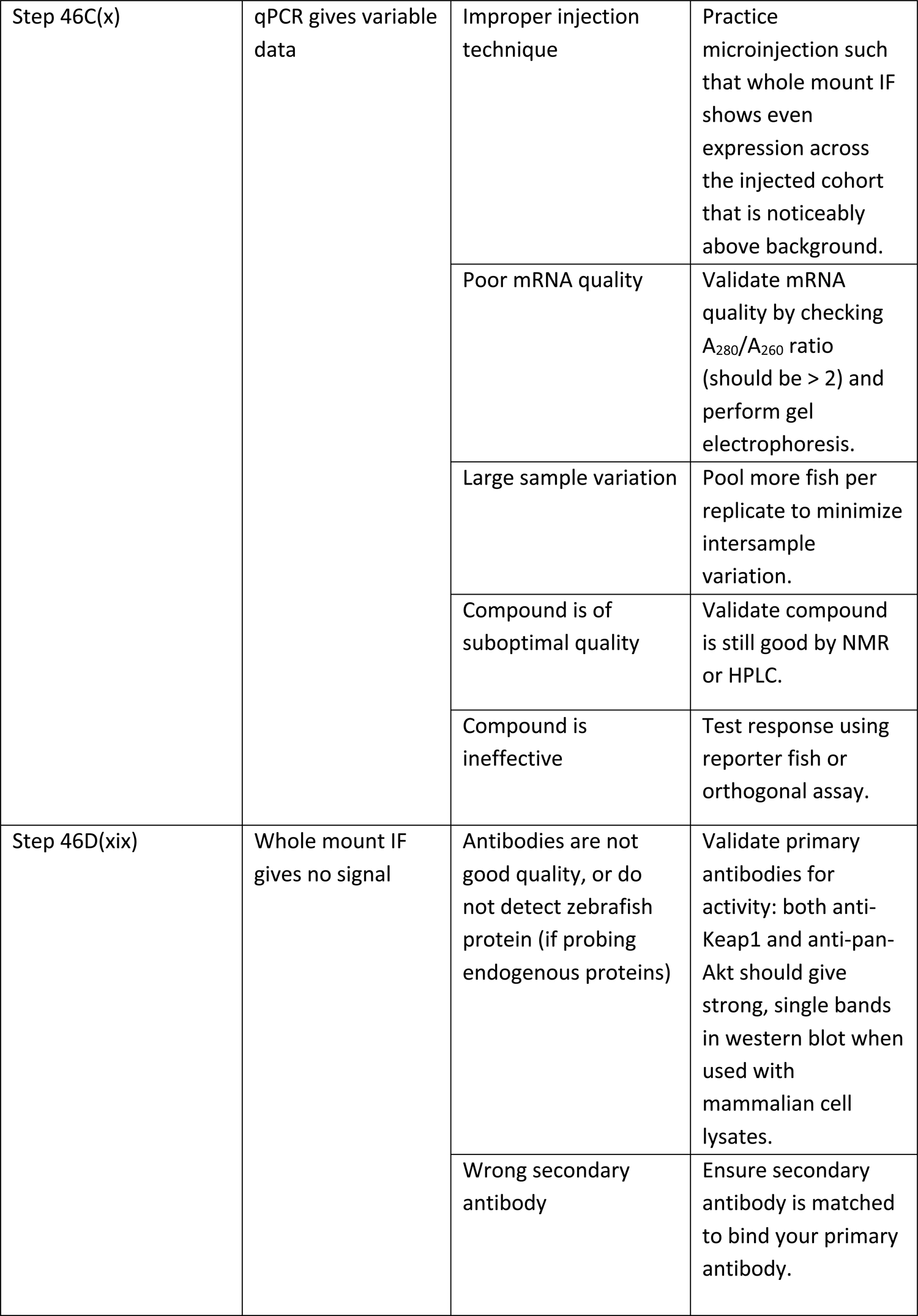
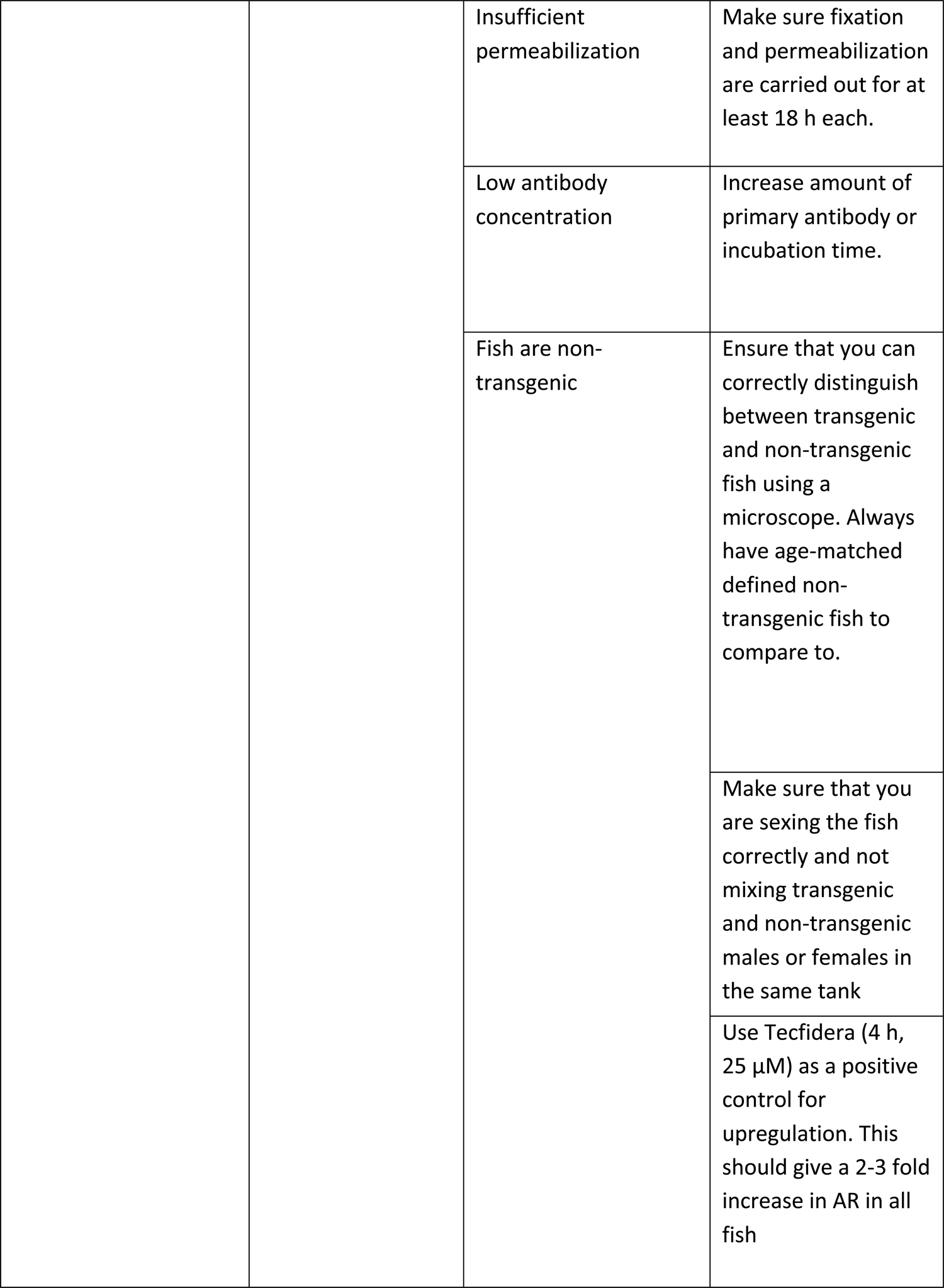
Troubleshooting table.

## Timing

Steps 1-33, setting up plasmid constructs for expression: 7 d. Steps 34-37, creating mRNA: 0.5 d.

Step 38, setting up fish crosses and obtaining fish eggs, 0.5 d (one half-evening and one half-morning) Step 39, microinjection, 3 h

Steps 40-45, Z-REX procedure, 2-8 h

Step 46, downstream analyses

A. Assessing targeting on POI by click coupling with Cy5 azide, 7 h
B. Assessing targeting on POI by click coupling with Biotin pull down, 3 d
C. Assessing downstream transcriptional response by qRT-PCR, 3 d
D. Assessing downstream signaling response by IF, 5 d

## ANTICIPATED RESULTS

Z-REX gives a target-specific mode to regulate cellular decision making in developing fish. Although an approximation to endogenous signaling modes, Z-REX none-the-less presents an innovative strategy to modulate specific pathways using a native chemical messenger targeted to a native signaling protein. Thus, Z-REX allows a specific, chemically-defined state to be interrogated in a native environment, which is otherwise inaccessible. Z-REX offers benefits to inducible gene upregulation: rapid and robust downstream responses under otherwise largely unperturbed endogenous conditions. Z-REX also offers benefits relative to bolus dosing methods in terms of low off-target effects and low toxicity. For instance, knockdown or knockout of identified targets of Tecfidera have not severely impacted well-known phenotypes observed upon Tecfidera treatment^20, 62^. We thus anticipate that Z-REX will be a go-to tool for those aiming to study effects of redox signaling on development, preconditioned responses, and other behavioral mechanisms. Z-REX, furthermore, allows troubleshooting at each specific step of the process.

### Validation of expression

We have deployed several experiments to validate and quantify relative expression the constructs we have used, in this case: Halo-TEV-Keap1-2xHA (“Halo-Keap1” hereafter) and Halo-2xHA-P2A-TEV-Keap1-2xHA (“Halo-P2A-Keap1” hereafter) (note: these constructs express human Keap1).

To validate specificity of Keap1 primary antibody (which can detect ectopically expressed human Keap1 and both isoforms of zebrafish Keap1), compare samples stained with and without primary antibody. Only the samples stained with primary antibody show strong fluorescent signal. (Extended Data Fig. 1a). These analyses validate that Keap1 primary antibody detects ubiquitous protein expression relative to the no primary antibody control. The relative ectopic expression of either Halo-Keap1 or Halo-P2A-Keap1 is compared to endogenous zebrafish Keap1 (Fig. 3a **and Extended Data** Fig. 1b and c). The two Keap1 constructs introduced by mRNA injection show around 2-fold higher anti-Keap1-specific signal than the non-mRNA injected samples (Fig. 3a–b). Although signals cannot be directly compared due to using human Keap1 in the ectopic constructs, given that this protein has high similarity to both zebrafish Keap1 paralogs, a conclusion is made that expression of Halo-Keap1 and Halo-P2A-Keap1 is similar and that both are expressed at a similar level to endogenous Keap1. We have reached a similar conclusion with another Halo-tagged protein in fish previously.

Relative to the non-injected fish, both constructs express HA-tagged proteins. Relative expression (by IF) is consistent across fish: Halo-P2A-Keap1 gives marginally more HA signal (having 2-fold greater total number of HA-tags) than Halo-Keap1 (Extended Data Fig. 1b–d). This observation indicates that expression levels of Halo-POI and Halo-P2A-POI are similar. A similar conclusion can be drawn from western blot (Fig. 3d).

### Validation of POI-electrophile targeting in fish

Cy5 click assay and biotin pulldown can be used to validate POI-electrophile targeting and examine whether the POI is a sensor of the electrophile delivered. In both assays, TEV protease is used to cleave fusion protein post Z-REX (Halo-TeV-POI mRNA is used). The TEV protease treatment is critical to differentiate whether the electrophile is bound to Halo or the POI. Although the probe uncaging efficiency is high (t_1/2_<1 min), some residual probe may remain on Halo after light exposure (protein band 3 in Fig. 3C). In our biotin pulldown assay, TeV treatment ensures the Keap1 pulldown can only happen when the electrophile is bound to Keap1 (but not the Halo in fusion protein construct). Note that, in both assays, (1) Keap1 protein band results from “photo-released” electrophile labeling on Keap1 protein;(2) Halo protein band results from “uncaged” probe binding to Halo protein; (3) non-cleaved Halo-Keap1 band can be results from either cases described above if TEV protease treatment is not applied.

In our depiction in Fig. 3C, we describe how to calculate delivery efficiency or occupancy of HNE on the POI. These two variables are important to assess how effective the total protein targeted is at sensing the electrophile in question. In the experiment shown in Fig. 3d, non-mRNA-injected fish; fish injected with HALO-TEV-KEAP1 or HALO-P2A-Keap1 were used. The two different mRNA-injected fish were either untreated or subjected to Z-REX conditions. Post harvest, dechorionation, Click coupling, as described above, Keap1 protein was shown to be labeled by HNE (i.e., enriched after streptavidin pulldown, only in the Halo-TeV-Keap1 samples subjected to Z-REX). Thus, HNE released during Z-REX is unable to label Keap1 if the Halo domain is no longer fused to Keap1 (see input gel). Successful replication of these data on any POI validates the P2A system as a useful control for adventitious HNE and other potential reactive species generated during Z-REX.

Critically, post Z-REX, delivery of the electrophile (e.g., HNE) only occurs when using Halo-(TEV)-Keap1, but does not occur when using the corresponding non-fused analog, Halo-P2A-Keap1 (Fig. 3d). This result proves that Z-REX proceeds by *pseudo-intramolecular delivery* from Halo to Keap1. Thus Z-REX provides a surgical tool to perturb a specific (intended target) protein and further allows an ideal negative control to account for affects due to off-target HNEylation. Control groups for the two assays above are demonstrated in Fig. 3c and d. The targeting efficiency and ligand occupancy calculation can be done by using the equation in Fig. 3c.

### Assessment of downstream signaling using a transgenic reporter system

In fish expressing Halo-Keap1, fluorescence in the tail is relatively low in unstimulated fish (Figs. 4a and c), but upon Z-REX, global dosing with Tecfidera (25 µM, known to elicit phenotypic effects at this concentration in cultured cells^62^ and zebrafish^26^) or dosing with lipid-derived electrophile e.g., HNE (25 µM), significant and selective AR upregulation in the tail can be seen [Figs. 2e, 4a and c (tail), and Extended Data Fig. 2a (head)]. Importantly, upon Z-REX, fish expressing Halo-P2A-Keap1 show similar basal levels of AR, but do not upregulate AR upon Z-REX in the tail (Fig. 4b–c). Of note, Tecfidera (25 µM) treatment elicits a similar response in Halo-P2A-Keap1-expressing fish to those expressing Halo-Keap1 (Fig. 4c). Additionally, this fluorescence is decreased upon Halo-Keap1 expression, but not upon Halo expression (Fig. 2c–d). This is consistent with the role of Keap1 as a suppressor of Nrf2-driven AR under basal conditions. This result validates the reporter and the function of the constructs (Fig. 2a).

Given that expression of Halo-Keap1 and Halo-P2A-Keap1 are similar (Fig. 3), and Keap1-specific delivery only occurs to Halo-Keap1 (Fig. 3d), the last result (Fig. 2 and 4, and Extended Data Fig. 2a) provides strong evidence that downstream responses measured are due to Keap1-modification, not due to any other effect associated with the probe, or the minimal amount of HNE released to the system that does not target Keap1. However, Z-REX and bolus dosing for the most part do not show significant AR upregulation in the head (Extended Data Fig. 2a). The differences between response in head and tail could be ascribed to the higher basal AR in the head (resisting further AR), the increased depth in tissue affecting photochemistry or sensitivity of the measurements, or possibly due to other regulatory mechanisms that resist AR changes downstream of Keap1 specific to these tissues.

Consistent with imaging of *Tg(gstp1:GFP)* reporter fish (*vide supra*), significant increases in several AR-driven genes by qRT-PCR in the tail upon Z-REX can be seen (Fig. 4d). Elevation of AR upon Z-REX also requires that Halo and Keap1 be fused, reaffirming that our phenotypes are Keap1-specific. There is also a change in mRNA levels of AR-associated genes in the head that are dependent upon Keap1 and Halo being fused (and Extended Data Fig. 2b). We thus suggest independently validating response observed in transgenic fluorescent reporter fish by qRT-PCR.

Moreover, tissue- or cell-type-specific signaling induction can be achieved using transgenic embryos expressing Halo-Keap1 in a certain tissue/cell-type, such as *Tg(gstp1:GFP;myl7:DsRed-P2A-Halo-TEV-Keap1-2xHA,cry:mRFP1)* embryos, expressing Halo-Keap1 in cardiomyocytes. Upon Z-REX, selective increase of AR can be seen in the heart: other tissues examined do not show a change in fluorescence in any condition studied (Fig. 5 and Extended Data Fig. 4).

## DATA AVAILABILITY STATEMENT

Plasmids and strains generated and reported in this protocol are available upon request. Uncropped blots for Fig. 3 are provided in **Source Data 1**. Figs. 2c and e, **3b**, **4c**-**d**, **5b** and Extended Data Figs. 1d, 2a-**b**, **4b** and **5b** have associated **Source Data 2**.

## ACKNOWLEDGMENTS.

Dr. Guillaume Valentin, Florian Lang, et al. from the EPFL zebrafish husbandry and microinjection/imaging facility [license no. VD-H230]; Nikki Gilbert from the Fetcho laboratory for assistance with fish husbandry and breeding; Professor Jin Zhang (University of California, San Diego and Johns Hopkins University) for providing AktAR reporter plasmid; Prof. Iain MacPherson for the ligase-free cloning technology; ZeClinics for assistance with transgenesis; the Cornell University zebrafish husbandry and microinjection/imaging facility [NIH R01 NS026593, J. Fetcho]; Novartis FreeNovation Award (Y.A.); European Research Council (ERC-2021-CoG to Y.A.); Swiss Federal Institute of Technology Lausanne (EPFL) (Aye Laboratory); NIH CBI training grant [NIH T32GM008500 (J.R.P. as a trainee fellow)] and AHA predoctoral fellowship (17PRE33670395 to J.R.P.); HHMI International Fellow (S.P.).

## AUTHOR CONTRIBUTIONS

K.-T.H., J.R.P, M.J.C.L., S.P, and S.R. collected the data. J.R.F. helped with experimental design and zebrafish methodologies. K.-T. H., J.R.P., M.J.C.L., S.P., and Y.A. analyzed the data.

B.M. instructed microinjection, assisted with fish collection, and consulted on early design of experiments.

Y.A. oversaw project supervision and funding acquisition. All authors contributed to the development of the protocols and co-wrote the paper.

## EXTENDED FIGURE LEGENDS

**Extended Data Figure 1.**
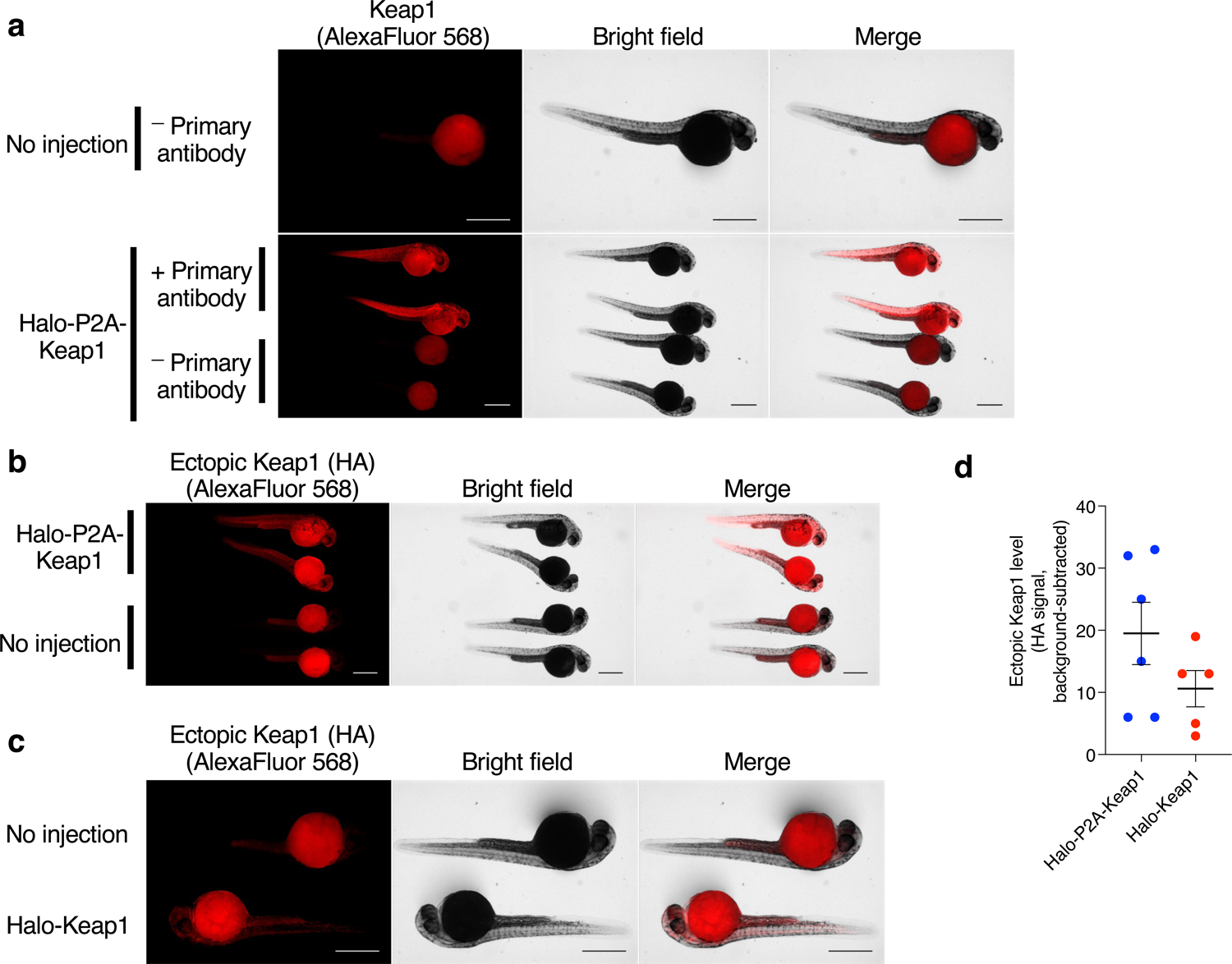
*mRNA injection of Halo-POI constructs gives uniform expression of POI in fish*. Keap1 is used as a representative POI^25^. **(a)** IF analysis of zebrafish embryos (34 hpf). **Top row:** non-injected embryos not stained with primary antibody; **bottom row:** Halo-P2A-Keap1 injected embryos [**note:** there are two sets of each fish in this row, corresponding to fish either stained (top set) or not stained (bottom set) with anti-Keap1 primary antibody]. **(b)** IF analysis of zebrafish embryos (34 hpf) injected with Halo-P2A-Keap (top), or non-injected control embryos (bottom). Both groups were stained with anti-HA primary antibody as described (**note:** all embryos in both groups were treated with the same primary and secondary antibody mix). (**c**) Same as (b) but no injection was compared to Halo-Keap1 injection. (**d**) Quantitation of data in (b) and (c). Halo-P2A-Keap1: n=6, SEM=5.012; Halo-Keap1: n=5, SEM=2.926. Scale bar, 500 µm in all images.

**Extended Data Figure 2.**
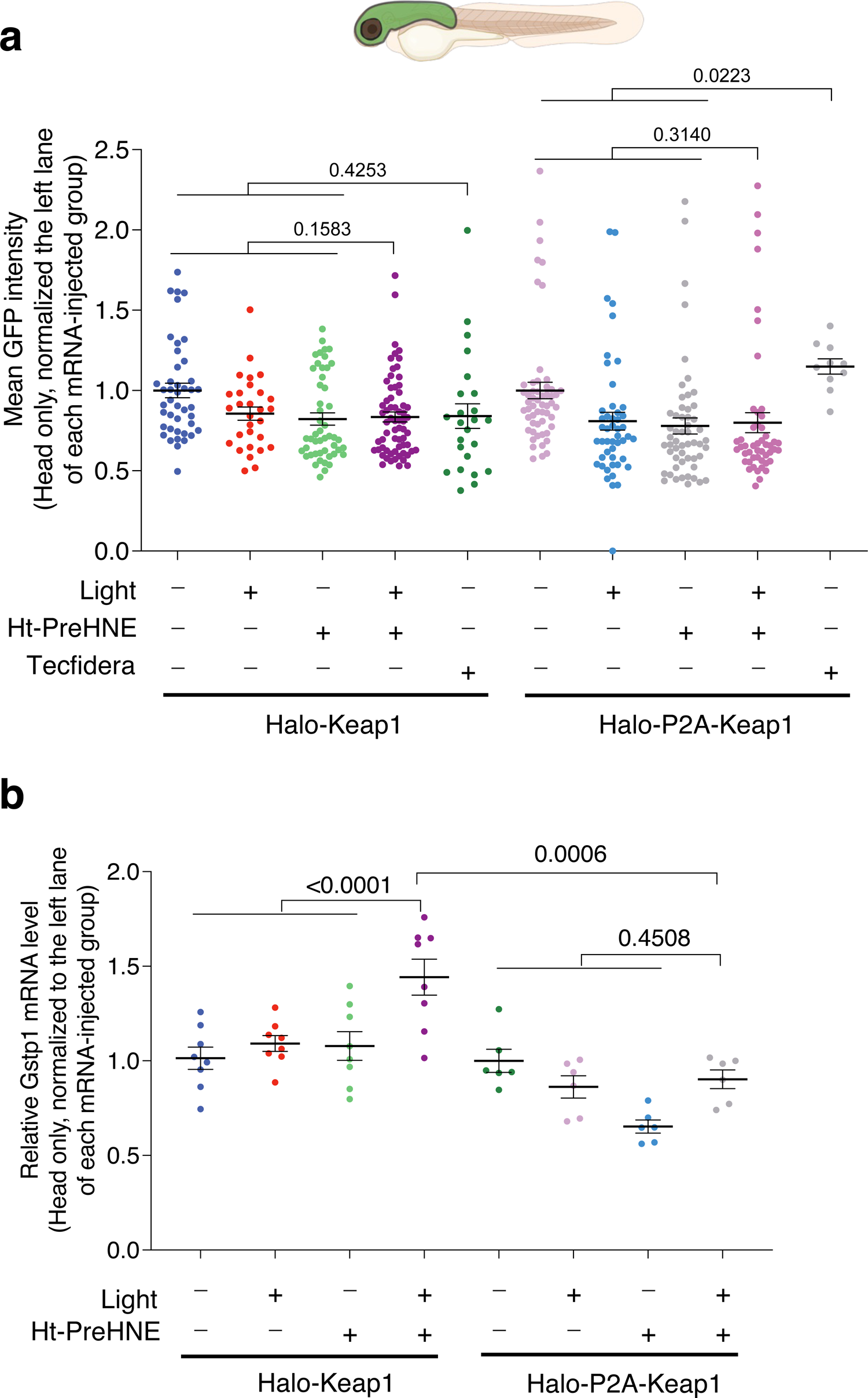
*Downstream pathway activation analyzed by transgenic reporter fish and qRT-PCR analysis.* In this example, responsivity differences were characterized for Keap1–Nrf2–AR^25^ pathway using *Tg(gstp1:GFP)* fish and an endogenous downstream gene *Gstp1* driven by Nrf2/AR. (**a**) Unlike in the fin, (see also Fig. 4), Z-REX-assisted Keap1-HNEylation or whole-animal treatment with Tecfidera (25 µM, 4 h treatment) do not cause elevation of AR in the head when measured using *Tg(gstp1:GFP)*. (34 hpf) Halo-Keap1 mRNA-injected (from left to right): n=43, SEM=0.0455; n=29, SEM=0.0416; n=49, SEM=0.0378; n=65, SEM=0.0313; n=24, SEM=0.0767. Halo-P2A-Keap1 mRNA-injected (from left to right): n=55, SEM=0.0510; n=48, SEM=0.0553; n=54, SEM=0.0497; n=49, SEM=0.0622; n=10, SEM=0.0480. **(b)** qRT-PCR analysis is able to detect a small increase in AR in the head upon Z-REX-assisted Keap1-HNEylation, that is selective to the Halo-Keap1 construct over the Halo-P2A-Keap1 construct. This is significantly less than what is observed in the tail (see Fig. 4d). Fish age: 32 hpf. Fish age: 32 hpf. Halo-Keap1 mRNA-injected (from left to right): n=8, SEM=0.0590; n=8, SEM=0.0418; n=8, SEM=0.0756; n=8, SEM=0.0948. Halo-P2A-Keap1 mRNA-injected (from left to right): n=6, SEM=0.0609; n=6, SEM=0.0586; n=6, SEM=0.0349; n=6, SEM=0.0493. P values were calculated with two-tailed unpaired Student’s t test.

**Extended Data Figure 3.**
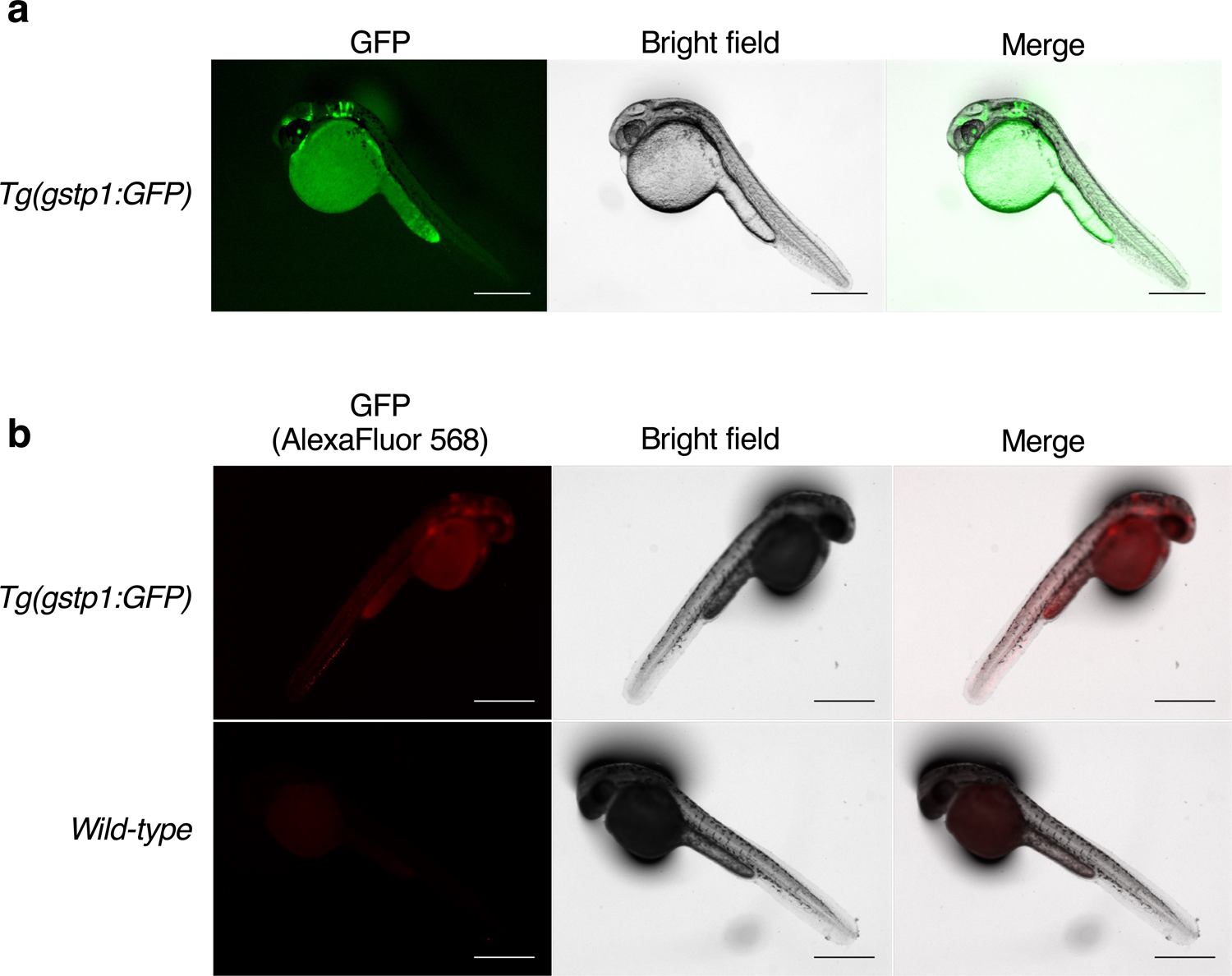
*Validation of the same outcome between whole-mount immunofluorescence and live imaging.* (**a**) Live *Tg(gstp1:GFP)* embryos (34 hpf) were dechorionated and placed on an agarose pad and imaged using GFP fluorescence (Ex. 495 nm, Em. 500-500 nm) and bright field. (**b**) Top row: *Tg(gstp1:GFP)* (34 hpf) were exposed to Z-REX conditions using Keap1-HNEylation as a representative example, and at 4-h post light exposure, dechorionated, fixed and immunostained for GFP as described. AlexaFluor568 channel shows GFP signal. GFP localization is similar for GFP intrinsic fluorescence [in (a)] and red fluorescence from IF (this figure). Bottom row: identical series of steps carried out as in top row except WT fish was used in place of *Tg(gstp1:GFP)* reporter fish. Scale bar, 500 µm in all images.

**Extended Data Figure 4.**
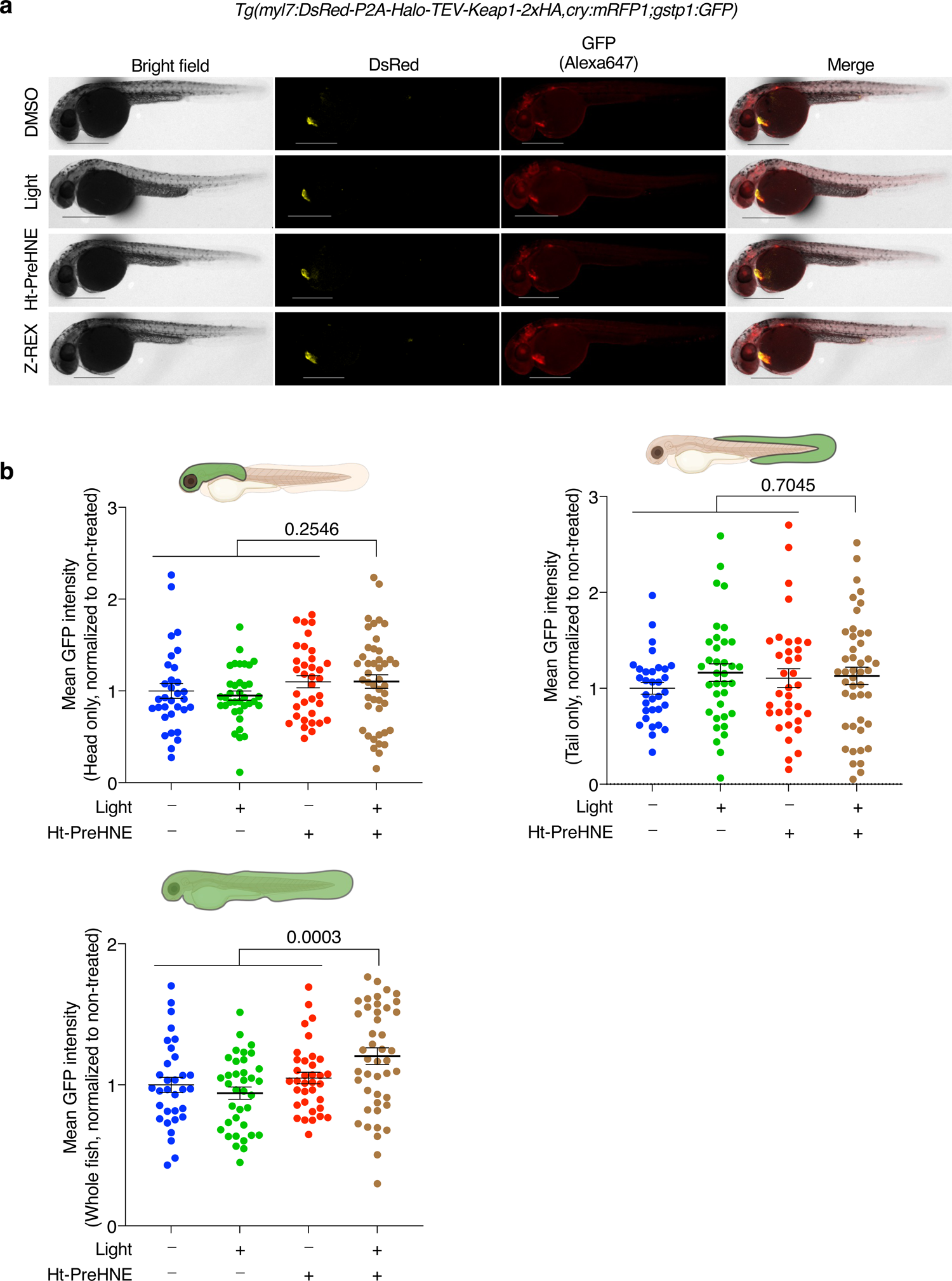
Z-REX selectively upregulates the AR in fish cardiomyocytes, but not other tissues. (**a**) Representative images of *Tg*(*myl7:DsRed-P2A-Halo-TEV-Keap1-2xHA,cry:mRFP1,gstp1:GFP*) fish (34 hpf) treated with DMSO, light, 0.3 μM Ht-PreHNE, or Z-REX (with 0.3 μM Ht-PreHNE). Scale bar, 500 µm in all images. See Fig. 5a for magnified images. (**b**) Quantification of mean GFP intensity. The quantification strategy is described in the discussion. Briefly, the head, tail (median fin fold) and whole fish are defined based on bright-field images. Image-J (NIH) quantification shows AR levels in the head, tail or whole fish were not changed (against all the control conditions) upon cardiomyocyte-specific Z-REX treatment (Fig. 5b). Fish age: 34 hpf. Sample size is the same for three plots (from left to right): n=32, n=36, n=35, n=45. SEM: head (from left to right): 0.0804, 0.0505, 0.0660, 0.0737; tail (from left to right): 0.0616, 0.0913, 0.0989, 0.0900; whole fish (from left to right): 0.0536, 0.0438, 0.0412, 0.0589. P values were calculated with two-tailed unpaired Student’s t test.

**Extended Data Figure 5.**
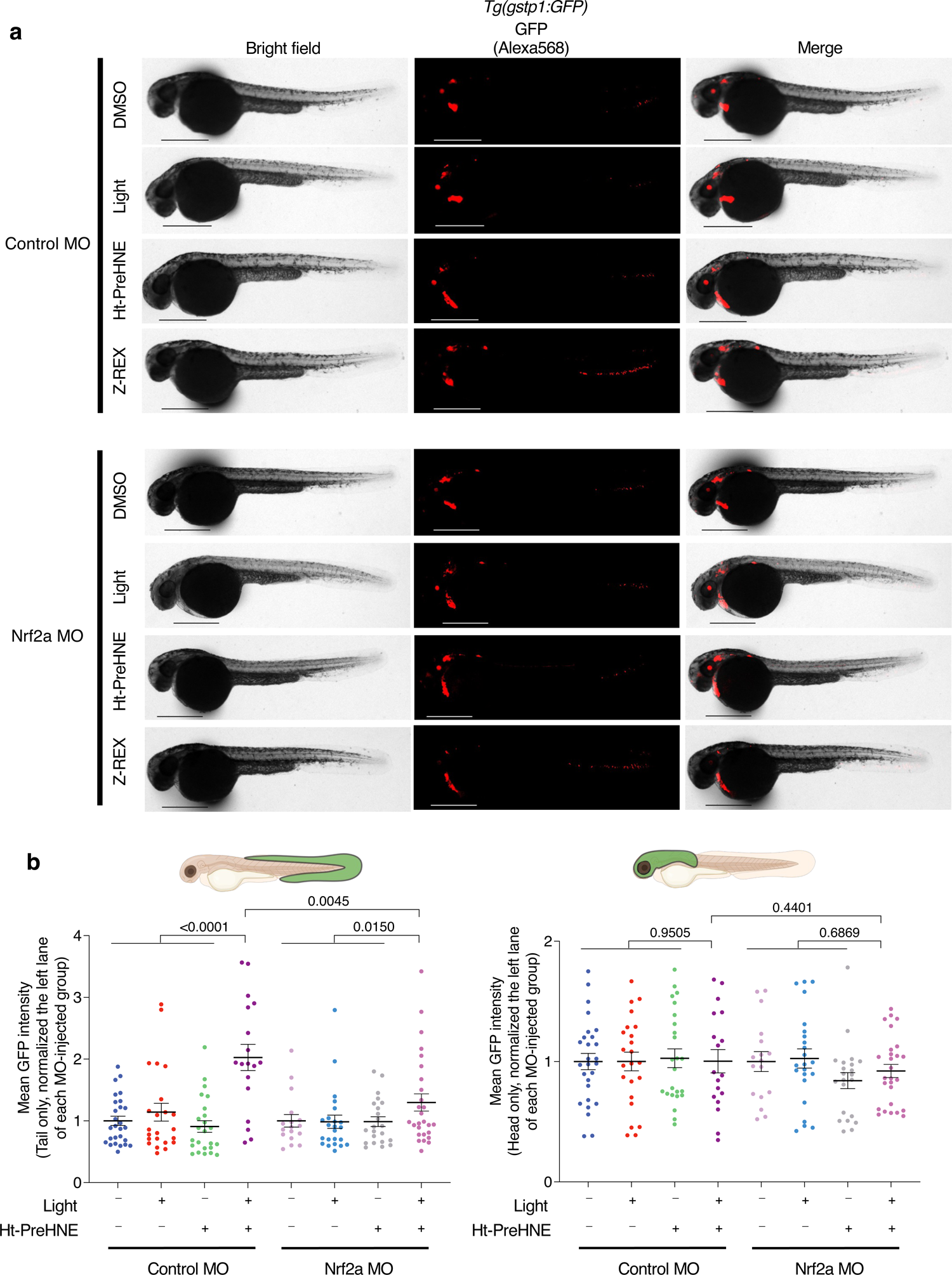
*Z-REX-mediated AR stimulation is Nrf2a-dependent.* (**a**) Representative IF-images of *Tg*(*gstp1:GFP*) fish (34 hpf) injected with 2 nl of 500 ng/μL Halo-(TEV)-Keap1-2xHA mRNA and 0.5 mM morpholino (control MO or Nrf2a MO), and treated with DMSO, light, 1 μM Ht-PreHNE, or Z-REX (with 1 μM Ht-PreHNE). Scale bar, 500 µm in all images. (**b**) Quantification of mean GFP intensity. The quantification strategy is described in the discussion. Briefly, the head and tail (median fin fold) are defined based on bright-field images. After Z-REX-treatment, control morpholino-injected fish show two-fold higher AR signal than other negative control groups (DMSO, light or Ht-PreHNE alone), whereas the Nrf2a MO-injected fish show only 1.3-fold higher AR signal, compared to corresponding negative control groups. The results demonstrate that Nrf2a is a necessary mediator in Z-REX-stimulated AR pathway. Fish age: 34 hpf. Sample size is the same for two plots (from left to right): n=27, n=23, n=24, n=18, n=17, n=23, n=22, n=27. SEM: tail (from left to right): 0.0749, 0.1438, 0.0941, 0.2122, 0.1021, 0.1074, 0.0791, 0.1394; head (from left to right): 0.0675, 0.0779, 0.0783, 0.0971, 0.0826, 0.0797, 0.0667, 0.0548. P values were calculated with two-tailed unpaired Student’s t test.

**Extended Data Figure 6.**
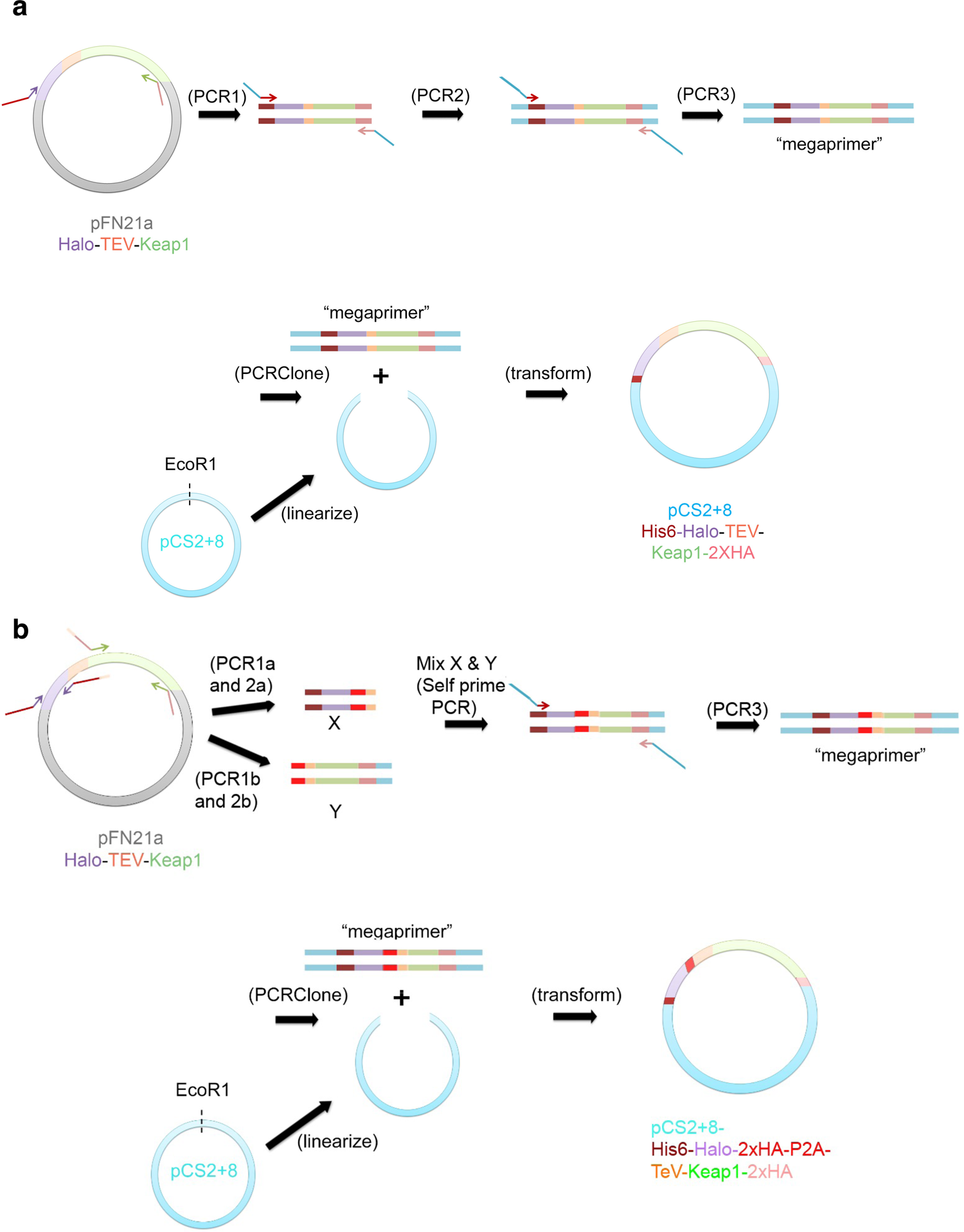
*A rapid method to generate HaloTagged-POI constructs in pCS2+8.* (**a**) Halo-Keap1 (or any desired HaloTagged POI) is amplified from the parent [pFN21a, if using Kazusa library (Promega)] using the primers stated in Table 2 (PCR1). Two subsequent PCRs, PCR2 and 3, generate megaprimer that contains the complete desired gene, tags, Kozak sequence, and flanking regions (blue) that anneal to the pCS2+8 plasmid downstream of the SP6 promoter and upstream of the SV40 poly-A tail. This megaprimer is used to prime a PCR reaction (PCRClone) with the linearized pCS2+8 and the crude mixture (with or without Dpn1 digestion) is directly transformed into *E coli*. (**b**). Halo and the POI (in this case Keap1) are amplified (PCR1a and 1b) *separately* by PCR from the original plasmid and extended (PCR2a and 2b). Primers are designed such that the 3’-end of the Halo amplicon (X) can overlap with the 5’-end of the Keap1 amplicon (Y). These two ends encode the linker region between Halo and Keap1 in the final construct. X and Y are used in a self-priming reaction to make the fused DNA (self prime PCR), that is then amplified by PCR using primers that will introduce 5’- and 3’ ends that anneal to pCS2+8 in the same position as in (a) (PCR3). This megaprimer is then used as in (a) (PCRClone).

**Extended Data Figure 7.**
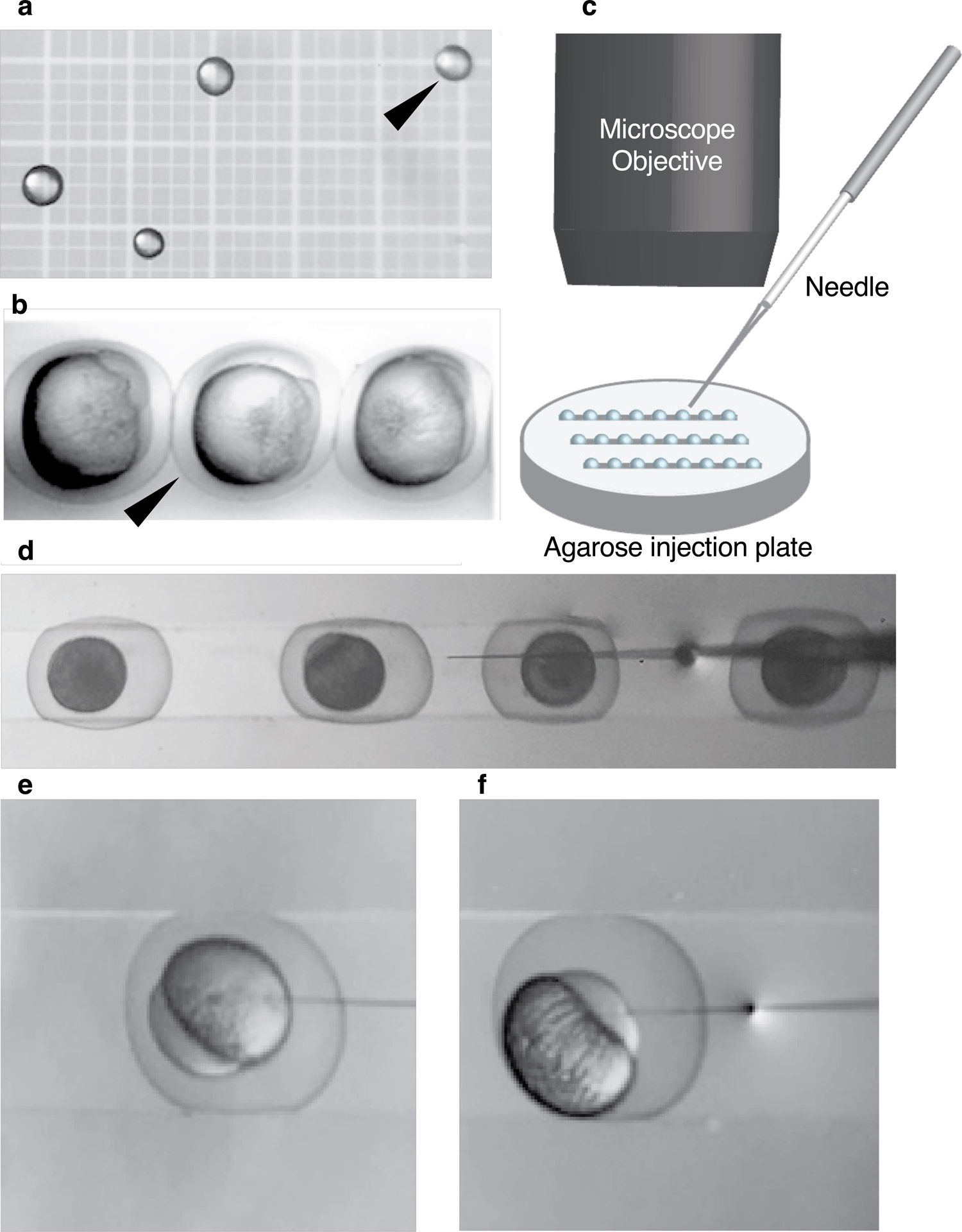
*Microinjection of zebrafish embryos*. (**a**) Calibration of injection using a hemocytometer. A cut needle was loaded with mRNA and the needle was cleared and wetted in 10% HBSS. Several injections were made into the oil overlaying the hemocytometer. The drop marked with an arrow is approximately 2 nl based on the grid of the hemocytometer. (**b**) Embryos at the two-cell stage are aligned in an injection pad. These embryos are acceptable for mRNA injection but not for plasmid injection [for single cells, see (f)]. (**c**) Schematic illustration showing a side-view of optimal set up of the embryos, microscope, and injection needle for creating zebrafish embryos expressing Halo-POI. Also see Extended Data Fig. 8. (**d**) Single-cell embryos aligned in an injection plate. These are ideal for plasmid and mRNA co-injection. The needle is above the embryos with the tip of the needle in HBSS (aiming at the yolk sac for mRNA injection). Embryos can be injected from left to right or vice versa by moving the plate to position subsequent embryos to align with the needle. (**e–f**) Injection into (**e**) the yolk sac of a single-celled embryo (mRNA injection) or (**f**) a single-cell embryo (mRNA and plasmid co-injection).

**Extended Data Figure 8.**
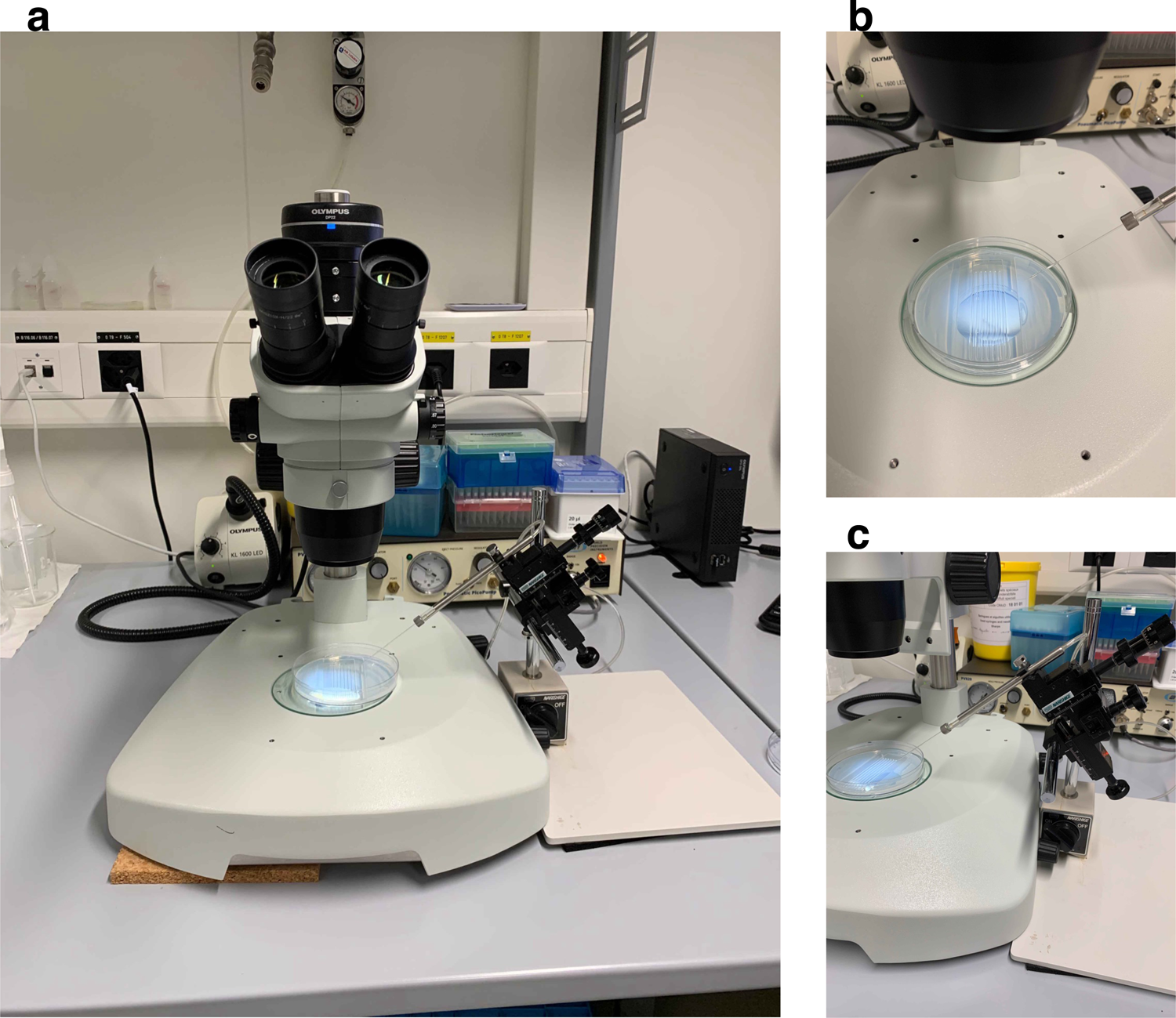
*Set up of microscope for microinjection*. One optimal setup for zebrafish embryo injection. **Note:** Whole area has been sprayed with RNaseZAP and wiped with a similarly wetted kimwipe or paper towel. (**a**) Front view, (**b**) top view, and (**c**) side view looking at injection plate and needle. Needle should be kept in HBSS once it is loaded with mRNA, to avoid clogging, and cleared at least once prior to injection.

**Extended Data Figure 9.**
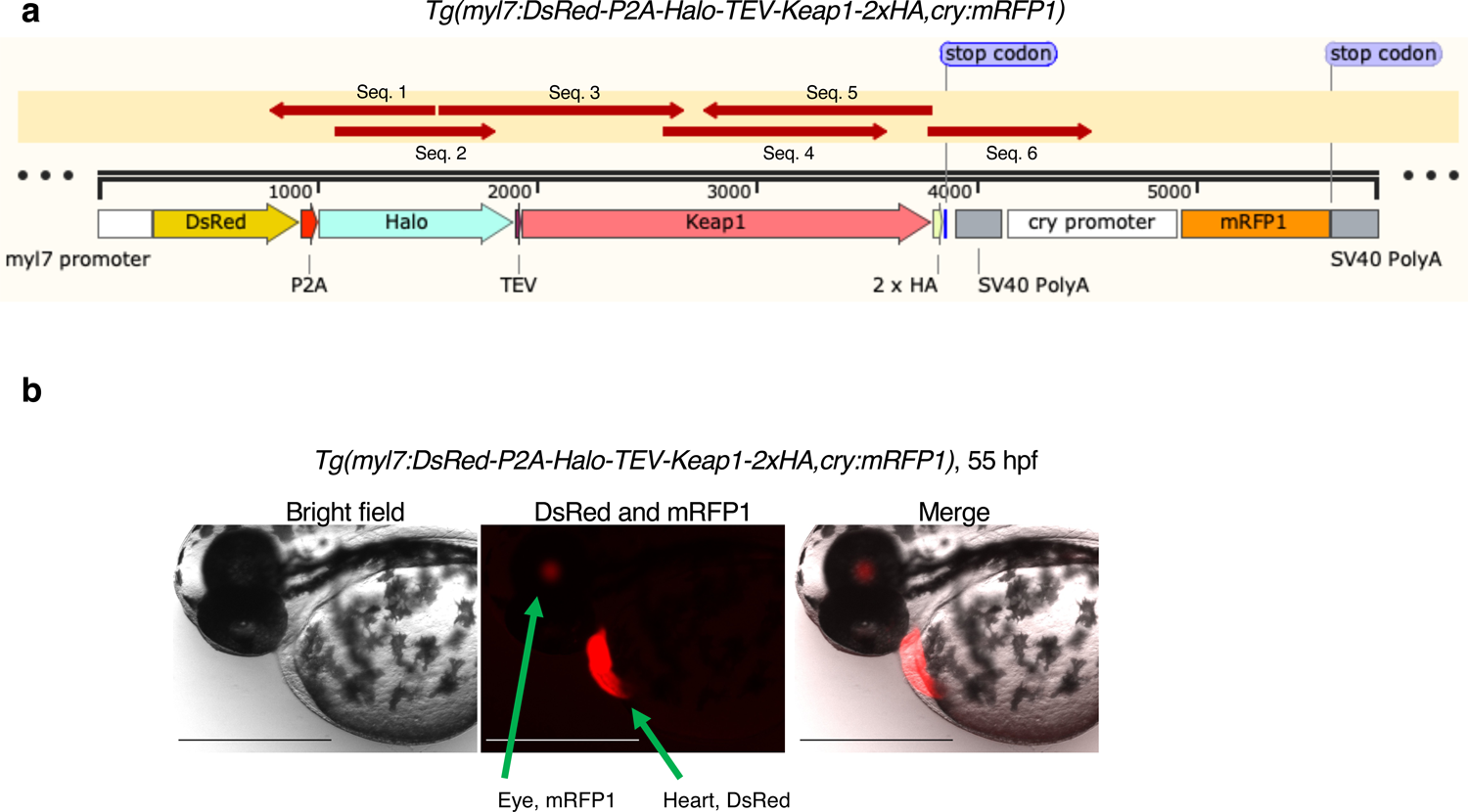
Transgenic fish line, Tg(myl7:DsRed-P2A-Halo-TEV-Keap1-2xHA,cry:mRFP1), expressing Halo-TEV-Keap1 in cardiomyocytes. (**a**) Scheme of the inserted myl7:DsRed-P2A-Halo-TEV-Keap1-2xHA-polyA sequence. P2A-Halo-TEV-Keap1-2xHA-polyA sequence was validated by six Sanger sequencing analyses: seq. 1 covers 785-1538; seq. 2 covers 1085-1811; seq. 3 covers 1560-2668; seq. 4 covers 2576-3597; seq. 5 covers 2760-3800; seq. 6 covers 3777-4525. Codon number in the scheme: *P2A*: 928-1011; *Halo*: 1012-1899; TEV protease recognition site: 1900-1932; *Keap1*: 1933-3801; 2 x HA tag: 3802-3855; left stop codon: 3856-3858; left SV40 polyA signal sequence: 3907-4113. Also see Table 3 for sequencing primers. (**b**) The *myl7:DsRed* expression in fish cardiomyocytes was visible in both live fish (55 hpf) and formaldehyde-fixed fish (see also Fig. 5, and Extended Data Fig. 4). The *cry:mRFP1* expression in fish eye lens was only seen in live fish, but not in formaldehyde-fixed fish. Scale bar = 500 µm in all images.

## Extended Data

Extended Data Figure 1–9

