## Supplementary figures and images for "Z-REX: Shepherding Reactive Electrophiles to Specific Proteins Expressed either Tissue-Specifically or Ubiquitously, and Recording the Resultant Functional Electrophile-Induced Redox Responses in Larval Fish"

### source data 1

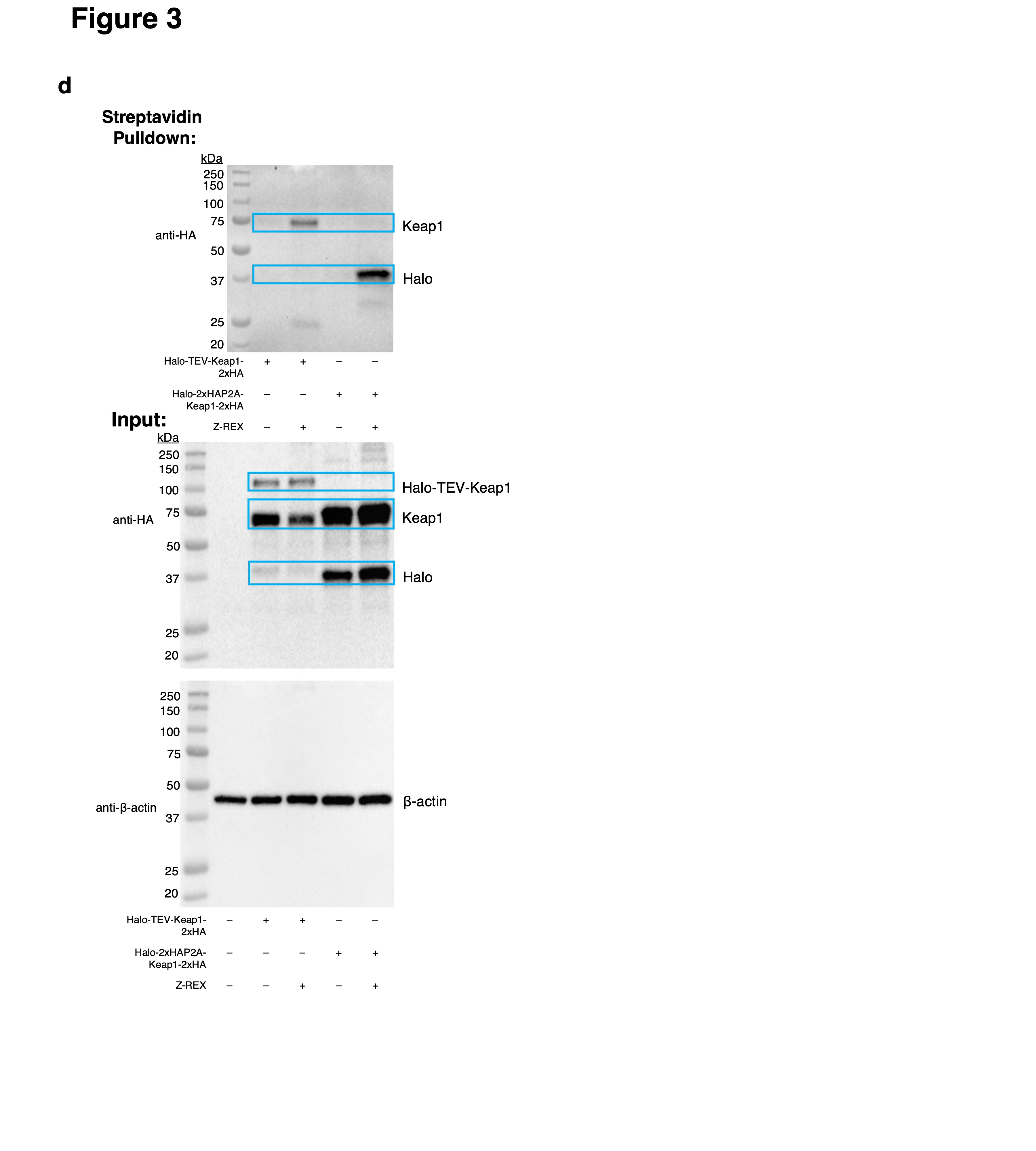
